# Origin and evolution of acrocentric chromosomes in human and great apes

**DOI:** 10.64898/2025.12.22.696095

**Authors:** Steven J. Solar, Prajna Hebbar, Leonardo Gomes de Lima, Alex Sweeten, Arang Rhie, Tamara Potapova, Luciana de Gennaro, Andrea Guarracino, Juhyun Kim, Brandon D. Pickett, Benedict Paten, Melissa A. Wilson, Sergey Koren, Erik Garrison, Evan E. Eichler, Mario Ventura, Jennifer L. Gerton, Adam M. Phillippy

## Abstract

The short arms of human acrocentric chromosomes are characterized by nucleolar organizer regions essential for ribosome biogenesis, but their highly repetitive nature has hindered genomic analysis. Leveraging the recently completed genomes of all major great ape lineages, we identified recurrent features of their acrocentrics, including enriched repeat classes, centromere repositioning by whole-arm inversion, interchromosomal sequence exchange, and birth-and-death evolution of multiple gene families under selection. Together, these processes have enabled the repeated amplification and diversification of the FRG1 gene family over 25 million years of ape evolution, and, in gorilla, the amplification of a novel IGSF3-GGT fusion gene. Similar evolutionary events also explain the distribution of segmental duplications and heterochromatin in the modern human genome, predisposing it to karyotypic abnormalities such as Robertsonian translocations. Our findings highlight acrocentric chromosomes as key drivers of evolution in the great apes, with implications for speciation, adaptation, and clinical genomics.

## Introduction

The evolutionary history of the apes is well established, with gibbons diverging before the great ape common ancestor, followed by splits of orangutan, gorilla, human, and chimp ^1^. This tree was first determined using molecular and cytogenetic techniques, including banding and fluorescence *in situ* hybridization (FISH) to identify relationships between segments of human and non-human ape chromosomes ^2–4^. It was similarly established that mammalian karyotype evolution occurs primarily through the rearrangement of syntenic blocks of DNA, in the form of chromosome fusion, fission, translocation, or inversion ^5^, and that these large-scale structural changes can facilitate adaptation and speciation ^6^. Conveniently, most human chromosomes are fully syntenic with their great ape orthologs, with the exception of a telomeric fusion of ancestral chromosomes 2A and 2B in human ^7,8^ and a balanced translocation of human orthologs 5 and 17 in gorilla ^9^. Within chromosomes, inversions are the most common large-scale structural variants between the great apes ^10,11^, but until recently, we lacked the complete genome sequences needed to fully characterize these events, especially within the short arms of the acrocentric chromosomes.

With the recent sequencing of complete, telomere-to-telomere (T2T) reference genomes for human and all other major ape lineages, it is now possible to investigate the structural evolution of ape karyotypes at the sequence level, including the 12.5–27.3% of bases within each genome that fail to align to the other apes ^11–15^. Here we analyze complete T2T genomes of the great apes: *Homo sapiens* (human, HSA), *Pan paniscus* (bonobo, PPA), *Pan troglodytes* (chimp, PTR), *Gorilla gorilla gorilla* (gorilla, GGO), *Pongo abelii* (Sumatran orangutan, PAB), and *Pongo pygmaeus* (Bornean orangutan, PPY), as well as the lesser ape *Symphalangus syndactylus* (siamang, SSY). Advances in long-read sequencing ^16,17^ and genome assembly ^18^ have unlocked previously dark regions of the genome including segmental duplications ^19^, amplified gene families ^20^, chromosomal fusions ^21^, and satellite tandem repeats ^22–24^. Such repeats are commonly enriched on the short arms of acrocentric chromosomes, and because of their tendency to recombine, are common sources of structural variation in the genome.

The short arms of ape acrocentric chromosomes frequently contain a nucleolar organizer region (NOR) comprising a tandem array of ribosomal DNA (rDNA) repeats, which contain the 45S ribosomal RNA (rRNA) genes and around which the nucleolus forms during interphase ^25,26^. Because of this relationship between acrocentric chromosomes and NORs, the two concepts are often conflated, but not all acrocentric chromosomes contain a NOR. Here we use the original cytogenetic definition of an acrocentric chromosome based solely on the asymmetric size of the arms ^27^, by which human Chromosomes 13, 14, 15, 21, 22, and Y are classified as acrocentric ^28^. We use rDNA+ and rDNA− to indicate the presence of a 45S rDNA array, inferring the presence of a NOR, but note that variable array loss or silencing can occur among individuals ^29^. Even though it is acrocentric in appearance, human ChrY is not typically referred to as an acrocentric in clinical contexts because it is consistently rDNA–. However, it does share many characteristics with the autosomal acrocentrics including satellite content ^15^, nucleolar association ^30^, and Robertsonian translocations ^31^. Additionally, ChrY is rDNA+ in both siamang and orangutan, and therefore may have been rDNA+ in the ancestor to all apes ^11^.

Short arms are generally gene-poor, heterochromatic, and enriched for segmental duplications and satellite repeats. Despite being spread across different chromosomes, the short arms of the human acrocentrics (excluding ChrY) share a windowed median sequence identity of 99% ^12^. This high similarity is a hallmark of concerted evolution ^32^, first observed within the highly homogenized sequence of rDNAs ^33^, but now understood to involve other regions of the short arms as well ^34^. Termed pseudo-homolog regions (PHRs), this high degree of interchromosomal sequence similarity is maintained by various forms of sequence exchange that homogenize the PHRs faster than they can diverge by mutation in a process known as molecular drive ^35^. Here, we use “exchange” to mean the transfer of sequence information between loci by any means, including allelic and non-allelic crossing over, gene conversion, sister chromatid exchange, template switching, break-induced replication, etc. Compared to the other chromosomes, interchromosomal exchange on the short arms may be more frequent due to their co-localization at the nucleolus and unique composition including rDNA and SST1 repeats ^36^.

Given their propensity for sequence exchange, it is not surprising that the number and identity of acrocentric chromosomes vary widely between species ^37^. For example, both orangutan species included here contain up to 10 rDNA+ acrocentric chromosomes per haplotype, including ChrY, while siamang contains as few as one ^11,38,39^. Additionally, 1 in 800 people contain a Robertsonian translocation ^21^, in which two acrocentric chromosomes are fused by their short arms. Thus, the short arms of acrocentric chromosomes have shown a propensity for structural variation that warrants a systematic investigation across the great apes.

In this study, we survey the newly complete sequence of great ape acrocentric chromosomes and revisit their evolutionary history. Our analyses uncover common mechanisms of evolution across the short arms of all great ape acrocentric chromosomes, highlighting the important role that they have played in evolution, adaptation, and potentially speciation.

## Results

### Organization of the acrocentric chromosomes

Orangutans appear closest to the ancestral hominoid karyotype, for which Stanyon *et al.* predicted 11 acrocentric chromosomes ^40^. All but two of these chromosomes (orthologs of HSA 11 and 20) remain acrocentric and rDNA+ in the orangutans. Additionally, both orangutan and siamang Y chromosomes contain active rDNA arrays. All other great ape acrocentrics are orthologous to one of the 10 orangutan acrocentrics, so we infer that the orthologs of human chromosomes 2A, 2B, 9, 13, 14, 15, 18, 21, 22, and Y were likely acrocentric and rDNA+ in the great ape common ancestor. Thus, for each species we annotate these chromosomes as rDNA+ acrocentric, rDNA− acrocentric, or ancestrally acrocentric, and refer to them in terms of their human ortholog, e.g. GGO-HSA22 refers to the gorilla ortholog of Human Chr22 (**Figure 1A**). Gibbons are unusual, having undergone at least eight chromosome fusions since the ancestral hominoid karyotype ^5,41^, and are included only as an outgroup.

**Figure 1.**
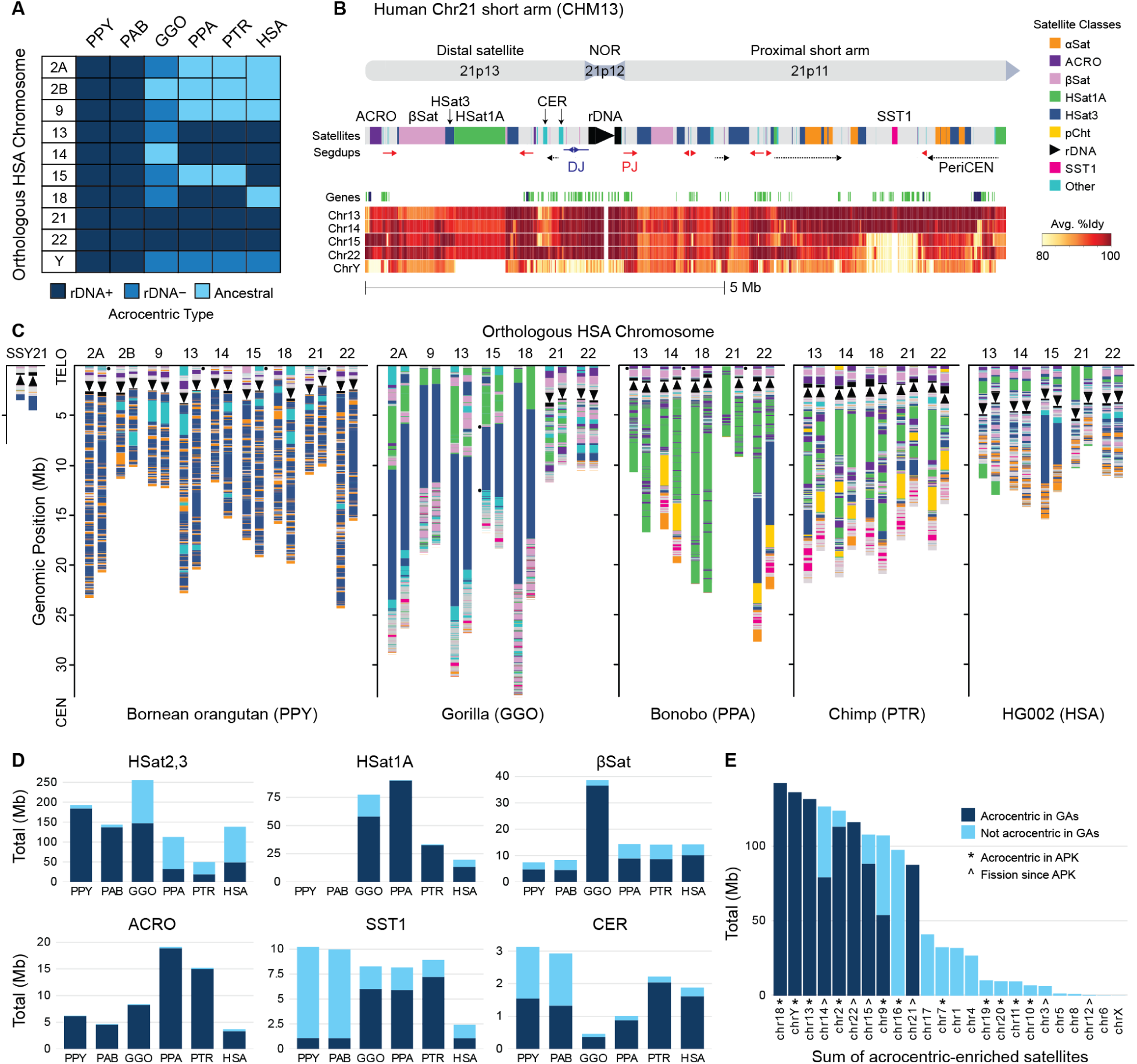
Short arms of the great ape acrocentric chromosomes. **(A)** rDNA array status of the 10 ancestral great ape acrocentric chromosomes in each species studied here. Ancestral refers to chromosomes that once were, but are no longer, acrocentric and have lost their rDNA array. **(B)** Example structure of HSA21 short arm from CHM13. Cytogenetic labels are given on top, followed by annotation of the satellites, segmental duplications (colored by homology to the DJ, PJ, or periCEN), gene predictions (green: noncoding, blue: coding), and pairwise identity heatmap. The identity heatmap shows, for each window of Chr21, the highest identity of a window in the other acrocentrics (not necessarily at the same position). The rDNA array has been artificially shortened at the visible break in the heatmap and the black triangle indicates its direction of transcription. **(C)** Short arm satellite annotations for all diploid pairs of autosomal acrocentric chromosomes from the telomere (top) up to, but not including, the centromere (bottom). Ancestral acrocentrics that have converted to metacentrics are excluded. Maternal haplotypes are on the left, except for PPY and PTR where the parent of origin is unknown and the “primary” haplotype is on the left as assigned by Yoo *et al.* ^11^. Small dots indicate missing telomeres or remaining gaps in the assemblies. **(D)** Total bases in each species for six different satellite classes, partitioned by bases on acrocentric chromosomes (dark blue) or non-acrocentrics (light blue). **(E)** Total bases for the same six acrocentric-associated satellites summed over each homologous human chromosome in the great apes (GA) and stacked by that chromosome’s acrocentric status in each species. Ancestral primate karyotype (APK) predictions are from ^40^.

The typical short arm of a human autosomal acrocentric chromosome is arranged around the rDNA array with a smaller distal region towards the telomere and a larger proximal region towards the centromere (**Figure 1B**). We replicated Figure 4 from Nurk *et al.* ^12^ using updated methods, and including ChrY ^13^. These self and pairwise alignments of T2T-CHM13 reveal three general classes of segmentally duplicated sequence on the human acrocentric short arms: distal junction (DJ), proximal junction (PJ), and pericentromeric sequence. The previously described DJ is a ∼400 kb conserved region upstream of the rDNA array that typically appears as a single copy per chromosome and contains a >200 kb palindrome that encodes a long noncoding RNA associated with nucleolar organization ^42^. In contrast, the PJ and pericentromeric sequences are variable in copy number, with the PJ widely duplicated on both sides of the rDNA array, and the pericentromeric duplications localized near the centromere. Inter-chromosomal similarity varies across the short arm and is highest in and around the rDNA arrays and the multi-megabase pericentromeric duplication on chromosomes 13, 14, and 21 that has been associated with Robertsonian translocations ^21^. These regions contain mostly non-coding or pseudogene annotations, but also the protein-coding gene *TPTE* (Transmembrane Phosphatase with TEnsin homology) in the pericentromeric regions of Chr13 and Chr21 ^43^. Compared to the rest of the genome, the human short arms are also strongly enriched for satellite repeats. Using T2T-HG002 as an example diploid human genome ^14^, we annotated 76% of bases on the short arms as satellite repeats, compared to 7% for the rest of the autosomes, including human satellite 1 (HSat1A and HSat1B), human satellite 3 (HSat3), beta satellite (βSat), alpha satellite (αSat), Walu, ACRO, and SST1. Note that human satellites 2 and 3 both derive from the same pentameric sequence (GGAAT), and we will refer to them jointly as HSat3 when not making a human-specific distinction ^44^.

The other great ape autosomal acrocentrics follow a similar pattern of interspersed segmental duplications and an enrichment of satellites. Similar to human, their short arms average 87% satellite. However, unlike the rest of the autosomes, which tend to be highly syntenic, the short arms vary widely in content and organization (**Figure 1C**). Like human, all rDNA arrays are uniformly oriented within a species and located towards the distal end of the chromosome, positioned an average of 1–3 Mb from the telomere and 7.5–20 Mb from the centromere. In human, gorilla, and orangutan, rDNA transcription proceeds in the direction of the centromere, while in chimp and bonobo it is towards the telomere ^42^. These inversions in chimp and bonobo include the DJ, keeping it always upstream of rDNA transcription and implying a functional link with the rDNA array.

The orangutan proximal short arms stand out for their lack of HSat1A (0% of bases), unique tandem arrays of L1PA3 retrotransposons, and domination by HSat3 (49% PAB, 59% PPY) and monomeric αSat satellites (19% PAB, 16% PPY) (Supplementary Figs. S1–S2). In contrast, human, gorilla, chimp, and bonobo show a more varied composition of structures and satellites, including an invasion of HSat1A, increased enrichment of SST1 and βSat, and multi-megabase pericentromeric segmental duplications (Supplementary Figs. S3–S6). These duplications are conspicuously amplified on the rDNA− gorilla acrocentrics (Supplementary Fig. S6). Siamang’s sole rDNA+ autosome is comparatively minuscule, containing little sequence other than its large 4.5–10 Mb rDNA array and smaller HSat3 and βSat arrays (Supplementary Fig. S7).

Human satellite repeats are often referred to as “centromeric satellites” due to their association with human centromeres ^22^. However, aside from centromeric αSat, this association is just as strong with the short arms of the acrocentric chromosomes (**Figure 1D**). In particular, 80% of all HSat3, HSat1A, βSat, ACRO, SST1, and CER bases are specific to acrocentric chromosomes in at least one great ape species (Supplementary Table S1;Tab1). Furthermore, the vast majority of these satellites found elsewhere in the genome are on chromosomes that we predict were acrocentric in the great ape ancestor, suggesting they are relics from their acrocentric origin (**Figure 1E**). Most other non-acrocentric chromosomes with an enrichment of acrocentric satellites in their pericentromere are either predicted to have been acrocentric in the ancestral primate karyotype or derived from a fission event ^5,40^. For example, HSA16 is predicted to have been acrocentric in the ancestral primate karyotype, and even though none of the great ape orthologs are acrocentric, they all contain both HSat3 and 45S rRNA pseudogenes in their pericentromeres (Supplementary Fig. S8). Similarly, HSA20 is predicted to have been acrocentric in the ancestral hominoid karyotype, and all modern orthologs retain acrocentric-like features in their pericentromeres, especially for human, chimp, and gorilla. Notable exceptions without a known acrocentric ancestor in the primates are HSA1 and GGO-HSA4, which contain species-specific satellite expansions in their pericentromeres, and the HSA17 orthologs, which contain a balance of acrocentric satellites in all great apes (Supplementary Fig. S9).

Tallying satellites by which chromosome they appear on reveals ACRO as the most strongly and universally enriched repeat class on the great ape acrocentrics, rivaling the specificity of rDNA itself (**Figure 1D**). ACRO is a 6 kb composite repeat, comprising ACRO1, L1PA10, MER21B, and L1MB8 Dfam annotations ^45^ that is primate-specific ^46^ and enriched for nucleolar-associated H1.0 histones ^47^. Unlike the other satellite classes, ACRO appears almost exclusively on acrocentric chromosomes (>90% of ACRO bases, Supplementary Table S1;Tab2), typically arranged as highly homogenized arrays up to a few hundred kb in length and consistently found on the distal ends of the rDNA+ short arms (**Figure 1C**). Intriguingly, ACRO1 shows evidence of expression in early embryogenesis, peaking at the 4-cell stage ^48^, but its absence from prior reference genomes has limited analysis.

Less exclusive to the acrocentrics, but also notable, is the ∼2 kb primate-specific macrosatellite SST1 ^49^ (also known as NBL2), which displays unusual methylation patterns in certain cancers^50^ and has been implicated as a common breakpoint of human Robertsonian translocations ^21^. SST1 repeats on the short arms of CHM13 are also highly methylated ^12^, actively transcribed ^23^, and enriched for PRDM9 binding motifs ^34^, indicating they may be more accessible and prone to exchange than the surrounding heterochromatin. Mirroring the invasion of HSat1A, the enrichment of SST1 on the acrocentrics rises from 11% in the orangutans to 73% in gorilla, 81% in chimp, and 45% in human (Supplementary Table S1;Tab2, **Figure 1D**). Subfamily SF1 is exclusively found on the great ape acrocentrics (or ancestrally acrocentric chromosomes), except for orangutans. This SST1 subfamily acts as a hotspot for non-allelic exchange between acrocentric chromosomes, leading to the formation of Robertsonian translocations in humans ^21^. Its origin, expansion, and restricted distribution suggests that this subfamily may be associated with structural variation events shaping acrocentric evolution in the African great apes, and if so, its absence from orangutan may explain the relative stability and uniformity of the orangutan acrocentrics (**Figure 1C**, Supplementary Figs. S1–S2).

### Karyotype evolution through short-arm inversion and rDNA loss

Orangutans have maintained the 10 acrocentrics of the inferred great ape common ancestor, but all other species show a gradual loss of acrocentrics with time. Only the orthologs of HSA 21 and 22 remain rDNA+ in all the great apes. We sought to understand the evolutionary events leading to rDNA loss and uncovered two primary causes: short-arm inversion and short-arm truncation.

Prior cytogenetic studies have noted lineage-specific pericentric inversions leading to centromere repositioning in the great ape acrocentrics ^37^, but lacked the necessary resolution to fully characterize these events. Pairwise sequence alignments of the complete great ape chromosomes revealed at least six large-scale inversion events that have converted acrocentric to metacentric chromosomes. Assuming parsimony, these events can be placed on specific branches of the phylogeny (**Figure 2A**). Beginning with the oldest, the short arm of the HSA2B ortholog was inverted in the human/gorilla common ancestor, followed by inversions of both HSA2A and 9 orthologs in the human/chimp ancestor. Additional lineage-specific inversions have since occurred on the human branch (HSA18), the chimp/bonobo branch (PTR-HSA15 and PPA-HSA15), and the gorilla branch (GGO-HSA14). It is unclear whether the original inversions included the rDNA array, but inverted or proximal rDNAs appear to be selected against in the great apes given their uniform direction and location in each species. Thus, since the orthologs of HSA 2A and 2B were both inverted prior to the human branch, they were likely rDNA− at the time, supporting the hypothesis of a telomeric or subtelomeric fusion rather than a Robertsonian translocation ^7,51^.

**Figure 2.**
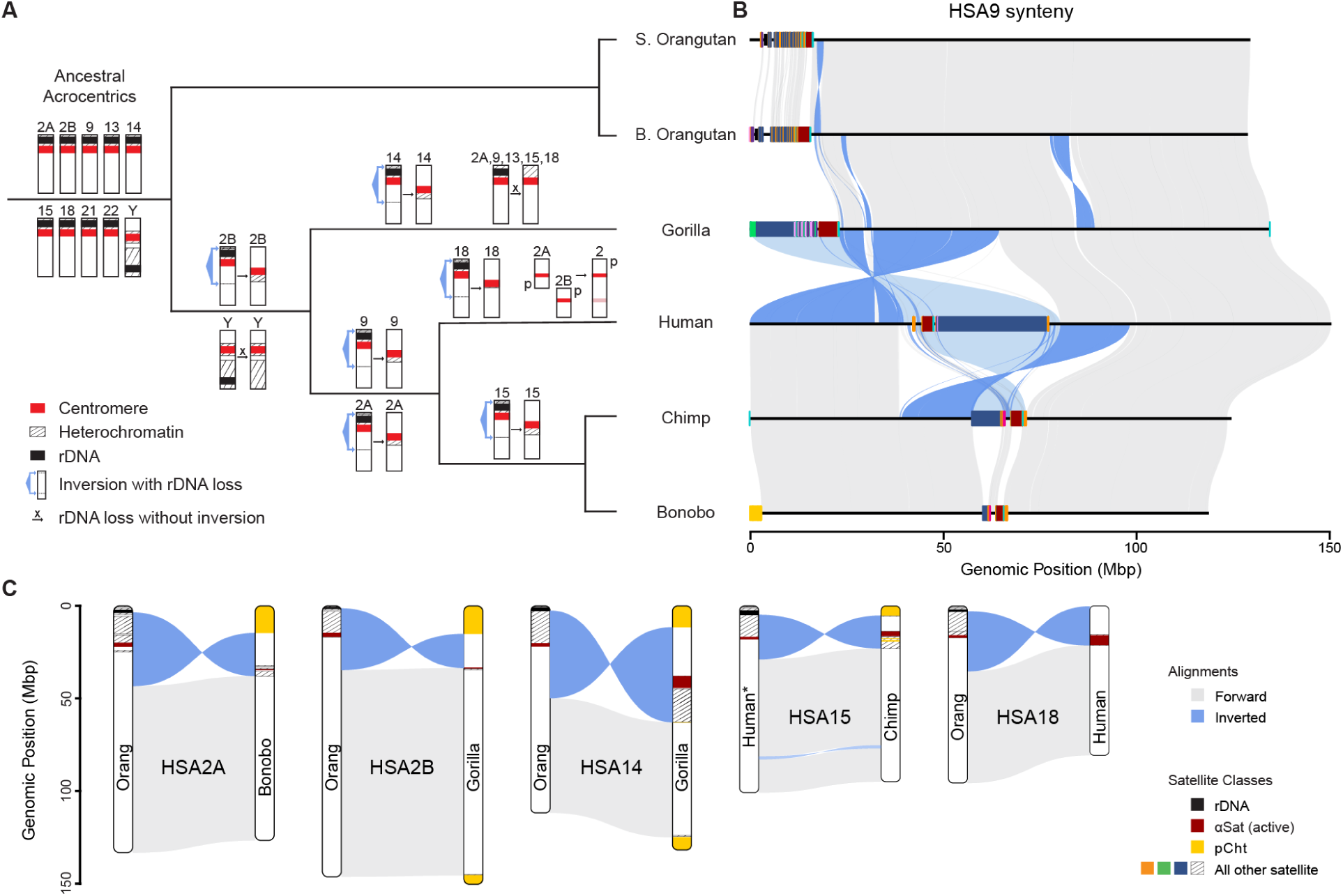
Inversion, fusion, and rDNA array loss events in the evolution of great apes. **(A)** Great ape phylogeny overlaid with the most parsimonious assignment of karyotype changes labeled on specific branches (whole chromosomes shown). **(B)** An example alignment of HSA9 orthologs across the six species studied here, showing acrocentric chromosomes in both orangutans and gorilla; conversion of acrocentric to metacentric in human, chimp, and bonobo; and a pericentric inversion between human and chimp/bonobo. The lighter blue highlights indicate the inferred repositioning of satellites that are too divergent to be reliably aligned, e.g. the repositioning of a large HSat3 block relative to the αSat centromere. **(C)** Simplified alignment schematics showing the five inversion events against an orangutan reference, with the exception of HSA15 shown against the human reference due to other complicating paracentric inversions relative to orangutan. pCht is a subterminal satellite cap that arose independently in the gorilla and Pan lineages ^52^. Complete alignments for all events are given in Supplementary Figs. S10–S19.

Each short-arm inversion was mediated by a break in the long arm of the ancestral chromosome, followed by a whole-arm inversion that repositioned the centromere and short-arm heterochromatin toward the middle of the chromosome. In the case of HSA9, the heterochromatin of the ancestral short arm was flipped into the middle of the chromosome, including a large HSat3 block and other acrocentric-specific repeats such as SST1 (**Figure 2B**). Although the satellite sequences have rapidly diverged due to sequence turnover and do not easily align, the location of the HSat3 block relative to the αSat centromere has swapped from gorilla to human, consistent with the inversion. Directly aligning chimp to gorilla, rather than through human, would show two adjacent inversions and a centromere repositioning, but this structure is better described as a single whole-arm inversion in the common ancestor of human and chimp followed by a pericentric inversion in chimp. Likely as a result of the retained acrocentric-like satellites and duplications, the centromere of HSA9 is also polymorphic in human, with heteromorphisms such as 9qh+ and inv(9)(p11q13) present in more than 1% of the population ^53^ and associated with infertility ^54^.

This pattern of acrocentric to metacentric conversion by short-arm inversion is consistent across the other chromosomes, with the long-arm breakpoints localized within euchromatin approximately 10–40 Mb from the modern orangutan centromeres (**Figure 2C**, Supplementary Figs. S10–S19). Given the high rate of turnover, the short-arm breakpoints are more difficult to pinpoint, and the amount of acrocentric-associated sequence retained in the pericentromere varies considerably by chromosome.

In addition to short-arm inversions, there have been multiple in-place rDNA array losses that have fixed, including from chromosome Y on the human/gorilla/chimp branch, and from GGO-HSA 2A, 9, 13, 15, and 18 on the gorilla branch. Transient rDNA loss appears relatively common in the great apes, given that three heterozygous rDNA array losses were observed in these six complete genomes alone ^11^, including PAB-HSA13 in Sumatran orangutan (near-complete short arm loss), PPY-HSA18 in Bornean orangutan (distal truncation), and PPA-HSA21 in bonobo (distal truncation) (**Figure 1C**). Thus, it is possible that short arm truncation or rDNA loss could have preceded the inversions, which would explain the smaller blocks of heterochromatin found in the inverted GGO-HSA2B and HSA18. Once the acrocentric was converted into a metacentric, the rDNA array could not be easily regained via sequence exchange with the remaining acrocentrics, and the loss would become fixed.

The mechanism of rDNA array loss on ChrY is more difficult to ascertain, given its high structural variability and rapid evolution among the great apes ^13,15,55^. However, like the autosomal arrays, the ChrY rDNA arrays in orangutan and siamang also appear on the end of the chromosome, allowing for loss by truncation without a substantial loss of ChrY sequence. Although no ChrY rDNA arrays remain in the other great apes, remnants of rRNA pseudogenes can be found near the ChrY centromere in human (Supplementary Fig. S20), and the DJ palindrome has undergone remarkable amplification on the Y chromosomes of chimp and bonobo along with the Y-specific genes *TSPY* and *RBMY* (Supplementary Fig. S21).

### Pseudo-homolog regions on the short arms

The complete CHM13 human genome revealed high similarity between all pairs of acrocentric chromosomes ^12^, which was later shown to be present in the population, suggesting ongoing sequence exchange ^34^. These so-called PHRs maintain high sequence identity between heterologous chromosomes, which can be used as a surrogate for detecting exchange in the absence of population-scale data.

Using a 99% pairwise alignment identity threshold over a 10 kb span, we found evidence of PHRs across the short arms of all great ape acrocentrics (**Figure 3A**). Considering all sequence classes, there is an average of ∼186.5 Mb of identified PHRs per haplotype in each species, and the largest sum of PHRs between a single pair of chromosomes is 9.8 Mb (PPY-HSA2A and PPY-HSA22). The rDNA arrays are the most highly conserved PHRs among all species, including the rDNA array and DJ sequence, which share >99% identity across the apes compared to 97–99% for the entire short arm. Aside from the rDNAs, most PHR bases (115.5 Mb of the total 186.5 Mb) are dominated by satellites, such as the similar short arms of the orangutans, and the subterminal satellite regions of chimp, bonobo, gorilla, and siamang. After filtering rDNAs, satellites, and subtelomeric regions, ∼40.6 Mb of segmentally duplicated PHR sequence remains and distinct patterns of exchange between certain acrocentrics become clear (**Figure 3A**).

**Figure 3.**
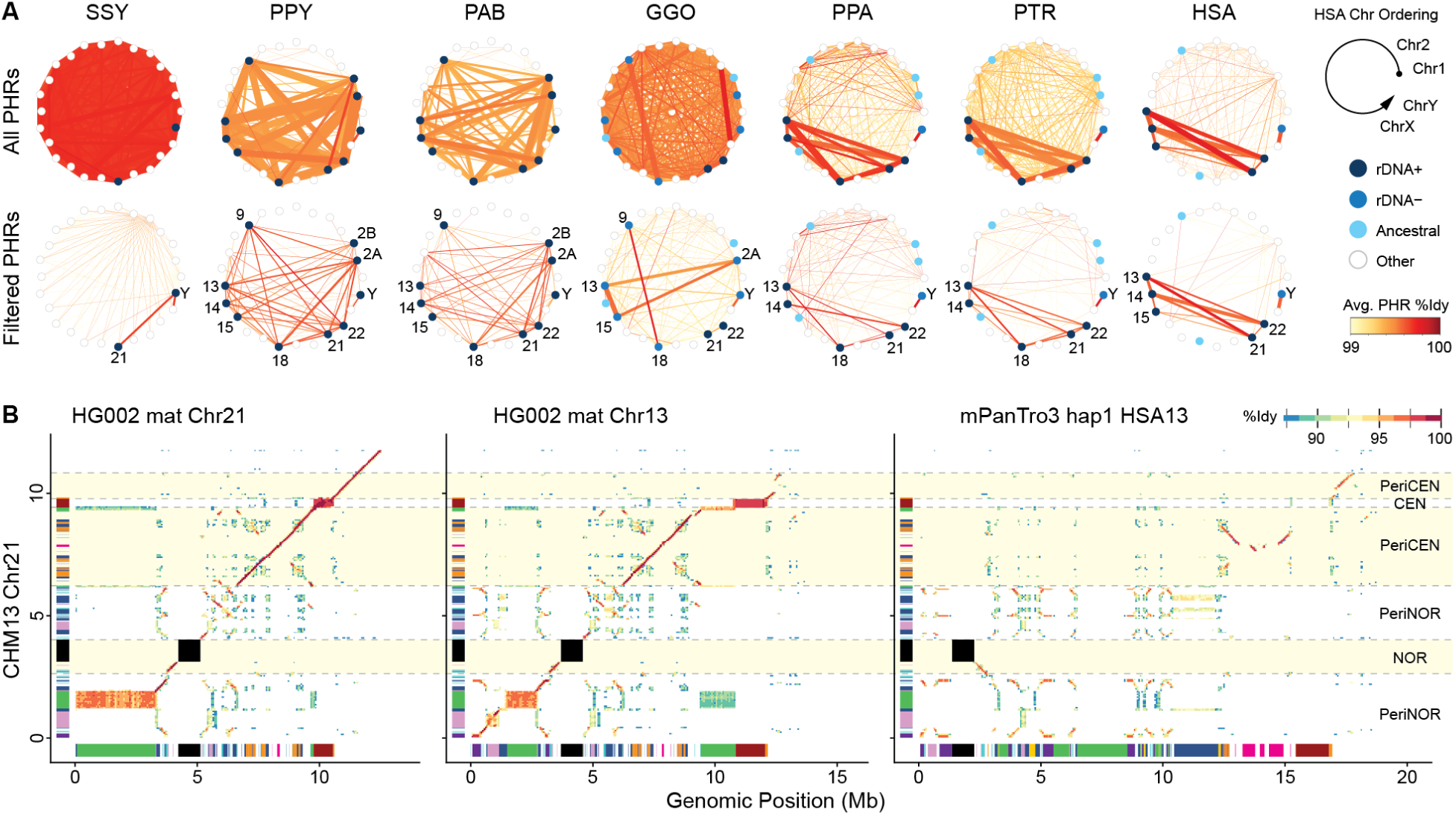
Substrates for sequence exchange between heterologous acrocentrics. **(A)** Putative PHRs identified by sequence similarity between pairs of chromosomes, where the thickness of the line indicates the number of shared bases (max 15 Mb) and color indicates the average identity (99–100%). Values are the average of the diploid pairs. Top row includes all identified similarities. Bottom row excludes rDNAs, satellites, and subterminal regions, leaving only segmentally duplicated sequences. The acrocentric chromosomes for each species are numbered. HSA is represented by HG002. **(B)** Percent identity dotplots between different human chromosomes illustrate a large, high-identity PHR between HSA 13 and 21 with residual similarity to PTR-HSA13. The corresponding satellite annotations from Figure 1 are displayed on the X and Y axes. The centromere, rDNA array, and their surrounding regions are highlighted with respect to CHM13 Chr21.

Certain pairs of chromosomes show stronger PHR relationships, such as Chr13 and Chr21 in human HG002, which share an average of 4.6 Mb per haplotype at 99.7% identity. In fact, maternal Chr13 is almost entirely co-linear with maternal Chr21 from the centromere to the rDNA array. Alignment of these chromosomes in HG002 to CHM13 Chr21 shows a high degree of similarity extending nearly 3 Mb into the short-arm pericentromeric region of all three chromosomes (**Figure 3B**), which is the PHR involved with Robertsonian formation, as noted above ^21^. The distal satellite region (distal PeriNOR) of CHM13 Chr21 is also a better match to HG002 Chr13 than Chr21, reflecting a past interchange. Indeed, ectopic exchange between the PHRs of HSA 13 and 21 was recently observed in a multi-generation pedigree ^56^.

Given the rapid evolution and high degree of turnover on the short arms, the sequence and location of PHRs are not well conserved between species and synteny quickly breaks down between species (Supplementary Figs. S10–S19). As noted previously, the rDNA array and DJ are inverted in chimp relative to human and surrounded by similar (but shuffled) classes of satellites and segmentally duplicated PJ sequences. In both species, the patterns of segmental duplication appear primarily organized around the rDNA array and centromere, with some local similarity remaining between species (**Figure 3B**). For example, >100 kb fragments of the pericentromeric PHR on HSA 13 and 21 can also be found on PTR-HSA13 and GGO-HSA13 with a gap-excluded nucleotide identity as high as 98% (similar to the genome-wide average).

All great apes show similar, segmentally duplicated PHRs between all pairs of rDNA+ acrocentric chromosomes. This includes the distal satellites, which tend to have a similar structure within each species (**Figure 1C**). ChrX and ChrY also show exchange due to their pseudoautosomal regions (PARs), but the total amount of PAR sequence is often less than the acrocentric PHRs (Supplementary Table S1;Tab3). Excluding subterminal satellites, the acrocentrics that have converted to metacentrics tend to lose their PHRs, indicating a cessation of sequence exchange (**Figure 3A**). Surprisingly, the gorilla rDNA− acrocentrics contain even more PHR sequence than the rDNA+ acrocentrics, suggesting that sequence exchange may be a general feature of short arms. In gorilla, three distinct acrocentric groups emerge from the PHR analysis: GGO-HSA2A/13/15 (rDNA−), GGO-HSA9/18 (rDNA−), and GGO-HSA21/22 (rDNA+).

### Sequence exchange between the short arms of gorilla acrocentrics

To further understand the patterns of sequence exchange between the gorilla PHRs, we focused on the SST1 macrosatellite that has been previously implicated as a translocation hotspot on the human acrocentrics ^21^ and centromeric rearrangements in gibbons ^57^. It is also a convenient marker of interchromosomal exchange because its arrays are composed of well-defined ∼1.4 kb monomers that are found on all gorilla acrocentrics. Both the GGO-HSA2A/13/15 and GGO-HSA9/18 PHRs contain prominent SST1 arrays. The phylogeny of all SST1 monomers in the gorilla genome mirrors the observed PHR associations, with GGO-HSA2A/13/15 and GGO-HSA9/18 co-clustering (**Figure 4A**), indicating that SST1 is exchanged between heterologous short arms, similar to human.

**Figure 4.**
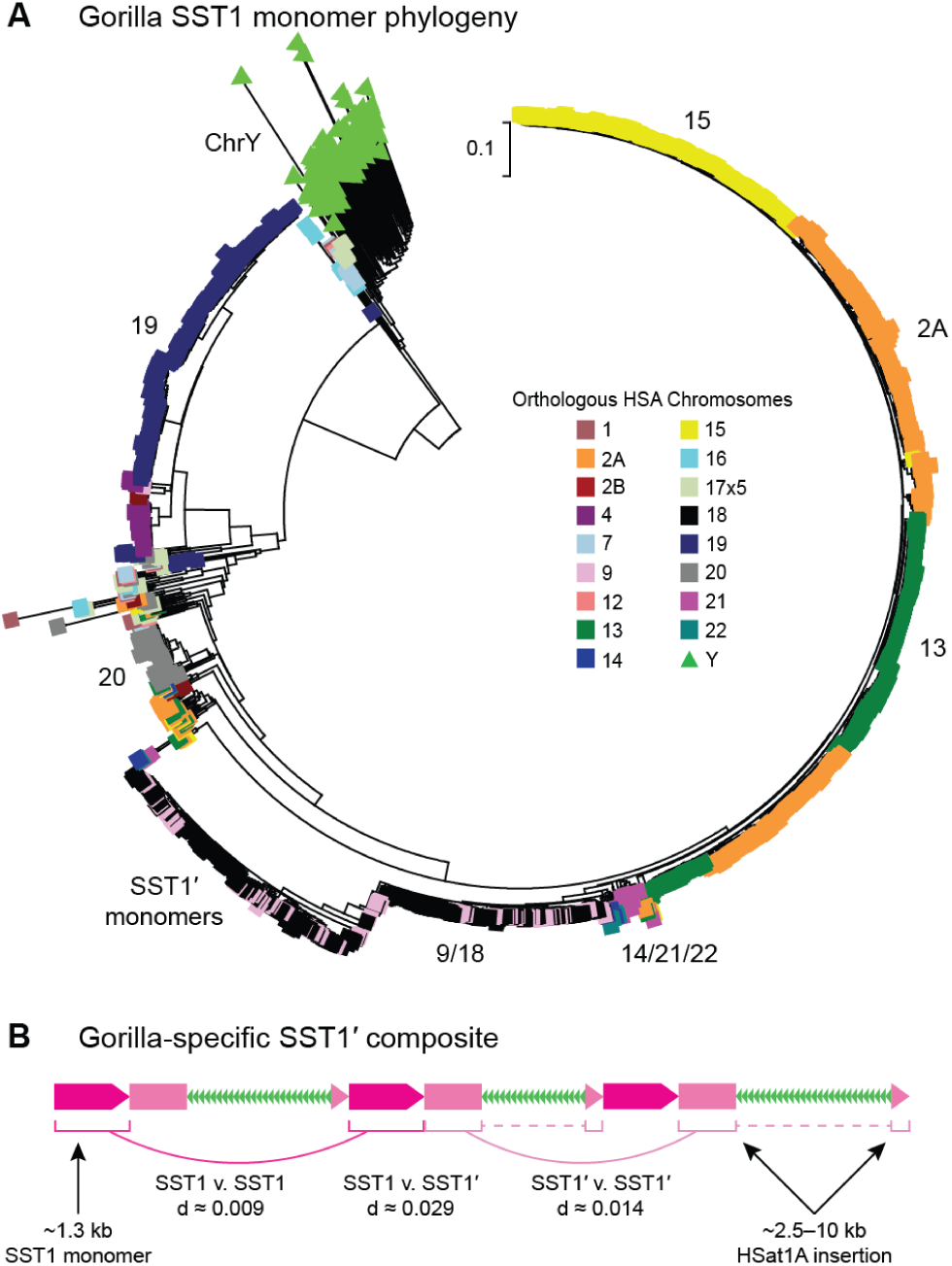
SST1 exchange between the gorilla acrocentrics. **(A)** Phylogenetic relationship of 3,078 full-length SST1 monomers in gorilla with color indicating the source chromosome. All SST1 copies on the acrocentric chromosomes are separated from all other chromosomes by a single branch, with the exception of GGO-HSA14, which was only recently converted to a metacentric. SST1 repeats from GGO-HSA9 (pink) and GGO-HSA18 (black) co-cluster, consistent with frequent recombinational exchange. A subclade of SST1′ monomers exclusive to GGO-HSA9/18 shows accelerated evolution. **(B)** Schematic representation of the SST1 composite found on gorilla chromosomes GGO-HSA9/18, where the arrays are organized as tandemly arranged dimers of SST1 (magenta) and SST1′ (pink) with HSat1A insertions (green). Average nucleotide distances are given between SST1 and SST1′ copies showing the increased divergence of SST1′ copies. Differential clustering of SST1 and SST1′ monomers in the phylogeny suggests the SST1′ copies are evolving as dimers.

In particular, the GGO-HSA9/18 monomers show the highest similarity between chromosomes and co-cluster within the same clade, confirming recent exchange between these chromosomes. Closer inspection of these monomers revealed a unique composite (SST1′) of SST1 and HSat1A elements that is only found within the PHRs of these two chromosomes (**Figure 4B**). Excluding the HSat1A insertions, the monomeric SST1′ sequence itself shows an elevated substitution rate compared to SST1, indicative of accelerated evolution within the subclade. The GGO-HSA21 SST1 monomers are also highly similar to GGO-HSA9/18 monomers, but branch ancestrally in the tree. Similarly, SST1 monomers on ancient hominoid acrocentrics, such as GGO-HSA20, cluster ancestrally to the copies on modern acrocentrics, indicating a cessation of exchange after the chromosome converted from acrocentric to metacentric.

Interchromosomal exchange between the gorilla rDNA− acrocentrics shows that PHRs are not unique to rDNA+ chromosomes and may be a general feature of acrocentric chromosomes. The three groups of acrocentric chromosomes in gorilla each have their own unique satellite organization that is consistent between haplotypes, with minor variation in array sizes (**Figure 1C**). For example, GGO-HSA9/18 and GGO-HSA2A/13/15 share a large HSat3 array, but this array is oppositely oriented between the two groups, while GGO-HSA21/22 share an rDNA array but carry very little HSat3. GGO-HSA9/18 and GGO-HSA2A/13/15 are very self-similar within groups, as are the distal ends of GGO-HSA21/22. These large blocks of homologous sequence between chromosome groups, including both satellites and segmental duplications, provide a substrate for sequence exchange between PHRs by mechanisms of non-allelic homologous recombination (NAHR). Notably, we did not observe the rDNA− acrocentrics spatially organizing with the rDNA+ chromosomes during mitotic interphase (Supplementary Fig. S22), limiting opportunities for exchange and potentially leading to the distinctive structures of the rDNA+ and rDNA− acrocentrics.

### Amplified protein-coding genes on the short arms of gorilla acrocentrics

Intrigued by the evidence of exchange between the gorilla rDNA− acrocentrics, we undertook a re-annotation of the gorilla short arms in the “primary” reference haplotype. Unlike for the other apes, this revealed 11 predicted protein-coding gene families on the gorilla short arms (**Figure 5A**). Some of these gorilla-specific amplifications have been previously described by array-based comparative genomic hybridization ^58^, FISH ^59^, and short-read sequencing ^60^, but the complete gorilla genome allowed us to determine their precise location, copy number, sequence, and functional potential of the involved genes (Supplementary Table S2). The newly annotated genes can be roughly organized into three segmentally duplicated gene clusters whose locations correspond with the similarly organized satellite structures and PHR relationships (**Figure 5B**).

**Figure 5.**
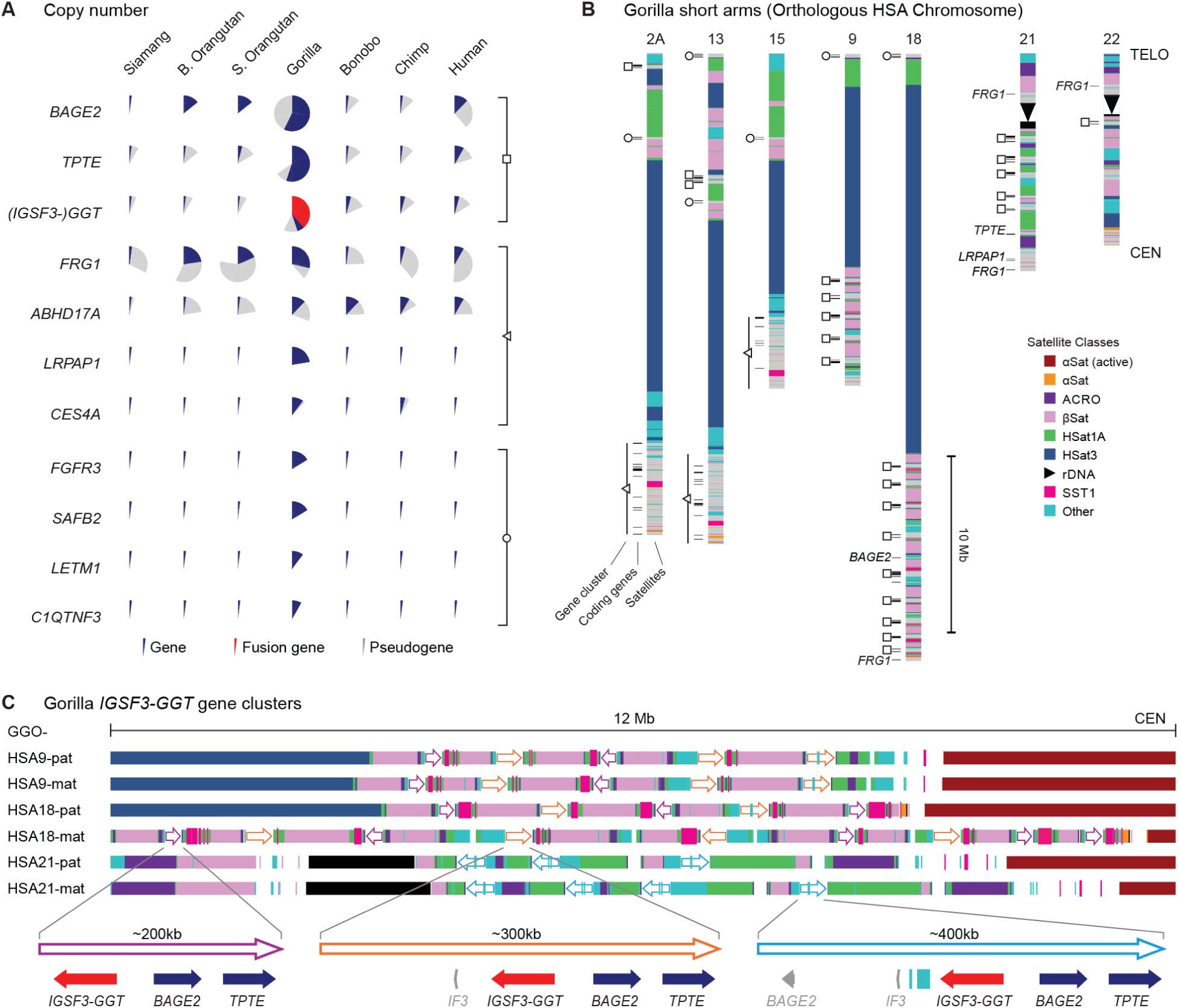
Gene amplification on the rDNA− gorilla acrocentrics. **(A)** Copy numbers of protein-coding genes (blue), fusion coding (red), and pseudogenes (gray) that have been amplified across the gorilla acrocentrics (primary haplotype). Three recurring segmentally duplicated gene clusters are indicated by a square, triangle, and circle. **(B)** Satellite and protein-coding gene annotations for the short arms of seven gorilla acrocentrics (primary haplotype, excluding ChrY), with recurring gene clusters summarized by the symbols in panel A and singleton genes annotated by name. **(C)** Pericentromeric satellites and segmental duplications on the short arm of chromosomes GGO-HSA 9, 18, and 21, illustrating amplification of the *IGSF3-GGT* gene cluster. The large blocks of “Other” satellites are mostly composites of mariner-like repeats. Hollow arrows represent the segmentally duplicated gene clusters with annotated genes below (colored by status as in panel A).

The largest amplification primarily involves GGO-HSA9/18, which share 18.1 Mb of PHR sequence at 99.6% identity (7.6 Mb after satellite filtering), including the gorilla-specific amplification of a novel fusion gene between *IGSF3* (Immunoglobulin Superfamily Member 3) and a member of the *GGT* (Gamma-Glutamyltransferase) family, along with fragments of *TPTE* (268 AA) and *BAGE2* (B Melanoma Antigen 2; 129 AA). GGT enzymes participate in glutathione metabolism and detoxification, potentially conferring an adaptive advantage for a diet rich in plant secondary metabolites ^58^. The primary gorilla haplotype contains 21 *IGSF3-GGT* fusion genes that are predicted to be coding (average length 1,335 AA). We confirmed the presence of this *GGT*-linked amplification on GGO-HSA9/18 by FISH in nine different gorilla individuals, along with weak signal on the other acrocentrics (including ChrY), suggesting this amplification is a conserved feature of gorilla genomes (Supplementary Figs. S23–25, Supplementary Table S3).

A second gene cluster occupies the pericentromeres of GGO-HSA2A/13/15, totaling 57.5 Mb of shared PHR sequence at 99.5% identity (41 Mb after satellite filtering). This cluster harbors multiple predicted protein-coding genes including *FRG1* (FSHD Region Gene 1), which is critical for muscular and vascular development ^61^, as well as *ABHD17A* (Abhydrolase Domain Containing 17A, Depalmitoylase), *LRPAP1* (LDL Receptor Related Protein Associated Protein 1), and *CES4A* (Carboxylesterase 4A). Separate from the larger cluster, individual *LRPAP1* and *FRG1* genes deriving from ancestral copies on the HSA4 ortholog are found in the pericentromere of GGO-HSA21, pointing to an initial duplication onto the acrocentrics preceding amplification. We also confirmed the presence of this *LRPAP1*-linked amplification on GGO-HSA2A/13/15 by FISH in four different gorilla individuals, with variable signal between haplotypes and again including weak signal on ChrY (Supplementary Fig. S26–S27).

Lastly, a small, amplified cluster of three genes is found towards the distal end of all gorilla rDNA− acrocentrics. The cluster always includes *SAFB2* (Scaffold Attachment Factor B2) and *FGFR3* (Fibroblast Growth Factor Receptor 3), with the third gene alternating between *C1QTNF3* (C1q And TNF Related 3) or *LETM1* (Leucine Zipper And EF-Hand Containing Transmembrane Protein 1). *SAFB2* has been associated with androgen receptor activity and male reproduction ^62^. *FGFR3* is a fibroblast growth factor receptor involved in skeletal development, with mutations in this gene known to cause achondroplasia in humans ^63^. *C1QTNF3* occurs exclusively within the clusters on the very distal ends of GGO-HSA13 and GGO-HSA9/18, while *LETM1* is restricted to spacer sequence between HSat1A and βSat on GGO-HSA2A/13/15.

Of these amplifications, the gene cluster including *IGSF3-GGT*, *BAGE2*, and *TPTE* reaches the highest copy number, with its arrays stretching more than 10 Mb in some cases (**Figure 5C**). These arrays are predominantly found on GGO-HSA 9, 18, and 21, consisting of alternating copies of the gene cluster, sometimes inverted, separated by blocks of acrocentric-associated satellites including βSat, HSat1A, ACRO, SST1, and a composite of mariner-like repeats. The array on GGO-HSA21 shows a distinct satellite pattern dominated by HSat1A and a longer version of the gene cluster, while the arrays on GGO-HSA9 and 18 are highly similar in terms of sequence and structure, indicative of more recent interchromosomal exchange.

Gorilla’s large number of rDNA− acrocentrics and their unique patterns of exchange go hand in hand with the amplified gene families, but not all are gorilla specific. For example, both human and gorilla have a similar number of *FRG1* paralogs, pointing to an initial amplification in a common ancestor. Under neutral evolution, these duplicated copies are expected to accumulate nonsense and frameshift mutations over time, but only 5 out of the 19 gorilla copies have pseudogenized compared to 17 of the 21 human copies (Supplementary Table S4;Tab1). Why have more gorilla copies maintained ORFs, and are they functional?

To demonstrate the coding potential of the amplified genes in gorilla, we measured their expression in fibroblasts and testes of the reference individual. Long-read, Iso-Seq transcriptome data mapped to the reference genome shows transcription of most genes in one or both tissues, with the combined expression of the duplicated copies often equal to or exceeding that of the ancestral copy (**Figure 6A**; Supplementary Tables S4;Tab1-2, S2). Thus, the positioning of these genes on the otherwise heterochromatic short arms does not silence their expression. We chose to focus specifically on the evolution and structure of the *FRG1* and the *IGSF3-GGT* fusion genes.

**Figure 6.**
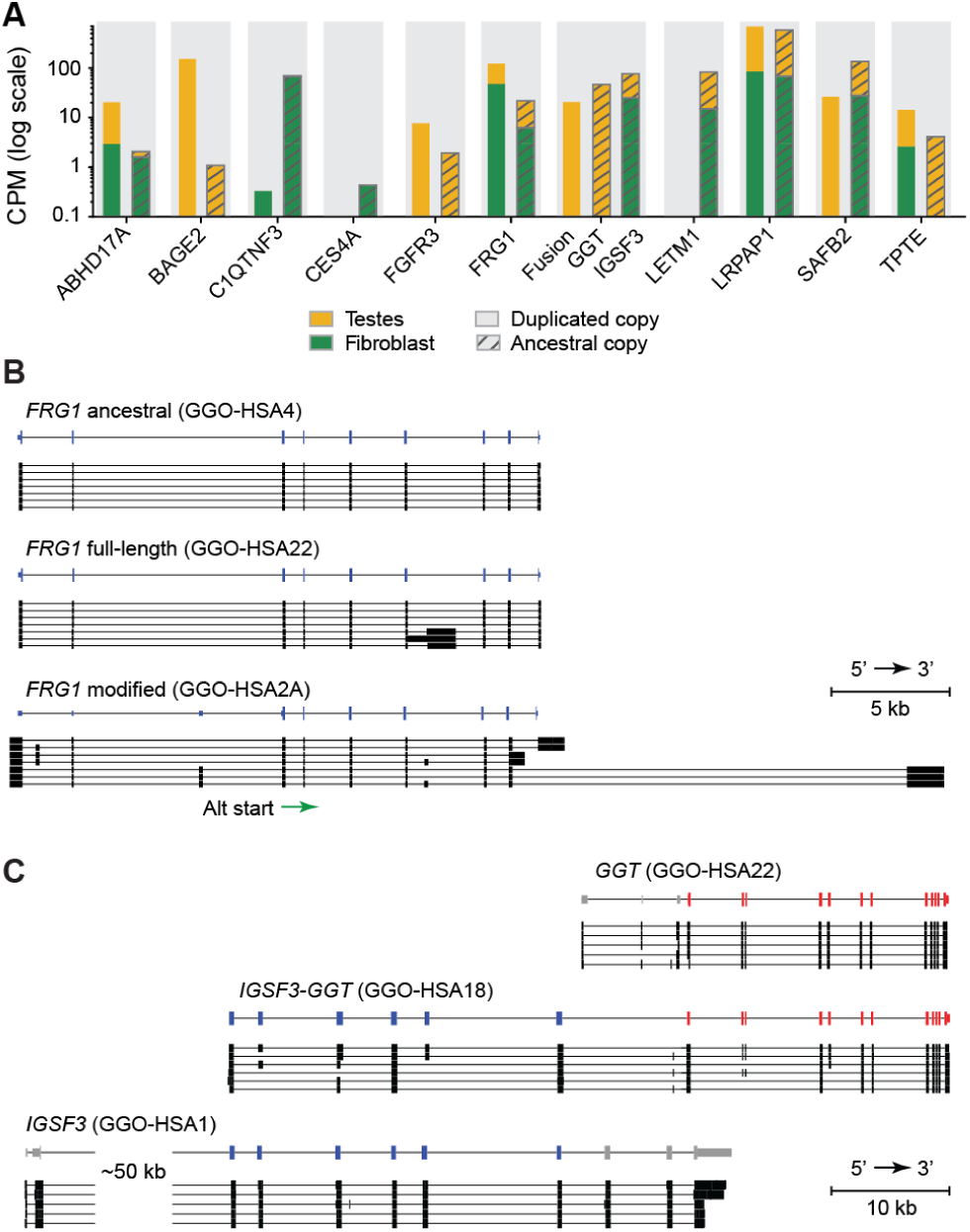
Expression and structure of amplified gorilla genes. **(A)** Expression (counts per million, CPM; log scale) of amplified protein-coding gene families on the gorilla acrocentrics, as measured by long-read PacBio Iso-Seq from testes (yellow) and fibroblasts (green) of the reference individual (n=4 technical replicates for fibroblast and n=1 technical replicates for testes). For each family, expression is partitioned between the amplified acrocentric copies (solid bars, “duplicated copy”) and the single ancestral copy (hatched bars, “ancestral copy”). “Fusion” denotes the *IGSF3-GGT* fusion gene, shown alongside its constituent *IGSF3* and *GGT* genes. **(B)** Transcript structure across the modified *FRG1* copies: the ancestral HSA4 copy present on GGO-HSA4; an example duplicated copy from GGO-HSA22; an example modified copy from GGO-HSA2A. The canonical gorilla *FRG1* transcript is shown at top. For each copy group, tracks show long-read transcript isoforms from the fibroblast Iso-Seq data. The green arrow marks the alternative start codon in exon 3 used by the modified copies. Untranslated exons have smaller heights. **(C)** Transcript structures of the *GGT1* (GGO-HSA22), *IGSF3* (GGO-HSA1), and *IGSF3-GGT* (example from GGO-HSA18). Gene structures are shown for *IGSF3*, *GGT*, and the *IGSF3-GGT* fusion with the *GGT* exons present in the fusion in red and the *IGSF3* exons present in the fusion in blue.

In addition to the ancestral copy on the orthologs of HSA4, all apes contain a large number of *FRG1* paralogs (11–38 copies per haplotype), but only gorilla and orangutan maintain a high number of predicted *FRG1* protein-coding copies (gorilla: 13, *B. orangutan*: 10, *S. orangutan*: 8) (Supplementary Table S4;Tab1). These fall into two structural classes: full-length and modified (**Figure 6B**). The full-length copies retain the canonical gene model and are found on the ends of GGO-HSA21/22, as well as all copies in orangutan. The modified copies lack the canonical start codon and instead initiate translation in exon 3, and are found in the pericentromeric PHR of GGO-HSA2A/13/15. In gorilla fibroblasts, both classes share a hypomethylated promoter preceding a hypermethylated gene body (Supplementary Fig. S28). This, along with the robust long-read transcript evidence, demonstrates that both duplicated and modified copies are expressed. In human and chimp, most duplicated *FRG1* copies have been pseudogenized by nonsense mutations, but a few modified copies maintain an ORF and are transcribed (human: 3, chimp: 1). Bonobo has the fewest number of *FRG1* paralogs and only the ancestral copy on PPA-HSA4 remains coding.

In contrast to *FRG1*, which is amplified in all the apes, the *IGSF3-GGT* fusion gene appears to be a gorilla-specific amplification that preserves a similar structure across all copies. The fusion product begins with six internal exons of *IGSF3* and ends with all coding exons of *GGT* (**Figure 6C**). Long read transcripts from testes show strong, but slightly lower, expression of the fusion gene compared to the isolated *GGT* copies (**Figure 6A**). Again, despite being surrounded by satellites, the amplified copies appear to escape silencing. Absence of *GGT* and *IGSF3-GGT* transcription in fibroblasts is consistent with *GGT* expression patterns in human, where it is most strongly expressed in the kidneys, liver, and intestine ^64^.

Whether these genes are under selection is harder to assess. Sequence exchange between the acrocentrics includes interlocus gene conversion (IGC), which is frequent within segmental duplications ^65^ and can confound tests of selection since it introduces recombinant gene sequences that cannot be described by a single phylogenetic tree ^66,67^. Additionally, tests based on dN/dS assume that paralogs diverge independently, but IGC acts to homogenize the copies, keeping both dN and dS low. With so few substitutions, dN/dS becomes unstable in either direction. Recent IGC can also restore an ORF in an existing paralog, so ORF retention alone may not imply constraint on all copies. We found evidence of IGC in most of the arrayed gene families, including *IGSF3-GGT* (Supplementary Table S5). Other genes such as *ABHD17A* and *LRPAP1* were difficult to align because of substantial variation in their gene models, indicative of either positive selection or relaxed constraint.

With these caveats in mind, we built phylogenies for the coding sequences (CDS) of each amplified gene family and tested for evidence of purifying, positive, and relaxed selection using HyPhy ^68^ (Supplementary Table S5, Supplementary Figures S29-40). To address some of the above limitations for the recently amplified gorilla copies, we focused on the branches leading to the amplification, which were generally stable between the alternative gene trees built from breaking the CDS at detected IGC breakpoints. Substitutions assigned to this branch will include shared changes that have spread among copies through their concerted evolution.

The most well-supported result was for *GGT*, whose inferred leading branch showed evidence of purifying selection (dN/dS = 0.29, 95% CI 0.16–0.49, p = 0.0064). This result is consistent with selection maintaining the shared *GGT* coding sequence of the amplified gene family, even if constraint on individual copies has relaxed. Branches within the clade do show dN/dS moving towards neutral (dN/dS = 0.99, CI 0.79–1.24), relaxation of selection (RELAX K = 0.17, p < 0.0001, where K<1 indicates relaxation), and a signal of episodic diversifying selection (BUSTED p = 0.0072). This may indicate that increased redundancy has reduced constraint of the individual genes, allowing for further diversification; however, the clade-level results should be interpreted with caution given the observed IGC. In contrast, the leading branch for the other half of the fusion, *IGSF3*, shows no evidence of purifying selection (dN/dS = 0.96, CI 0.62–1.41) and significant relaxation (RELAX K = 0.06, p = 0.044), suggesting lower constraint than the *GGT* portion. This could indicate promoter hijacking, in which the *IGSF3* portion was retained primarily to drive expression.

The *FRG1* amplification is not gorilla-specific, so we focused instead on the clade including all great ape copies descending from the ancestral HSA4 ortholog copies, as well as the subclade of modified copies in gorilla, human, and chimp. In contrast to the arrayed gene families, no IGC breakpoints were identified within *FRG1* and there were fewer substitutions on the leading branch, suggesting less homogenization among the older paralogs. Selection testing was therefore conducted across the great ape clade, restricted to the shared portion of the full-length and modified copies. No significant results were observed for the smaller, modified clade, but the full *FRG1* clade showed evidence of intensified (RELAX K = 2.67, p = 0.0001), positive diversifying selection (BUSTED p = 0.0012).

The other gene families show mixed results. For example, *LETM1*, *FGFR3*, and *SAFB2* are in the same gene cluster but show leading branch dN/dS values <1, ≈1, and >1 respectively (CIs: 0.21–1.11, 0.51–1.68, 0.90–4.62). However, these leading branches do not contain enough substitutions to draw firm conclusions, leading to the wide dN/dS confidence intervals. Thus, these results should be interpreted with caution, especially considering the lack of population-scale data, but the combined evidence of maintained ORFs, active expression, and substitution patterns differing substantially between adjacent genes is indicative of functional selection within a subset of the amplified gene families.

### Gene birth and death on the great ape acrocentrics

Shared gene amplifications with variable pseudogenization across the ape acrocentrics point to a birth and death evolutionary process, whereby initial amplification is followed by gradual decay of the genes not maintained by selection ^69^. To determine when these gene families were first amplified, we constructed time-measured phylogenies from their intron sequences, which are assumed to be evolving neutrally (**Figure 7**).

**Figure 7.**
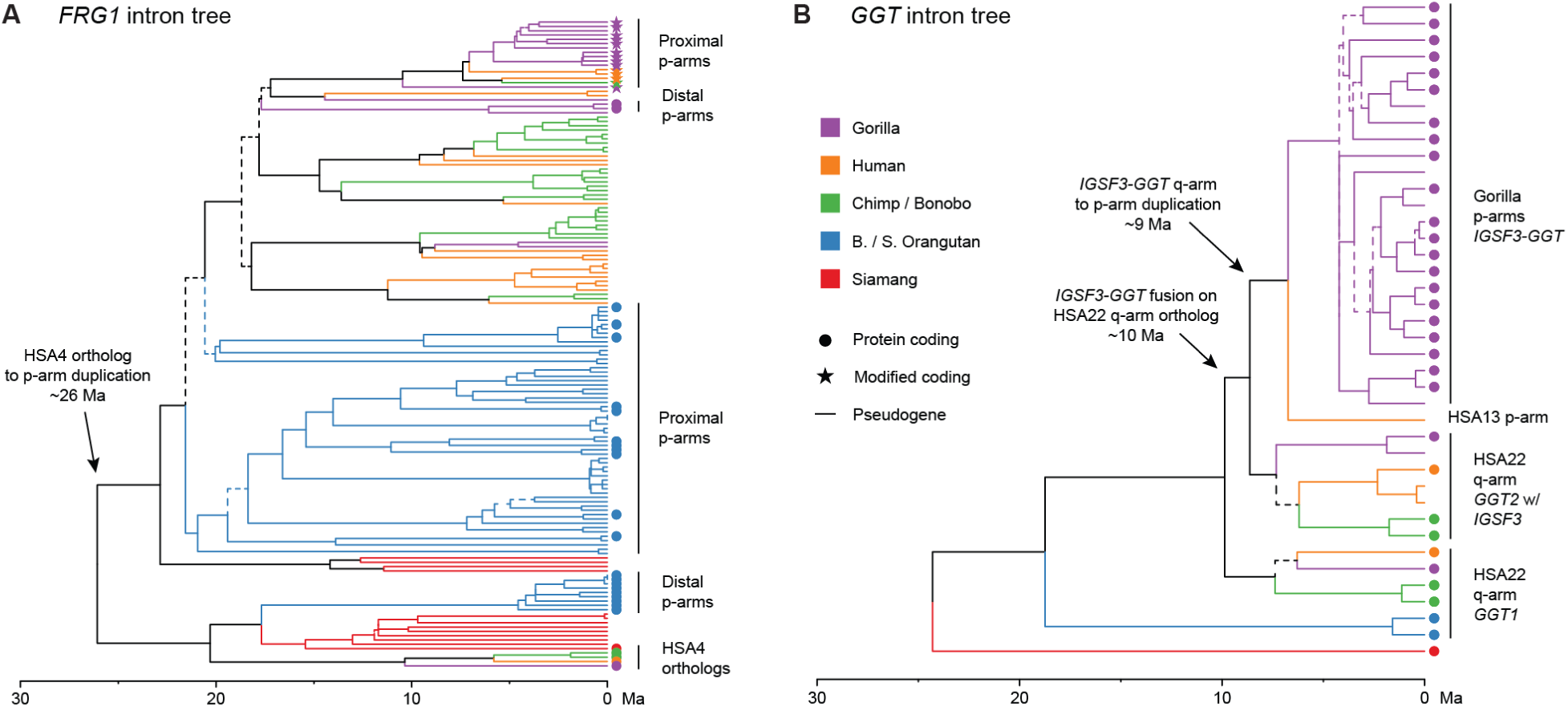
Timing of gene duplication onto the acrocentric short arms. **(A)** Time-measured intron phylogeny of all *FRG1* genes. **(B)** Time-measured intron phylogeny of all *GGT* genes including the *GGT* portion of the *IGSF3-GGT* fusion gene. Note, all *GGT2* copies in the African great apes have an upstream *IGSF3* fragment, reflective of the ancestral fusion event, but only *GGT2* has maintained coding potential on the long arms. Intact *IGSF3-GGT* fusion copies are exclusive to the short arms of the gorilla acrocentrics, with one additional pseudogenized copy on the short arm of HSA13. For both trees, branches with a bootstrap <90% are dashed, and marmoset and macaque were used as outgroups (not shown). Fully labeled trees, including bootstraps and split times, are given in Supplementary Figs. S41 and S42.

As noted above, all apes (including siamang) have a large number of *FRG1* copies, pointing to an ancient amplification in their common ancestor. The *FRG1* intron phylogeny supports this hypothesis and places an initial duplication of the ancestral HSA4 sequence onto the short arms of the acrocentrics 24.8–27.4 million years ago (Ma), from where it repeatedly amplified and diversified (**Figure 7A**, Supplementary Fig. S41). All additional *FRG1* copies in human, chimp, bonobo, and gorilla derive from this ancestral duplication and do not cluster cleanly by species, consistent with their amplification in a common ancestor. These copies coalesce 16.8–20.9 Ma, corresponding with the currently estimated human/orangutan split 18.2–19.6 Ma ^11^. The modified human and chimp copies group with the modified gorilla copies, consistent with a single origin of the modified structure in a common ancestor. Orangutan and siamang also show evidence of this original duplication/amplification, as well as an independent and more recent amplification of the HSA4 copy ∼20 Ma, which has since spread across the distal ends of the orangutan short arms and completely pseudogenized in siamang. Thus, the large number of *FRG1* copies observed in modern apes likely arose from ancestral duplications onto the short arms followed by lineage-specific amplification and differential retention.

Although similar in mechanism, the *IGSF3-GGT* amplification is more recent. This fusion was first reported on the long arm of HSA22 ^70^, and we find that all *GGT2* copies in the African great apes are preceded by a segment of *IGSF3*, indicating that the fusion originated in their common ancestor. The *GGT* intron phylogeny places this duplication and fusion with *IGSF3* on the long arm of the HSA22 ancestor 8.6–11.6 Ma (**Figure 7B**, Supplementary Fig. S42), around the estimated split between human and gorilla 10.6–10.9 Ma ^11^. From here, the fusion was duplicated onto the short arms of the acrocentrics 7.2–10.0 Ma, and the two loci followed different fates. On the long arms of all species, the *IGSF3* segment has been degraded and no longer yields fusion transcripts, while on the short arms of gorilla, the fusion has been maintained and amplified alongside *TPTE* (**Figure 5C**). In CHM13, a single full-length *IGSF3-GGT* fusion copy also remains on the short arm of HSA13 next to *TPTE*, but has been pseudogenized by nonsense mutations. CDS trees for both *IGSF3* and *GGT* show accelerated evolution of the fusion gene after duplication onto the short arms, especially for the GGO-HSA9/18 copies (Supplementary Figs. S35 and S36). The GGO-HSA21/22 *GGT* coding sequences branch early in this tree, paralleling the SST1 tree and diversification of SST1′ within the GGO-HSA9/18 PHRs (**Figure 4**). Together, the shared satellite patterns and phylogenies demonstrate that the gorilla *IGSF3-GGT* copies are evolving in concert.

## Discussion

We have shown that acrocentric chromosomes played a key role in the evolution of our ancestors and continue to play an important role in our health today. Leveraging the first complete genomes of the great apes, our sequence-level analysis revealed common features of the short arms that were unclear from cytogenetics alone, including (1) highly variable composition of segmental duplications and specific satellite classes such as ACRO, SST1, HSat1A, and HSat3, (2) recurrent whole-arm inversions converting acrocentrics to (sub)metacentrics and forming potential speciation barriers, (3) non-allelic sequence exchange between the short arms of all great apes, leading to concerted evolution of more than the rDNAs, and (4) accelerated birth-and-death evolution of multiple protein-coding gene families, enabled by these unique characteristics of the short arms.

Pericentric and paracentric inversions are the primary driver of karyotype evolution in the great apes, driving speciation through suppressed meiotic recombination in heterokaryotypes ^10^. We found that at least one whole short-arm inversion event can be placed on each major branch of the species tree, converting acrocentric chromosomes to metacentric chromosomes and forming distinct karyotypes among the major ape lineages. This fits the hypothesis that pericentric inversions are statistically directional, i.e. they favor the conversion of acrocentric to metacentric chromosomes ^71^, which may then become fixed in the population through selection, meiotic drive, or genetic drift ^72^. The result is a progressive loss of acrocentrics in the great apes, and a trend towards metacentrics compared to the ancestral karyotype. Notably, the two species that have experienced the fewest acrocentric to metacentric conversions, orangutan and gorilla, both retain large, transcribed amplified gene families on their short arms (e.g. *FRG1*), potentially countering the forces leading to acrocentric loss in the other species.

Further, we have shown that the unique compositional and recombinational properties of the short arms, including satellite DNAs and PHRs, lend themselves not only to the homogenization of rDNA arrays but also the amplification of satellites and multicopy gene families through birth-and-death evolution. The short arms of the acrocentrics appear especially tolerant of genomic repeats, perhaps due to a suppression of meiotic recombination ^56^, similar to what is observed within centromeres and the non-recombining region of ChrY. However, enough sequence exchange remains between the short arms to facilitate sequence amplification and concerted evolution. Our results suggest that many of the pericentromeric repeats observed across the human genome, such as the HSat3 arrays, were first seeded and amplified on the short arms of acrocentric chromosomes before being moved proximally by a chromosomal fusion or inversion. This may include Chr16, whose ancient conversion to metacentric and subsequent cessation of sequence exchange with the acrocentrics could explain the divergence of its pericentromeric HSat2 array.

We have primarily focused on the evolution of acrocentric chromosomes following the human and orangutan split ∼19 Ma, but not the events leading to their emergence in the great ape ancestor. This is a more complicated picture involving multiple chromosome fission and fusion events, and only Chr13 and ChrY are acrocentric in both human and the inferred primate common ancestor ^5^. The hypothesized ancestral primate karyotype contained as many as 14 acrocentric chromosomes ^40^, many presumably carrying rDNA. Some of these ancient acrocentrics were lost by fusions, including orthologs of HSA 10, 16, and 19, and others by short-arm inversions, like those presented here. In the Old World monkeys, *Macaca mulatta* lost all acrocentric rDNA arrays and now carries a single pericentromeric rDNA array resulting from the fusion of HSA20 and HSA22 orthologs ^37,73^. Conversely, new acrocentrics have been created by fission of the ancestral primate karyotype, including HSA14/15, HSA3/21, and HSA12/22 orthologs ^40,74^. The subsequent gain of rDNA arrays on these chromosomes points to a general tendency of acrocentric chromosomes to exchange and acquire rDNA from each other, and a similar mechanism of rDNA gain through centric fission and satellite expansion has been hypothesized in ants ^75^. Thus, karyotype evolution by rDNA-associated fission and fusion events may be a general mechanism contributing to eukaryotic speciation.

It is debatable whether the occurrence of multiple rDNA arrays, as seen in the great apes, is advantageous or simply a consequence of the different selective pressures acting on the short arms of acrocentric chromosomes. Dispersing onto multiple chromosomes may offer the benefit of redundancy and help balance overall rDNA copy number and fidelity, reducing the impact of deleterious variants, array silencing, or array loss ^29^. Combined with strong purifying selection, the mechanisms of molecular drive can maintain a high degree of similarity within and between rDNA arrays ^33,35^. Due to their propensity for exchange, selection appears to favor either a single pericentromeric rDNA array, as in *M. mulatta*, or multiple, uniformly oriented rDNA arrays on the ends of chromosomes, as in the great apes, since a crossover between either proximal or inverted distal arrays would result in a translocation and subfertility.

The gorilla genome illustrates that these mechanisms of redundancy and molecular drive are not limited to the rDNA arrays and can involve the entire short arms of rDNA− chromosomes, possibly facilitated by homologous recognition between their shared heterochromatin ^76,77^. This has enabled the amplification and concerted evolution of protein-coding genes in a manner similar to the rRNA genes. Thus, given their ability to flexibly amplify and recombine all manner of genomic sequences with minimal fitness costs, the short arms of acrocentric chromosomes appear to be underappreciated hubs of genomic innovation. Similar effects have also been reported within the subterminal heterochromatin caps of chimp and gorilla chromosomes ^52^. As Ohno stated in summarizing the importance of duplications in evolution, “natural selection merely modified, while redundancy created” ^78^. In this way, the redundancy generated by interchromosomal exchange between the acrocentrics generates additional variation for natural selection to act upon, and the degree of homogenization versus diversification can be fine-tuned by the rate of exchange. This manifests as both the highly homogenized nature of the tandemly repeating rRNA genes (high rates of exchange) and the segmentally duplicated PHRs (lower rates of exchange). For example, by decoupling from the nucleolus, the gorilla rDNA− acrocentrics have been freed to fine-tune their own degree of concerted evolution. These PHRs will continue to evolve in concert until random mutation outpaces the rate of interchromosomal exchange and the process halts ^79^, as seen with sequences on chromosomes that have converted from acrocentric to metacentric, thus, lowering their rate of exchange and allowing mutations to dominate.

The amplified gene families found on the gorilla acrocentrics obviously raise questions of adaptation, and these genes are broadly associated with common drivers of rapid evolution, including immunity and inflammation (*ABHD17A*, *C1QTNF3, GGT*, *SAFB2*), metabolism and detoxification (*C1QTNF3*, *CES4A*, *GGT*, *LRPAP1*), musculoskeletal development (*FGFR3*, *FRG1*), reproduction (*SAFB2*, *TPTE, FGFR3*), and cancer/testis genes (*BAGE2, TPTE*). Since many of these copies have maintained intact, transcribed ORFs over millions of years of amplification, it is tempting to speculate how these unique genomic features may be linked to gorilla-specific traits such as their unique diet and sexual dimorphism. The *FRG1* and *IGSF3-GGT* gene families provide a beautifully contrasting view of birth-and-death evolution in the apes, with the newer *IGSF3-GGT* copies recently “born” and the much older *FRG1* copies having mostly pseudogenized outside of gorilla and orangutan. While our selection results were mixed, similar patterns have been reported for multicopy gene families on the great ape Y chromosomes ^80^. There, substitutions predicted to alter protein function were typically expressed alongside unaltered copies, suggesting that functional conservation can accompany sequence diversification within gene arrays, mirroring our observations for *IGSF3-GGT*.

In human, remnants of the ancient *FRG1* amplifications persist on the acrocentrics, and future work is needed to better understand the potential function of these previously inaccessible genes. Many of these *FRG1* copies localize to the same PHRs recently implicated in Robertsonian formation ^21^. Other remnants of past amplifications remain polymorphic, including the pericentromeric heterochromatin of Chr9 and Chr16 ^81^, where variation and epigenetic dysregulation have potential health implications ^54,82^. Evolution is often a double-edged sword—the processes that may have orchestrated adaptation in our ancestors now predispose us to certain genomic disorders. More T2T genomes are needed to better understand the role of duplication and accelerated evolution within these once “dark” areas of the genome.

### Limitations of the Study

The highly repetitive short arms remain challenging to sequence and assemble, restricting our analysis to a single T2T genome per species and limiting our ability to study intraspecific variation beyond a single pair of haplotypes. As a result, our PHR definitions are only approximate. Additionally, the rapid turnover of satellite repeats limited our ability to precisely identify the short-arm inversion breakpoints, and our reliance on parsimony for mapping inversion events to the great ape tree disregards the possibility of convergent or reversed inversions. Lastly, our analyses rely on publicly available T2T genome assemblies, which were thoroughly validated by previous studies but could still contain some residual misassemblies (most likely as expansions or collapses with near-identical repeats).

Inference of the evolutionary history and function of the amplified gene families was constrained by ongoing sequence exchange between the copies, lack of population data, and limited functional evidence. Selection estimates from a single phylogeny may be unreliable when sequence exchange is not modeled, and the lack of population data further limits inference. Similarly, our divergence estimates may be affected by sequence exchange involving the introns. Lastly, functional potential of the annotated genes was primarily inferred from ORF maintenance and expression evidence, and will require additional functional validation to confirm.

## Methods

### Key Resources Table

| REAGENT or RESOURCE | SOURCE | IDENTIFIER |
| --- | --- | --- |
| **Antibodies** |  |  |
| **Bacterial and virus strains** |  |  |
| **Biological samples** |  |  |
| **Chemicals, peptides, and recombinant proteins** |  |  |
| PureLink Quick Plasmid Miniprep Kit | Invitrogen | K210011 |
| Cy3-dUTP | Sigma-Aldrich | GEPA53022 |
| human Cot DNA | Invitrogen | 15279101 |
| SSC | Sigma-Aldrich | S6639 |
| Hybridization Mix | Sigma-Aldrich | H7782 |
| KaryoMAX colcemid | Thermo Fisher | Cat#: 11519876 |
| RNase A | Qiagen | Cat#: 19101 |
| VECTASHIELD mounting medium with DAPI | Vector Laboratories | Cat#: H-1200-10 |
| Streptavidin-Cy5 | Thermo Fisher | Cat#: 434316 |
| **Critical commercial assays** |  |  |
| **Deposited data** |  |  |
| **Experimental models: Cell lines** |  |  |
| Western lowland gorilla fibroblast cell line GGO | Ventura Lab at University of Bari | GGO |
| Western lowland gorilla fibroblast cell line GGO1 | Ventura Lab at University of Bari | GGO1 |
| Western lowland gorilla fibroblast cell line GGO2 | Ventura Lab at University of Bari | GGO2 |
| Western lowland gorilla fibroblast cell line GGO5 | Ventura Lab at University of Bari | GGO5 |
| Western lowland gorilla fibroblast cell line GGO9 | Ventura Lab at University of Bari | GGO9 |
| Western lowland gorilla fibroblast cell line AG20600 | Ventura Lab at University of Bari | GGO14 |
| Western lowland gorilla fibroblast cell line KB3781 | San Diego Zoo Wildlife Alliance | KB3781 |
| Western lowland gorilla fibroblast cell line AG20600 | Coriell Institute | AG20600 |
| Western lowland gorilla fibroblast cell line AG21765 | Coriell Institute | AG21765 |
| Western lowland gorilla fibroblast cell line AG05251 | Coriell Institute | AG05251 |
| **Recombinant DNA** |  |  |
| BAC-clone RP11-341D18 | CHORI-BACPAC genomic resources | RP11-341D18 |
| BAC-clone RP11-529E10 | CHORI-BACPAC genomic resources | RP11-529E10 |
| BAC-clone RP11-450E20 | CHORI-BACPAC genomic resources | RP11-450E20 |
| BAC probe RP11-450E20 (rDNA), fluorescently labeled | Empire Genomics | RP11-450E20 |
| BAC probe RP11-341D18 (AL356585.7), biotin labeled | Empire Genomics | RP11-341D18 |
| BAC probe RP11-529E10 (AL590235.23), biotin labeled | Empire Genomics | RP11-529E10 |
| Human whole chromosome 21 paint | Applied Spectral Imaging | FPRPR0167 |
| **Software and algorithms** |  |  |
| AniAnn's | Sweeten et al. <sup>83</sup> | github.com/marbl/anianns, git commit 8dd4e1b |
| ModDotPlot v0.9.0 | Sweeten et al. <sup>84</sup> | https://github.com/marbl/ModDotPlot/ |
| minimap2 v2.26 | Li <sup>85</sup> | https://github.com/lh3/minimap2/ |
| Rustybam v0.1.33 | Vollger <sup>86</sup> | https://github.com/vollgerlab/rustybam/ |
| Wfmash v0.21.0 | Garrison et al. | https://github.com/waveygang/wfmash/ |
| SVbyEye | Porubsky et al. <sup>87</sup> | https://github.com/daewoooo/SVbyEye |
| MashMap3 v3.1.3 | Kille et al. <sup>88</sup> | https://github.com/marbl/MashMap/ |
| NetworkX | Hagberg et al. <sup>89</sup> | https://github.com/networkx/networkx/releases#release-networkx-3.4 |
| BLAT | Kent <sup>90</sup> | https://genome.ucsc.edu/cgi-bin/hgBlat |
| IsoQuant v3.10.0 | Prjibelski et al. <sup>91</sup> | https://github.com/ablab/IsoQuant/ |
| samtools v1.21 | Danecek et al. <sup>92</sup> | https://www.htslib.org/ |
| Guppy v6.3.8 | Oxford Nanopore Technologies | https://community.nanoporetech.com/ |
| Winnowmap v2.03 | Jain et al. <sup>93</sup> | https://github.com/marbl/Winnowmap |
| meryl | Rhie et al. <sup>94</sup> | https://github.com/marbl/meryl |
| modkit v0.6.1 | Oxford Nanopore Technologies | https://github.com/nanoporetech/modkit |
| MAFFT v7.5 | Katoh et al. <sup>95</sup> | https://mafft.cbrc.jp/alignment/software/source.html |
| RAXML-NG v1.2.2 | Kozlov et al. <sup>96</sup> | https://github.com/amkozlov/raxml-ng |
| MEGA 12 | Kumar et al. <sup>97</sup> | https://www.megasoftware.net/ |
| BUSTED (HyPhy) | Murrell et al. <sup>98</sup> | https://datamonkey.org/ |
| RELAX (HyPhy) | Wertheim et al. <sup>99</sup> | https://datamonkey.org/ |
| GARD (HyPhy) | Pond et al. <sup>67</sup> | https://datamonkey.org/ |
| SLAC (HyPhy) | Pond et al. <sup>100</sup> | https://datamonkey.org/ |
| IQ-TREE2 v2.4.0 | Minh et al. <sup>101</sup> | https://github.com/iqtree/iqtree2 |
| RepeatMasker | Smit et al. <sup>102</sup> | https://www.repeatmasker.org/ |
| Dotlet | Junier et al. <sup>103</sup> | https://dotlet.vital-it.ch/ |
| jModelTest2 | Darriba et al. <sup>104</sup> | https://github.com/ddarriba/jmodeltest2 |
| Fiji (ImageJ) | Schindelin et al. <sup>105</sup> | https://fiji.sc/ |
| IPLab | Scanalytics / BD Biosciences | https://bio-formats.readthedocs.io/en/v7.3.0/formats/iplab.html |
| Adobe Photoshop | Adobe | https://www.adobe.com/products/photoshop.html |
| **Other** |  |  |
| Nikon TiE spinning-disk confocal microscope (100× NA 1.45 objective; Yokogawa CSU-W1; ORCA-Fusion BT C15440 CMOS camera) | Nikon / Yokogawa / Hamamatsu | N/A |
| Leica DMRXA epifluorescence microscope with cooled CCD camera | Leica / Princeton Instruments | N/A |
| PacBio Iso-Seq long-read transcript sequencing | Pacific Biosciences | N/A |

### Experimental model and study participant details

#### Gorilla cell lines

Western lowland gorilla (*Gorilla gorilla gorilla*) fibroblast cell lines were obtained from the San Diego Zoo Wildlife Alliance (KB3781) and from the Coriell Institute (AG20600, AG21765, and AG05251). KB3781 (“Jim”) is the same animal originally used for the T2T gorilla reference assembly. All cell lines were initially cultured in alpha-MEM (Gibco) with 2 mM L-glutamine supplemented with 15% fetal bovine serum (FBS) and were subsequently switched to DMEM/F12 (Gibco) with 15% FBS. Cell cultures were maintained at 37°C with 5% CO2.

For independent replication of FISH experiments, additional *Gorilla gorilla gorilla* skin-derived fibroblast cell lines were obtained from the Ventura laboratory at the University of Bari (GGO, GGO1, GGO2, GGO5, GGO9, and GGO14). GGO14 is the same individual as AG20600 (above); all other Ventura-lab lines are unique individuals derived from European zoos. In total, nine unique gorilla individuals were analyzed by FISH. This study did not involve human participants or vertebrate animal procedures; all analyses used established cell lines and publicly available genome assemblies.

### Method details

#### Satellite and rDNA annotations

Satellite annotations were primarily drawn from prior work ^11^, but a few gaps were identified in the original annotation. To fill these gaps, the genomes were additionally annotated using AniAnn’s (github.com/marbl/anianns, git commit 8dd4e1b). AniAnn’s identifies tandem repeats by splitting each sequence into windows (2,000 bp) and measuring the average nucleotide identity (ANI) between adjacent windows. Previously unannotated regions exceeding an ANI threshold of 86% were identified as unknown satellite arrays and added as “other” satellites to the original annotation.

~~~
anianns annotate -f $species.fa -c $kmer_dir > $species.bed
~~~

All percent identity dotplots were created using ModDotPlot v0.9 ^84^.

~~~
moddotplot static -f $species.fa -a $max_length
~~~

To calculate satellite enrichment on the acrocentrics, the above satellite annotations were subsetted for various satellite classes and partitioned by species and chromosome type.

To show the orientations of rDNA arrays are consistent within species, each ape genome was mapped to the human 45S gene extracted from the KY962518.1 rDNA reference sequence ^106^ using minimap2 ^85^. Mappings were inverted and filtered for a minimum of 95% identity and over a 10 kb alignment block to ensure accuracy and inspected in the UCSC Genome Browser ^107^.

~~~
minimap2 -c -x asm20 --eqx --MD -t 2 45S.fa $species.fa > $species.to.45S.paf
rustybam invert $species.to.45S.paf > 45S.to.$species.paf
awk -v OFS=’\t’ ’{if($11>=10000 && $10/$11 >= 0.95) print $6,$8,$9, $10/$11}’
45S.to.$species.paf > 45S.to.$species.filt.bed
~~~

The same was done for the DJ, aligning a representative copy from CHM13 Chr13 but filtering for 100 kb matches at 90% identity.

~~~
minimap2 -c -x asm20 --eqx --MD -t 2 DJ_palindrome.fa $species.fa >
$species.to.DJ_palindrome.paf
rustybam invert $species.to.DJ_palindrome.paf > DJ_palindrome.to.$species.paf
awk -v OFS=’\t’ ’{if($11>=100000 && $10/$11 >= 0.9) print $6,$8,$9, $10/$11}’
DJ_palindrome.to.$species.paf > DJ_palindrome.to.$species.filt.bed
~~~

The above DJ alignment and filtering methods were also used to identify remnants of the DJ on the chimp and bonobo Y chromosomes.

#### Pairwise alignment of whole chromosomes

Primary haplotypes for each chromosome belonging to the superset of great ape acrocentric chromosomes (HSA 2A, 2B, 9, 13, 14, 15, 18, 21, 22, Y) were extracted from each species and renamed to indicate the species of origin and HSA chromosome designation. HSA9 and Y were inverted in chimp and bonobo to be consistent with the other great apes. Whole chromosomes were aligned pairwise using wfmash (https://github.com/waveygang/wfmash, version 0.21.0) with default parameters, except for a 50 kb seed size to reduce spurious alignments in repetitive sequences.

~~~
wfmash -t 16 -s 50000 -p 85 $fa_2 $fa_1 > $out.paf
~~~

The resulting PAF files were concatenated to contain one of each species pair in a central PAF file for each homologous chromosome. The resulting alignments and corresponding satellite annotations were visualized using the AVA mode of SVbyEye ^87^, showing large synteny blocks and inversions between pairs of species.

#### Pseudo-homolog region identification

To identify PHRs, each chromosome was compared in a pairwise fashion with MashMap3 ^88^ to all other chromosomes using 10 kb chunks and an identity threshold of 85%. The resulting matches were filtered to only retain hits above 99% identity for the purpose of identifying putative PHRs. When tested on CHM13, this threshold showed a 93% sensitivity and 70% specificity for recovering the PHR bases previously identified by Guarracino *et al.*, excluding satellites and telomeres ^34^. To generate heatmap tracks for viewing, the best MashMap hit to any position on the query chromosome was retained for each 10 kb window on the reference chromosome. A custom script was used to convert percent identity to an RGB value on the YlOrRd scale, ranging from 80–100%, with values below excluded and appearing white.

The mappings were then summarized to calculate the amount of shared sequence above 99% identity between all pairs of chromosomes in each species. Total bases and average percent identity of the shared sequence were averaged for each diploid pair of chromosomes to obtain haploid counts. A custom script using NetworkX ^89^ was used to plot the interchromosomal relationships. The average percent identity of shared sequence between chromosomes was plotted as line color by filtering alignments for those above 99% identity, subtracting 99%, and normalizing between 0 and 1. The line width was normalized by dividing the total shared sequence by 2.5 Mb and using this value as the line width in pixels. Since the amount of shared sequence between chromosomes was dominated by satellite DNA, rDNA, and subterminal spacers in the pCht arrays, these were filtered to isolate the segmental duplications.

The segmental duplication structure of the GGO-HSA9/18/21 amplifications was first assessed with dotplots and satellite annotations. This revealed three approximate gene clusters of varying lengths and orientation. One representative cluster was extracted for each class and realigned to the gorilla genome with minimap2 to identify all instances of each cluster aligning with at least 90% average identity over 100 kb.

#### Amplified gene annotation

Initial gene annotations for the short arms of the gorilla acrocentrics were taken from NCBI RefSeq ^108^ and Comparative Annotation Toolkit (CAT) annotation datasets ^11^. Due to incompleteness in both count and transcript-level detail within the duplicated regions, we performed systematic manual curation of all genes appearing at least once on the short arms of the gorilla acrocentrics. For these genes, all potential paralogs were identified across the great apes using BLAT ^90^ sequence alignment against the complete assemblies and visualized in the UCSC Genome Browser. Each paralog was manually assessed for protein-coding potential by considering exon structure integrity, nonsense mutations leading to pseudogenization, and transcriptional activity confirmation using long-read PacBio Iso-Seq transcripts from testes and fibroblast samples of the reference individual (KB3781).

#### Gene family alignment and phylogenetics

Protein-coding copies for each gene family were extracted from the curated annotations and aligned using MAFFT v7.5 with default parameters ^95^. Phylogenetic relationships were reconstructed using maximum likelihood methods implemented in RAxML-NG under a GTR+G+I substitution model with default settings ^96^. Before conducting selection tests, frameshifts and novel exons in the CDS alignments were trimmed using MEGA to visualize and edit the alignments ^97^.

To estimate the age of *FRG1* and *GGT* duplications, intronic sequences were extracted from both gene bodies and aligned with MAFFT using parameters:

~~~
--anysymbol --reorder --maxiterate 1000 --thread 16
~~~

The alignment was then processed with IQ-TREE2 ^101^ using parameters:

~~~
--date {divergence_time_file} --keep-ident --date-tip 0 --date-ci 100 -B 1000 -T 36
~~~

to generate a maximum likelihood phylogeny that was bootstrapped using UFBoot2 ^109^ with 1,000 replicates and dated using LSD2 ^110^, using marmoset and macaque as outgroups, with an estimated human/macaque divergence time of 28.8 Ma and human/marmoset divergence time of 43 Ma. In the resulting phylogenies, branches with bootstrap support <90% are indicated as dashed.

#### SST1 monomer characterization

SST1 arrays were initially identified using RepeatMasker ^102^. For each identified array, sequences were retrieved along with 1 kb of flanking sequence. All clusters were manually curated through visual inspection. This was performed by generating dotplots using the Dotlet applet ^103^ with a word size of 15 bp and a similarity cut-off of 60%. The consensus sequences were refined through an iterative process, which enabled precise sequence characterization. This manual curation allowed for the definitive identification of the start and end points of each array and its constituent monomers relative to the generated consensus. To ensure consistency in subsequent alignments, all monomeric sequences were characterized using identical start and end points defined by the consensus. All retrieved full-length SST1 monomeric sequences from assembled genomes were aligned using MAFFT with the G-INS-1 model. A maximum likelihood phylogenetic analysis was conducted under the best-fit nucleotide substitution model (GTR +G, with a shape parameter [α] = 5.5047), as determined by jModelTest2 ^104^. The robustness of the inferred topology was assessed with 1,000 bootstrap replicates.

#### DNA methylation analysis

Oxford Nanopore reads from the gorilla mGorGor1 individual basecalled with Guppy v6.3.8 were aligned to the T2T genome (GCA_029281585.2) with Winnowmap v2.03 ^93^ (-ax map-ont -k 15 -y), using a meryl ^94^ 15-mer table of high-copy k-mers. Library BAMs were merged with samtools v1.21 ^92^ and restricted to alignments with mapping quality ≥ 60. Site-level CpG methylation was summarized with modkit v0.6.1 (pileup --preset traditional --force-allow-implicit), which reports 5mC at CpG dinucleotides, combines strands, and treats implicitly canonical bases as unmodified.

#### Gorilla fluorescence in situ hybridization

Two human BAC clones were independently tested by FISH experiments in gorilla to confirm the primary amplifications were not isolated to the assembled individual: RP11-341D18 (AL356585.7), overlapping the *IGSF3*-*GGT* amplification, and RP11-529E10 (AL590235.23), overlapping the *LRPAP1* (*FRG1*) amplification.

For the preparation of chromosome spreads, cells were blocked in mitosis by the addition of KaryoMAX colcemid solution (0.1 µg/ml, Life Technologies) for 6–7 h and collected by trypsinization. Collected cells were spun down and incubated in hypotonic 0.4% KCl solution for 12 min and prefixed by addition of methanol:acetic acid (3:1) fixative solution (1% total volume). Pre-fixed cells were spun down and then fixed in methanol:acetic acid (3:1). Chromosome spreads were dropped on a glass slide and incubated at 65°C overnight. Before hybridization, slides were treated with 0.1 mg/ml RNase A (Qiagen) in 2× SSC for 45 min at 37°C and dehydrated in a 70%, 80%, and 100% ethanol series for 2 minutes each. Slides were denatured in 70% deionized formamide/2× SSC solution pre-heated to 72°C for 1.5 min. Denaturation was stopped by immersing slides in a 70%, 80%, and 100% ethanol series chilled to −20°C. Fluorescently and biotin-labeled BAC probes RP11-450E20 (rDNA), RP11-341D18 (*GGT*), and RP11-529E10 (*LRPAP1*) were obtained from Empire Genomics. Human Chromosome 21 paint was obtained from Applied Spectral Imaging. Probes used in combination were denatured in a hybridization buffer by heating to 80°C for 7 minutes before applying to denatured slides. Specimens were hybridized to the probes under a HybriSlip hybridization cover (GRACE Biolabs) sealed with Cytobond (SciGene), in a humidified chamber at 37°C for 48 h. After hybridization, slides were washed in 50% formamide/2× SSC three times for 5 min per wash at 45°C, then in 1× SSC solution at 45°C for 5 minutes twice and at room temperature once. For biotin detection, slides were incubated with streptavidin conjugated to Cy5 (Thermo) for 2–3 h in PBS containing 0.1% Triton X-100 and 5% bovine serum albumin (BSA), and then washed 3 times for 5 minutes with PBS/0.1% Triton X-100. Slides were rinsed in water, air-dried in the dark, and mounted in VECTASHIELD containing DAPI (Vector Laboratories). Z-stack confocal images of chromosome spreads and interphase nuclei were acquired on a Nikon TiE microscope equipped with a 100× objective (NA 1.45), Yokogawa CSU-W1 spinning disk, and ORCA-Fusion BT C15440 CMOS camera (Hamamatsu). Image processing was performed in Fiji ^105^.

FISH experiments with RP11-341D18 were independently replicated for additional *Gorilla gorilla gorilla* skin-derived cell lines from the Ventura lab at the University of Bari (GGO, GGO1, GGO2, GGO5, GGO9, GGO14). For this preparation, all BACs were isolated using the PureLink Quick Plasmid Miniprep Kit (Invitrogen), labeled by nick translation with Cy3-dUTP, and subsequently subjected to ion-exchange alcohol precipitation in the presence of human Cot DNA ^111^. Following denaturation at 70°C for 2 minutes, hybridization was performed overnight at 37°C. Post-hybridization washes were performed three times in 0.1× SSC at 60°C under high-stringency conditions. Slides were counterstained with DAPI, and fluorescence signals from Cy3 and DAPI were independently detected using a Leica DMRXA epifluorescence microscope equipped with a cooled CCD camera (Princeton Instruments). Images were acquired in grayscale, pseudo-colored, and merged using Adobe Photoshop. Signal intensity was recorded semi-quantitatively (+ to ++++) and localization noted.

### Quantification and statistical analysis

#### Expression quantification

For each amplified gene family, counts per million (CPM) were calculated using IsoQuant ^91^ for each sample type (testes and fibroblast) separately and combined. We used 4 technical replicates for fibroblast and 1 technical replicate for testes. Reads were first aligned using minimap2 v2.26 (-ax splice:hq -uf), and then sorted and indexed with samtools v1.21. IsoQuant v3.10.0 was then run using default parameters and the consensus NCBI/CAT gene annotations. Expression was partitioned between the amplified acrocentric copies and the ancestral copy for each family. Full-length reads exhibiting the modified transcript structure characteristic of the duplicated copies were assigned to the duplicated class. Reads with canonical structure were assigned by unique alignment, requiring a mapping quality (MAPQ) ≥ 40 to attribute a read to a single locus; the divergence of the ancestral copy from the amplified cluster was generally sufficient to permit unique mapping. For reads with canonical structure that mapped ambiguously between the ancestral and duplicated copies, the partition relies on mapping quality alone and the ancestral and duplicated contributions may not be fully separable.

#### Selection analysis

Gene-wide tests for selection were performed on the CDS alignments. Alignments were first trimmed to remove large gaps and cleaned with hyphy clean-names.bf. They were then screened for recombination using GARD ^67^; inferred breakpoint positions, or their absence, are reported for each gene. dN/dS was estimated with hyphy FitMG94.bf with --type local --lrt Yes to get both the mean dN/dS value with a 95% confidence interval on the test branch along with the likelihood-ratio p-value for the null hypothesis that mean dN/dS = 1; the branch length is also reported for the leading-branch analysis. The inferred numbers of nonsynonymous (nN) and synonymous (nS) substitutions were estimated with SLAC ^100^. Episodic diversifying selection was tested with BUSTED ^98^ (--branches Foreground --srv Yes), and relaxation or intensification of selection with RELAX ^99^ (CPU=1 --starting-points 100 --grid-size 2000 --models Minimal --srv Yes), for which the selection-intensity parameter K is reported (K > 1 indicating intensification and K < 1 relaxation). Each test was run under two foreground definitions: the single leading branch to the amplified copies (“Leading branch to the expansion”) and all branches of the expanded clade (“Full expansion”), with the number of foreground branches reported for the latter. In all cases the gorilla lineage/copies constituted the foreground, except for FRG1, where all modified copies were considered as foreground (this included the GGO copies along with the PTR and HSA copies).

## Resource Availability

### Lead contact

Adam M. Phillippy

### Materials availability

No new materials were generated from this study.

### Data and code availability

Supplementary data including custom analysis code, self-identity heatmaps for all ape chromosomes, multiple-sequence alignments (CDS, introns, SST1), and satellite annotations are available from doi: 10.5281/zenodo.18022857

Original data underlying the SST1 analyses present in this manuscript can be accessed from the Stowers Original Data Repository at https://www.stowers.org/research/publications/libpb-2603

### Supplemental information

Figures S1-S42. Tables S2–3,S5.

Table S1. Satellite bases and Pseudo-homolog regions

Table S4. Ape gene loci and consensus CDS.

## Acknowledgments

This work was supported, in part, by the Intramural Research Program of the US National Human Genome Research Institute, National Institutes of Health (S.J.S., A.S., A.R., J.K., B.D.P., M.A.W., S.K., and A.M.P.) and grants: R01CA266339 (J.L.G., T.P.); R50CA305001 (T.P.); R01HG002385, R01HG010169 (E.E.E.); R01HG014490, U01HG013748, U41HG010972, U24HG007234 (P.H., B.P.); and the National Recovery and Resilience Plan (NRRP), Mission 4, Component 2, Investment 1.1, Call for tender No. 104 published on 2 February 2022 by the Italian Ministry of University and Research (MUR), funded by the European Union—NextGenerationEU—Project Title “Telomere-to-telomere sequencing: the new era of Centromere and neocentromere eVolution (CenVolution)”, CUP H53D23003260006, grant assignment decree no. 1015 adopted on 7 July 2023 by the Italian MUR (M.V.). P.H. was supported, in part, by the Jack Baskin and Peggy Downes-Baskin Fellowship. E.E.E. is an investigator of the Howard Hughes Medical Institute. The contributions of NIH authors are considered Works of the United States Government. The findings and conclusions presented in this paper are those of the authors and do not necessarily reflect the views of the NIH or the U.S. Department of Health and Human Services. This work utilized the computational resources of the NIH HPC Biowulf cluster (http://hpc.nih.gov).

## Author Contributions

Conception: S.J.S., A.M.P. Initial draft: S.J.S., A.M.P. Writing/editing: S.J.S., P.H., L.G.L., T.P., L.G., A.G., M.A.W., E.E.E., M.V., J.L.G., A.M.P. Analysis: S.J.S., P.H., L.G.L., A.S., A.R., T.P., L.G., A.G., J.K., B.D.P., S.K., M.V., A.M.P. Figures: S.J.S., P.H., L.G.L., A.S., A.M.P. Supervision: B.P., E.G., E.E.E., M.V., J.L.G., A.M.P. Project coordination: A.M.P. All authors read and approved the final manuscript.

## References

1. Yunis, J.J., and Prakash, O. (1982). The origin of man: a chromosomal pictorial legacy. Science 215, 1525–1530. 10.1126/science.7063861.

2. Lichter, P., Cremer, T., Borden, J., Manuelidis, L., and Ward, D.C. (1988). Delineation of individual human chromosomes in metaphase and interphase cells by in situ suppression hybridization using recombinant DNA libraries. Hum Genet 80, 224–234. 10.1007/BF01790090.

3. Wienberg, J., Jauch, A., Stanyon, R., and Cremer, T. (1990). Molecular cytotaxonomy of primates by chromosomal in situ suppression hybridization. Genomics 8, 347–350. 10.1016/0888-7543(90)90292-3.

4. Wienberg, J. (2005). Fluorescence in situ hybridization to chromosomes as a tool to understand human and primate genome evolution. Cytogenet Genome Res 108, 139–160. 10.1159/000080811.

5. Ferguson-Smith, M.A., and Trifonov, V. (2007). Mammalian karyotype evolution. Nat Rev Genet 8, 950–962. 10.1038/nrg2199.

6. Lowry, D.B., and Willis, J.H. (2010). A widespread chromosomal inversion polymorphism contributes to a major life-history transition, local adaptation, and reproductive isolation. PLoS Biol 8, e1000500. 10.1371/journal.pbio.1000500.

7. IJdo, J.W., Baldini, A., Ward, D.C., Reeders, S.T., and Wells, R.A. (1991). Origin of human chromosome 2: an ancestral telomere-telomere fusion. Proc Natl Acad Sci U S A 88, 9051–9055. 10.1073/pnas.88.20.9051.

8. Miga, K.H. (2017). Chromosome-Specific Centromere Sequences Provide an Estimate of the Ancestral Chromosome 2 Fusion Event in Hominin Genomes. J Hered 108, 45–52. 10.1093/jhered/esw039.

9. Dutrillaux, B., Rethoré, M.O., Prieur, M., and Lejeune, J. (1973). [Analysis of the structure of chromatids of Gorilla gorilla. Comparison with Homo sapiens and Pan troglodytes (author’s transl)]. Humangenetik 20, 343–354. 10.1007/BF00273338.

10. Catacchio, C.R., Maggiolini, F.A.M., D’Addabbo, P., Bitonto, M., Capozzi, O., Lepore Signorile, M., Miroballo, M., Archidiacono, N., Eichler, E.E., Ventura, M., et al. (2018). Inversion variants in human and primate genomes. Genome Res 28, 910–920. 10.1101/gr.234831.118.

11. Yoo, D., Rhie, A., Hebbar, P., Antonacci, F., Logsdon, G.A., Solar, S.J., Antipov, D., Pickett, B.D., Safonova, Y., Montinaro, F., et al. (2025). Complete sequencing of ape genomes. Nature 641, 401–418. 10.1038/s41586-025-08816-3.

12. Nurk, S., Koren, S., Rhie, A., Rautiainen, M., Bzikadze, A.V., Mikheenko, A., Vollger, M.R., Altemose, N., Uralsky, L., Gershman, A., et al. (2022). The complete sequence of a human genome. Science 376, 44–53. 10.1126/science.abj6987.

13. Rhie, A., Nurk, S., Cechova, M., Hoyt, S.J., Taylor, D.J., Altemose, N., Hook, P.W., Koren, S., Rautiainen, M., Alexandrov, I.A., et al. (2023). The complete sequence of a human Y chromosome. Nature 621, 344–354. 10.1038/s41586-023-06457-y.

14. Hansen, N.F., Dwarshuis, N., Ji, H.J., Rhie, A., Loucks, H., Logsdon, G.A., Vollger, M.R., Storer, J.M., Kim, J., Adam, E., et al. (2026). A complete diploid human genome benchmark for personalized genomics. Cell 189, 4857–4875.e31. 10.1016/j.cell.2026.06.016.

15. Makova, K.D., Pickett, B.D., Harris, R.S., Hartley, G.A., Cechova, M., Pal, K., Nurk, S., Yoo, D., Li, Q., Hebbar, P., et al. (2024). The complete sequence and comparative analysis of ape sex chromosomes. Nature 630, 401–411. 10.1038/s41586-024-07473-2.

16. Jain, M., Olsen, H.E., Paten, B., and Akeson, M. (2016). The Oxford Nanopore MinION: delivery of nanopore sequencing to the genomics community. Genome Biol 17, 239. 10.1186/s13059-016-1103-0.

17. Wenger, A.M., Peluso, P., Rowell, W.J., Chang, P.-C., Hall, R.J., Concepcion, G.T., Ebler, J., Fungtammasan, A., Kolesnikov, A., Olson, N.D., et al. (2019). Accurate circular consensus long-read sequencing improves variant detection and assembly of a human genome. Nat Biotechnol 37, 1155–1162. 10.1038/s41587-019-0217-9.

18. Rautiainen, M., Nurk, S., Walenz, B.P., Logsdon, G.A., Porubsky, D., Rhie, A., Eichler, E.E., Phillippy, A.M., and Koren, S. (2023). Telomere-to-telomere assembly of diploid chromosomes with Verkko. Nat Biotechnol 41, 1474–1482. 10.1038/s41587-023-01662-6.

19. Vollger, M.R., Guitart, X., Dishuck, P.C., Mercuri, L., Harvey, W.T., Gershman, A., Diekhans, M., Sulovari, A., Munson, K.M., Lewis, A.P., et al. (2022). Segmental duplications and their variation in a complete human genome. Science 376, eabj6965. 10.1126/science.abj6965.

20. Bolognini, D., Halgren, A., Lou, R.N., Raveane, A., Rocha, J.L., Guarracino, A., Soranzo, N., Chin, C.-S., Garrison, E., and Sudmant, P.H. (2024). Recurrent evolution and selection shape structural diversity at the amylase locus. Nature 634, 617–625. 10.1038/s41586-024-07911-1.

21. de Lima, L.G., Guarracino, A., Koren, S., Potapova, T., McKinney, S., Rhie, A., Solar, S.J., Seidel, C., Fagen, B.L., Walenz, B.P., et al. (2025). The formation and propagation of human Robertsonian chromosomes. Nature 647, 952–961. 10.1038/s41586-025-09540-8.

22. Altemose, N., Logsdon, G.A., Bzikadze, A.V., Sidhwani, P., Langley, S.A., Caldas, G.V., Hoyt, S.J., Uralsky, L., Ryabov, F.D., Shew, C.J., et al. (2022). Complete genomic and epigenetic maps of human centromeres. Science 376, eabl4178. 10.1126/science.abl4178.

23. Hoyt, S.J., Storer, J.M., Hartley, G.A., Grady, P.G.S., Gershman, A., de Lima, L.G., Limouse, C., Halabian, R., Wojenski, L., Rodriguez, M., et al. (2022). From telomere to telomere: The transcriptional and epigenetic state of human repeat elements. Science 376, eabk3112. 10.1126/science.abk3112.

24. Logsdon, G.A., Rozanski, A.N., Ryabov, F., Potapova, T., Shepelev, V.A., Catacchio, C.R., Porubsky, D., Mao, Y., Yoo, D., Rautiainen, M., et al. (2024). The variation and evolution of complete human centromeres. Nature 629, 136–145. 10.1038/s41586-024-07278-3.

25. Antonarakis, S.E. (2022). Short arms of human acrocentric chromosomes and the completion of the human genome sequence. Genome Res 32, 599–607. 10.1101/gr.275350.121.

26. McStay, B. (2023). The p-Arms of Human Acrocentric Chromosomes Play by a Different Set of Rules. Annu Rev Genomics Hum Genet 24, 63–83. 10.1146/annurev-genom-101122-081642.

27. Levan, A., Fredga, K., and Sandberg, A.A. (1964). NOMENCLATURE FOR CENTROMERIC POSITION ON CHROMOSOMES. Hereditas 52, 201–220. 10.1111/j.1601-5223.1964.tb01953.x.

28. Lejeune, J., Levan, A., Böök, J.A., Chu, E.H.Y., Ford, C.E., Fraccaro, M., Harnden, D.G., Hsu, T.C., Hungerford, D.A., Jacobs, P.A., et al. (1960). A PROPOSED STANDARD SYSTEM OF NOMENCLATURE OF HUMAN MITOTIC CHROMOSOMES. The Lancet 275, 1063–1065. 10.1016/S0140-6736(60)90948-X.

29. Potapova, T.A., Kostos, P., McKinney, S., Borchers, M., Haug, J., Guarracino, A., Solar, S.J., Mattingly, M., Anez, G.M., de Lima, L.G., et al. (2025). Chromosome-specific epigenetic control and transmission of ribosomal DNA arrays in Hominidae genomes. Cell Genom 5, 101031. 10.1016/j.xgen.2025.101031.

30. Bobrow, M., Pearson, P.L., and Collacott, H.E. (1971). Para-nucleolar position of the human Y chromosome in interphase nuclei. Nature 232, 556–557. 10.1038/232556a0.

31. Subrt, I., and Blehová, B. (1974). Robertsonian translocation between the chromosome Y and 15. Humangenetik 23, 305–309. 10.1007/BF00272514.

32. Zimmer, E.A., Martin, S.L., Beverley, S.M., Kan, Y.W., and Wilson, A.C. (1980). Rapid duplication and loss of genes coding for the alpha chains of hemoglobin. Proc Natl Acad Sci U S A 77, 2158–2162. 10.1073/pnas.77.4.2158.

33. Brown, D.D., Wensink, P.C., and Jordan, E. (1972). A comparison of the ribosomal DNA’s of Xenopus laevis and Xenopus mulleri: the evolution of tandem genes. J Mol Biol 63, 57–73. 10.1016/0022-2836(72)90521-9.

34. Guarracino, A., Buonaiuto, S., de Lima, L.G., Potapova, T., Rhie, A., Koren, S., Rubinstein, B., Fischer, C., Human Pangenome Reference Consortium, Gerton, J.L., et al. (2023). Recombination between heterologous human acrocentric chromosomes. Nature 617, 335–343. 10.1038/s41586-023-05976-y.

35. Dover, G. (1982). Molecular drive: a cohesive mode of species evolution. Nature 299, 111–117. 10.1038/299111a0.

36. Gerton, J.L. (2024). A working model for the formation of Robertsonian chromosomes. J Cell Sci 137, jcs261912. 10.1242/jcs.261912.

37. Chiatante, G., Giannuzzi, G., Calabrese, F.M., Eichler, E.E., and Ventura, M. (2017). Centromere Destiny in Dicentric Chromosomes: New Insights from the Evolution of Human Chromosome 2 Ancestral Centromeric Region. Mol Biol Evol 34, 1669–1681. 10.1093/molbev/msx108.

38. Tantravahi, R., Miller, D.A., Dev, V.G., and Miller, O.J. (1976). Detection of nucleolus organizer regions in chromosomes of human, chimpanzee, gorilla, orangutan and gibbon. Chromosoma 56, 15–27. 10.1007/BF00293725.

39. Henderson, A.S., Atwood, K.C., and Warburton, D. (1976). Chromosomal distribution of rDNA in Pan paniscus, Gorilla gorilla beringei, and Symphalangus syndactylus: comparison to related primates. Chromosoma 59, 147–155. 10.1007/BF00328483.

40. Stanyon, R., Rocchi, M., Capozzi, O., Roberto, R., Misceo, D., Ventura, M., Cardone, M.F., Bigoni, F., and Archidiacono, N. (2008). Primate chromosome evolution: ancestral karyotypes, marker order and neocentromeres. Chromosome Res 16, 17–39. 10.1007/s10577-007-1209-z.

41. Carbone, L., Harris, R.A., Gnerre, S., Veeramah, K.R., Lorente-Galdos, B., Huddleston, J., Meyer, T.J., Herrero, J., Roos, C., Aken, B., et al. (2014). Gibbon genome and the fast karyotype evolution of small apes. Nature 513, 195–201. 10.1038/nature13679.

42. van Sluis, M., Gailín, M.Ó., McCarter, J.G.W., Mangan, H., Grob, A., and McStay, B. (2019). Human NORs, comprising rDNA arrays and functionally conserved distal elements, are located within dynamic chromosomal regions. Genes Dev 33, 1688–1701. 10.1101/gad.331892.119.

43. Tapparel, C., Reymond, A., Girardet, C., Guillou, L., Lyle, R., Lamon, C., Hutter, P., and Antonarakis, S.E. (2003). The TPTE gene family: cellular expression, subcellular localization and alternative splicing. Gene 323, 189–199. 10.1016/j.gene.2003.09.038.

44. Altemose, N., Miga, K.H., Maggioni, M., and Willard, H.F. (2014). Genomic characterization of large heterochromatic gaps in the human genome assembly. PLoS Comput Biol 10, e1003628. 10.1371/journal.pcbi.1003628.

45. Storer, J., Hubley, R., Rosen, J., Wheeler, T.J., and Smit, A.F. (2021). The Dfam community resource of transposable element families, sequence models, and genome annotations. Mob DNA 12, 2. 10.1186/s13100-020-00230-y.

46. Boots, J.L., von Pelchrzim, F., Weiss, A., Zimmermann, B., Friesacher, T., Radtke, M., Żywicki, M., Chen, D., Matylla-Kulińska, K., Zagrovic, B., et al. (2020). RNA polymerase II-binding aptamers in human ACRO1 satellites disrupt transcription in cis. Transcription 11, 217–229. 10.1080/21541264.2020.1790990.

47. Mayor, R., Izquierdo-Bouldstridge, A., Millán-Ariño, L., Bustillos, A., Sampaio, C., Luque, N., and Jordan, A. (2015). Genome distribution of replication-independent histone H1 variants shows H1.0 associated with nucleolar domains and H1X associated with RNA polymerase II-enriched regions. J Biol Chem 290, 7474–7491. 10.1074/jbc.M114.617324.

48. Yandım, C., and Karakülah, G. (2019). Expression dynamics of repetitive DNA in early human embryonic development. BMC Genomics 20, 439. 10.1186/s12864-019-5803-1.

49. Epstein, N.D., Karlsson, S., O’Brien, S., Modi, W., Moulton, A., and Nienhuis, A.W. (1987). A new moderately repetitive DNA sequence family of novel organization. Nucleic Acids Res 15, 2327–2341. 10.1093/nar/15.5.2327.

50. Dumbovic, G., Biayna, J., Banús, J., Samuelsson, J., Roth, A., Diederichs, S., Alonso, S., Buschbeck, M., Perucho, M., and Forcales, S.-V. (2018). A novel long non-coding RNA from NBL2 pericentromeric macrosatellite forms a perinucleolar aggregate structure in colon cancer. Nucleic Acids Res 46, 5504–5524. 10.1093/nar/gky263.

51. Yang, Z., Zhang, L., Jiang, X., Yang, X., Ma, K., Yoo, D., Lu, Y., Zhang, S., Chen, J., Nie, Y., et al. (2026). Incomplete lineage sorting of segmental duplications defines the human chromosome 2 fusion site early during African great ape speciation. Cell Genom 6, 101079. 10.1016/j.xgen.2025.101079.

52. Yoo, D., Munson, K.M., and Eichler, E.E. (2026). Epigenetic and evolutionary features of ape subterminal heterochromatin. Genome Res 36, 38–49. 10.1101/gr.280987.125.

53. Kosyakova, N., Grigorian, A., Liehr, T., Manvelyan, M., Simonyan, I., Mkrtchyan, H., Aroutiounian, R., Polityko, A.D., Kulpanovich, A.I., Egorova, T., et al. (2013). Heteromorphic variants of chromosome 9. Mol Cytogenet 6, 14. 10.1186/1755-8166-6-14.

54. Minocherhomji, S., Athalye, A.S., Madon, P.F., Kulkarni, D., Uttamchandani, S.A., and Parikh, F.R. (2009). A case-control study identifying chromosomal polymorphic variations as forms of epigenetic alterations associated with the infertility phenotype. Fertil Steril 92, 88–95. 10.1016/j.fertnstert.2008.05.071.

55. Hallast, P., Ebert, P., Loftus, M., Yilmaz, F., Audano, P.A., Logsdon, G.A., Bonder, M.J., Zhou, W., Höps, W., Kim, K., et al. (2023). Assembly of 43 human Y chromosomes reveals extensive complexity and variation. Nature 621, 355–364. 10.1038/s41586-023-06425-6.

56. Lin, J., Mastrorosa, F.K., Noyes, M.D., Yoo, D., Rhie, A., Porubsky, D., Hoekzema, K., Munson, K.M., Koundinya, N., Watkins, W.S., et al. (2026). Human acrocentric chromosome short-arm de novo mutation and recombination. Cell 189, 4876–4890.e8. 10.1016/j.cell.2026.05.035.

57. Hartley, G.A., Okhovat, M., O’Neill, R.J., and Carbone, L. (2021). Comparative Analyses of Gibbon Centromeres Reveal Dynamic Genus-Specific Shifts in Repeat Composition. Mol Biol Evol 38, 3972–3992. 10.1093/molbev/msab148.

58. Dumas, L., Kim, Y.H., Karimpour-Fard, A., Cox, M., Hopkins, J., Pollack, J.R., and Sikela, J.M. (2007). Gene copy number variation spanning 60 million years of human and primate evolution. Genome Res 17, 1266–1277. 10.1101/gr.6557307.

59. Ventura, M., Catacchio, C.R., Alkan, C., Marques-Bonet, T., Sajjadian, S., Graves, T.A., Hormozdiari, F., Navarro, A., Malig, M., Baker, C., et al. (2011). Gorilla genome structural variation reveals evolutionary parallelisms with chimpanzee. Genome Res 21, 1640–1649. 10.1101/gr.124461.111.

60. Sudmant, P.H., Huddleston, J., Catacchio, C.R., Malig, M., Hillier, L.W., Baker, C., Mohajeri, K., Kondova, I., Bontrop, R.E., Persengiev, S., et al. (2013). Evolution and diversity of copy number variation in the great ape lineage. Genome Res 23, 1373–1382. 10.1101/gr.158543.113.

61. Sun, C.-Y.J., van Koningsbruggen, S., Long, S.W., Straasheijm, K., Klooster, R., Jones, T.I., Bellini, M., Levesque, L., Brieher, W.M., van der Maarel, S.M., et al. (2011). Facioscapulohumeral muscular dystrophy region gene 1 is a dynamic RNA-associated and actin-bundling protein. J Mol Biol 411, 397–416. 10.1016/j.jmb.2011.06.014.

62. Jiang, S., Katz, T.A., Garee, J.P., DeMayo, F.J., Lee, A.V., and Oesterreich, S. (2015). Scaffold attachment factor B2 (SAFB2)-null mice reveal non-redundant functions of SAFB2 compared with its paralog, SAFB1. Dis Model Mech 8, 1121–1127. 10.1242/dmm.019885.

63. Narayana, J., and Horton, W.A. (2015). FGFR3 biology and skeletal disease. Connect Tissue Res 56, 427–433. 10.3109/03008207.2015.1051224.

64. Uhlén, M., Fagerberg, L., Hallström, B.M., Lindskog, C., Oksvold, P., Mardinoglu, A., Sivertsson, Å., Kampf, C., Sjöstedt, E., Asplund, A., et al. (2015). Proteomics. Tissue-based map of the human proteome. Science 347, 1260419. 10.1126/science.1260419.

65. Dishuck, P.C., Munson, K.M., Lewis, A.P., Dougherty, M.L., Underwood, J.G., Harvey, W.T., Hsieh, P., Pastinen, T., and Eichler, E.E. (2025). Structural variation, selection, and diversification of the NPIP gene family from the human pangenome. Cell Genom 5, 100977. 10.1016/j.xgen.2025.100977.

66. Kosakovsky Pond, S.L., Posada, D., Gravenor, M.B., Woelk, C.H., and Frost, S.D.W. (2006). Automated phylogenetic detection of recombination using a genetic algorithm. Mol Biol Evol 23, 1891–1901. 10.1093/molbev/msl051.

67. Kosakovsky Pond, S.L., Posada, D., Gravenor, M.B., Woelk, C.H., and Frost, S.D.W. (2006). GARD: a genetic algorithm for recombination detection. Bioinformatics 22, 3096–3098. 10.1093/bioinformatics/btl474.

68. Kosakovsky Pond, S.L., Poon, A.F.Y., Velazquez, R., Weaver, S., Hepler, N.L., Murrell, B., Shank, S.D., Magalis, B.R., Bouvier, D., Nekrutenko, A., et al. (2020). HyPhy 2.5-A Customizable Platform for Evolutionary Hypothesis Testing Using Phylogenies. Mol Biol Evol 37, 295–299. 10.1093/molbev/msz197.

69. Nei, M., and Rooney, A.P. (2005). Concerted and birth-and-death evolution of multigene families. Annu Rev Genet 39, 121–152. 10.1146/annurev.genet.39.073003.112240.

70. Babcock, M., Pavlicek, A., Spiteri, E., Kashork, C.D., Ioshikhes, I., Shaffer, L.G., Jurka, J., and Morrow, B.E. (2003). Shuffling of genes within low-copy repeats on 22q11 (LCR22) by Alu-mediated recombination events during evolution. Genome Res 13, 2519–2532. 10.1101/gr.1549503.

71. Imai, H.T., and Maruyama, T. (1978). Karyotype evolution by pericentric inversion as a stochastic process. J Theor Biol 70, 253–261. 10.1016/0022-5193(78)90375-2.

72. Pardo-Manuel de Villena, F., and Sapienza, C. (2001). Female meiosis drives karyotypic evolution in mammals. Genetics 159, 1179–1189. 10.1093/genetics/159.3.1179.

73. Henderson, A.S., Warburton, D., and Atwood, K.C. (1974). Localization of rDNA in the chromosome complement of the rhesus (Macaca mulatta). Chromosoma 44, 367–370. 10.1007/BF00284896.

74. Giannuzzi, G., Pazienza, M., Huddleston, J., Antonacci, F., Malig, M., Vives, L., Eichler, E.E., and Ventura, M. (2013). Hominoid fission of chromosome 14/15 and the role of segmental duplications. Genome Res 23, 1763–1773. 10.1101/gr.156240.113.

75. Hirai, H., Yamamoto, M.T., Ogura, K., Satta, Y., Yamada, M., Taylor, R.W., and Imai, H.T. (1994). Multiplication of 28S rDNA and NOR activity in chromosome evolution among ants of the Myrmecia pilosula species complex. Chromosoma 103, 171–178. 10.1007/BF00368009.

76. Dernburg, A.F., Sedat, J.W., and Hawley, R.S. (1996). Direct evidence of a role for heterochromatin in meiotic chromosome segregation. Cell 86, 135–146. 10.1016/s0092-8674(00)80084-7.

77. Karpen, G.H., Le, M.H., and Le, H. (1996). Centric heterochromatin and the efficiency of achiasmate disjunction in Drosophila female meiosis. Science 273, 118–122. 10.1126/science.273.5271.118.

78. Ohno, S. (1970). Evolution by Gene Duplication (Springer Berlin Heidelberg) 10.1007/978-3-642-86659-3.

79. Ohta, T., and Dover, G.A. (1983). Population genetics of multigene families that are dispersed into two or more chromosomes. Proc Natl Acad Sci U S A 80, 4079–4083. 10.1073/pnas.80.13.4079.

80. Pal, K., Greshnova, A., Park, S., Ko, B.J., Palova, H., Steinegger, M., Kosakovsky Pond, S.L., Canzar, S., and Makova, K.D. (2026). How and why multicopy genes survive on the human Y chromosome. Preprint at bioRxiv, 10.1101/2024.04.02.587783 https://doi.org/10.1101/2024.04.02.587783.

81. Hsu, L.Y., Benn, P.A., Tannenbaum, H.L., Perlis, T.E., and Carlson, A.D. (1987). Chromosomal polymorphisms of 1, 9, 16, and Y in 4 major ethnic groups: a large prenatal study. Am J Med Genet 26, 95–101. 10.1002/ajmg.1320260116.

82. Hansen, R.S., Wijmenga, C., Luo, P., Stanek, A.M., Canfield, T.K., Weemaes, C.M., and Gartler, S.M. (1999). The DNMT3B DNA methyltransferase gene is mutated in the ICF immunodeficiency syndrome. Proc Natl Acad Sci U S A 96, 14412–14417. 10.1073/pnas.96.25.14412.

83. Sweeten, A., Schatz, M.C., and Phillippy, A.M. (2026). AniAnn’s: alignment-free annotation of tandem repeat arrays using fast average nucleotide identity estimates. Bioinformatics 42, btag581. 10.1093/bioinformatics/btag581.

84. Sweeten, A.P., Schatz, M.C., and Phillippy, A.M. (2024). ModDotPlot-rapid and interactive visualization of tandem repeats. Bioinformatics 40, btae493. 10.1093/bioinformatics/btae493.

85. Li, H. (2018). Minimap2: pairwise alignment for nucleotide sequences. Bioinformatics 34, 3094–3100. 10.1093/bioinformatics/bty191.

86. Vollger, M.R. (2026). rustybam: a composable toolkit for alignment analysis and visualization with SafFire. Preprint at bioRxiv, 10.64898/2026.02.16.706142 https://doi.org/10.64898/2026.02.16.706142.

87. Porubsky, D., Guitart, X., Yoo, D., Dishuck, P.C., Harvey, W.T., and Eichler, E.E. (2025). SVbyEye: a visual tool to characterize structural variation among whole-genome assemblies. Bioinformatics 41, btaf332. 10.1093/bioinformatics/btaf332.

88. Kille, B., Garrison, E., Treangen, T.J., and Phillippy, A.M. (2023). Minmers are a generalization of minimizers that enable unbiased local Jaccard estimation. Bioinformatics 39, btad512. 10.1093/bioinformatics/btad512.

89. Hagberg, A.A., Schult, D.A., and Swart, P.J. (2008). Exploring Network Structure, Dynamics, and Function using NetworkX. In Proceedings of the 7th Python in Science Conference, G. Varoquaux, T. Vaught, and J. Millman, eds., pp. 11–15.

90. Kent, W.J. (2002). BLAT--the BLAST-like alignment tool. Genome Res 12, 656–664. 10.1101/gr.229202.

91. Prjibelski, A.D., Mikheenko, A., Joglekar, A., Smetanin, A., Jarroux, J., Lapidus, A.L., and Tilgner, H.U. (2023). Accurate isoform discovery with IsoQuant using long reads. Nat Biotechnol 41, 915–918. 10.1038/s41587-022-01565-y.

92. Danecek, P., Bonfield, J.K., Liddle, J., Marshall, J., Ohan, V., Pollard, M.O., Whitwham, A., Keane, T., McCarthy, S.A., Davies, R.M., et al. (2021). Twelve years of SAMtools and BCFtools. Gigascience 10, giab008. 10.1093/gigascience/giab008.

93. Jain, C., Rhie, A., Hansen, N.F., Koren, S., and Phillippy, A.M. (2022). Long-read mapping to repetitive reference sequences using Winnowmap2. Nat Methods 19, 705–710. 10.1038/s41592-022-01457-8.

94. Rhie, A., Walenz, B.P., Koren, S., and Phillippy, A.M. (2020). Merqury: reference-free quality, completeness, and phasing assessment for genome assemblies. Genome Biol 21, 245. 10.1186/s13059-020-02134-9.

95. Katoh, K., and Standley, D.M. (2013). MAFFT multiple sequence alignment software version 7: improvements in performance and usability. Mol Biol Evol 30, 772–780. 10.1093/molbev/mst010.

96. Kozlov, A.M., Darriba, D., Flouri, T., Morel, B., and Stamatakis, A. (2019). RAxML-NG: a fast, scalable and user-friendly tool for maximum likelihood phylogenetic inference. Bioinformatics 35, 4453–4455. 10.1093/bioinformatics/btz305.

97. Kumar, S., Stecher, G., Suleski, M., Sanderford, M., Sharma, S., and Tamura, K. (2024). MEGA12: Molecular Evolutionary Genetic Analysis Version 12 for Adaptive and Green Computing. Mol Biol Evol 41, msae263. 10.1093/molbev/msae263.

98. Murrell, B., Weaver, S., Smith, M.D., Wertheim, J.O., Murrell, S., Aylward, A., Eren, K., Pollner, T., Martin, D.P., Smith, D.M., et al. (2015). Gene-wide identification of episodic selection. Mol Biol Evol 32, 1365–1371. 10.1093/molbev/msv035.

99. Wertheim, J.O., Murrell, B., Smith, M.D., Kosakovsky Pond, S.L., and Scheffler, K. (2015). RELAX: detecting relaxed selection in a phylogenetic framework. Mol Biol Evol 32, 820–832. 10.1093/molbev/msu400.

100. Kosakovsky Pond, S.L., and Frost, S.D.W. (2005). Not so different after all: a comparison of methods for detecting amino acid sites under selection. Mol Biol Evol 22, 1208–1222. 10.1093/molbev/msi105.

101. Minh, B.Q., Schmidt, H.A., Chernomor, O., Schrempf, D., Woodhams, M.D., von Haeseler, A., and Lanfear, R. (2020). IQ-TREE 2: New Models and Efficient Methods for Phylogenetic Inference in the Genomic Era. Mol Biol Evol 37, 1530–1534. 10.1093/molbev/msaa015.

102. Smit, A.F.A., Hubley, R., and Green, P. (2013). RepeatMasker Open-4.0. http://www.repeatmasker.org.

103. Junier, T., and Pagni, M. (2000). Dotlet: diagonal plots in a web browser. Bioinformatics 16, 178–179. 10.1093/bioinformatics/16.2.178.

104. Darriba, D., Taboada, G.L., Doallo, R., and Posada, D. (2012). jModelTest 2: more models, new heuristics and parallel computing. Nat Methods 9, 772. 10.1038/nmeth.2109.

105. Schindelin, J., Arganda-Carreras, I., Frise, E., Kaynig, V., Longair, M., Pietzsch, T., Preibisch, S., Rueden, C., Saalfeld, S., Schmid, B., et al. (2012). Fiji: an open-source platform for biological-image analysis. Nat Methods 9, 676–682. 10.1038/nmeth.2019.

106. Kim, J.-H., Dilthey, A.T., Nagaraja, R., Lee, H.-S., Koren, S., Dudekula, D., Wood Iii, W.H., Piao, Y., Ogurtsov, A.Y., Utani, K., et al. (2018). Variation in human chromosome 21 ribosomal RNA genes characterized by TAR cloning and long-read sequencing. Nucleic Acids Res 46, 6712–6725. 10.1093/nar/gky442.

107. Perez, G., Barber, G.P., Benet-Pages, A., Casper, J., Clawson, H., Diekhans, M., Fischer, C., Gonzalez, J.N., Hinrichs, A.S., Lee, C.M., et al. (2025). The UCSC Genome Browser database: 2025 update. Nucleic Acids Res 53, D1243–D1249. 10.1093/nar/gkae974.

108. Goldfarb, T., Kodali, V.K., Pujar, S., Brover, V., Robbertse, B., Farrell, C.M., Oh, D.-H., Astashyn, A., Ermolaeva, O., Haddad, D., et al. (2025). NCBI RefSeq: reference sequence standards through 25 years of curation and annotation. Nucleic Acids Res 53, D243–D257. 10.1093/nar/gkae1038.

109. Hoang, D.T., Chernomor, O., von Haeseler, A., Minh, B.Q., and Vinh, L.S. (2018). UFBoot2: Improving the Ultrafast Bootstrap Approximation. Mol Biol Evol 35, 518–522. 10.1093/molbev/msx281.

110. To, T.-H., Jung, M., Lycett, S., and Gascuel, O. (2016). Fast Dating Using Least-Squares Criteria and Algorithms. Syst Biol 65, 82–97. 10.1093/sysbio/syv068.

111. Ventura, M., Mudge, J.M., Palumbo, V., Burn, S., Blennow, E., Pierluigi, M., Giorda, R., Zuffardi, O., Archidiacono, N., Jackson, M.S., et al. (2003). Neocentromeres in 15q24-26 map to duplicons which flanked an ancestral centromere in 15q25. Genome Res 13, 2059–2068. 10.1101/gr.1155103.

