## Supplementary Figures for "Origin and evolution of acrocentric chromosomes in human and great apes"

### Supplemental Materials for
### Origin and evolution of acrocentric chromosomes in human and great apes

#### Table of Contents

#### Supplemental Figures S1-S42

S1. Self-alignment heatmaps for the short arms of the S. orangutan acrocentrics.
S2. Self-alignment heatmaps for the short arms of the B. orangutan acrocentrics.
S3. Self-alignment heatmaps for the short arms of the human (HG002) acrocentrics.
S4. Self-alignment heatmaps for the short arms of the chimp acrocentrics.
S5. Self-alignment heatmaps for the short arms of the bonobo acrocentrics.
S6. Self-alignment heatmaps for the short arms of the gorilla acrocentrics.
S7. Self-alignment heatmaps for the short arms of the siamang acrocentrics.
S8. Location of 45S rRNA pseudogenes in the pericentromeres of HSA16.
S9. Acrocentric-like satellites per chromosome for each of the great apes.
S10. Whole-chromosome pairwise alignments for HSA2A.
S11. Whole-chromosome pairwise alignments for HSA2B.
S12. Whole-chromosome pairwise alignments for HSA9.
S13. Whole-chromosome pairwise alignments for HSA13.
S14. Whole-chromosome pairwise alignments for HSA14.
S15. Whole-chromosome pairwise alignments for HSA15.
S16. Whole-chromosome pairwise alignments for HSA18.
S17. Whole-chromosome pairwise alignments for HSA21.
S18. Whole-chromosome pairwise alignments for HSA22.
S19. Whole-chromosome pairwise alignments for HSAY.
S20. Location of 45S rDNA fragments in the pericentromere of human HG002 ChrY.
S21. Amplification of a 45S rDNA DJ-like sequence alongside TSPY and RBMY on ChrY of chimp and bonobo.
S22. Localization of amplified gene regions and rDNA in representative interphase fibroblast nuclei of four different gorillas.
S23. Alignment of BAC RP11-341D18 to the gorilla genome.
S24. Metaphase FISH of the *GGT* amplification in gorilla.
S25. Chromosome-labeled FISH of the *GGT* amplification in gorilla.
S26. Alignment of BAC RP11-529E10 to the gorilla genome.
S27. Chromosome-labeled FISH of the *LRPAP1* (*FRG1*) amplification in gorilla.
S28. Methylation pattern and transcript structure across the ancestral, duplicated full copies, and modified *FRG1* copies
S29. CDS tree for *ABHD17A*.
S30. CDS tree for *BAGE2*.

S31. CDS tree for *C1QTNF3*.
S32. CDS tree for *CES4A*.
S33. CDS tree for *FGFR3*.
S34. CDS tree for *FRG1*.
S35. CDS tree for *GGT*.
S36. CDS tree for *IGSF3*.
S37. CDS tree for *LETM1*.
S38. CDS tree for *LRPAP1*.
S39. CDS tree for *SAFB2*.
S40. CDS tree for *TPTE*.
S41. Time-measured intron phylogeny for *FRG1*.
S42. Time-measured intron phylogeny for *GGT*.

#### Supplemental Tables S2,3,5

Table S2. Gorilla amplified gene CPMs
Table S3. Gorilla GGT FISH Summary
Table S5. Selection Testing

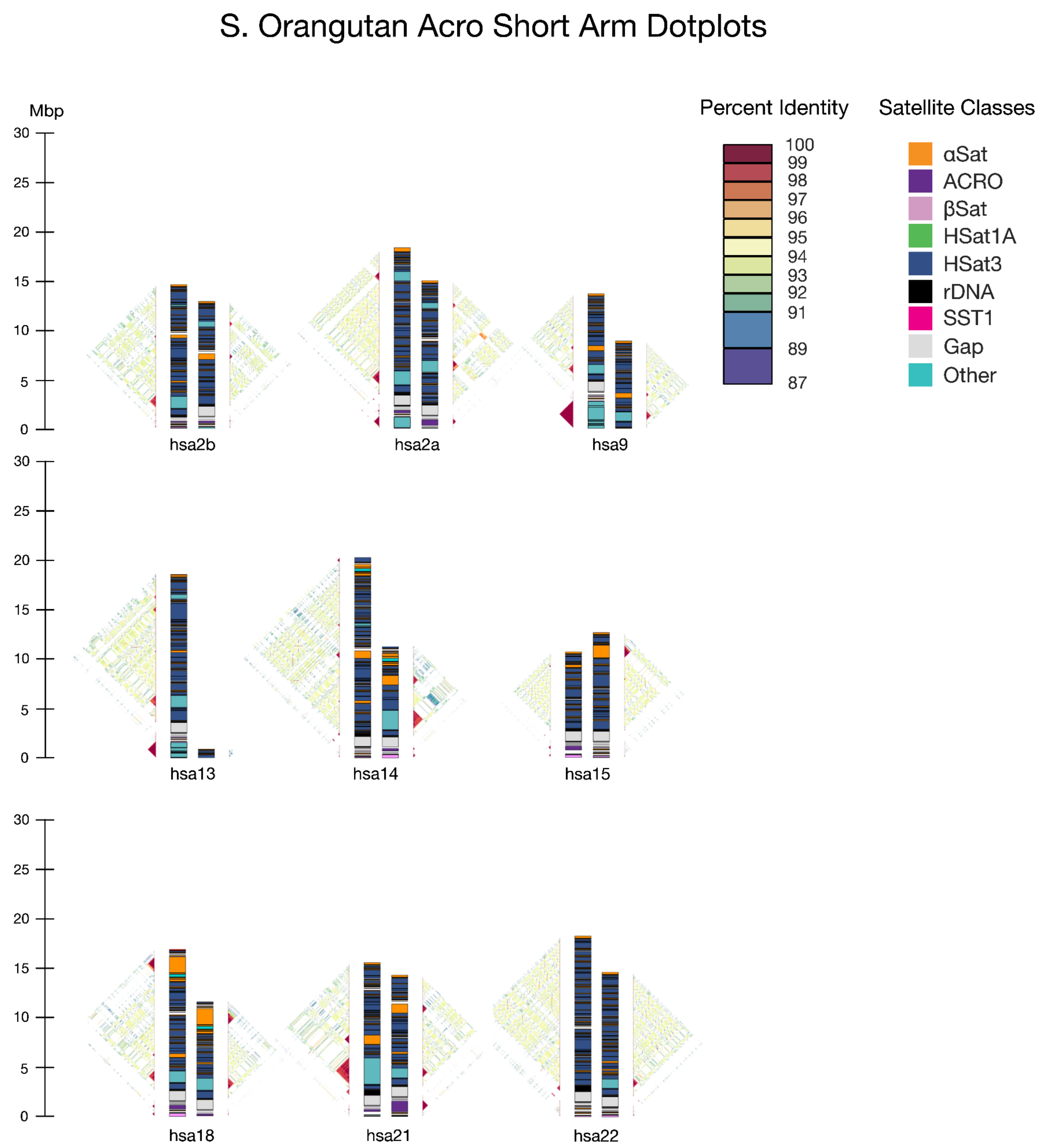


**Figure S1. Self-alignment heatmaps for the short arms of the S. orangutan acrocentrics.** ModDotPlot figures illustrate the differences in size and content between primary (left) and alternate (right) haplotypes. Corresponding satellite annotation tracks are provided in between each plot.

#### B. Orangutan Acro Short Arm Dotplots

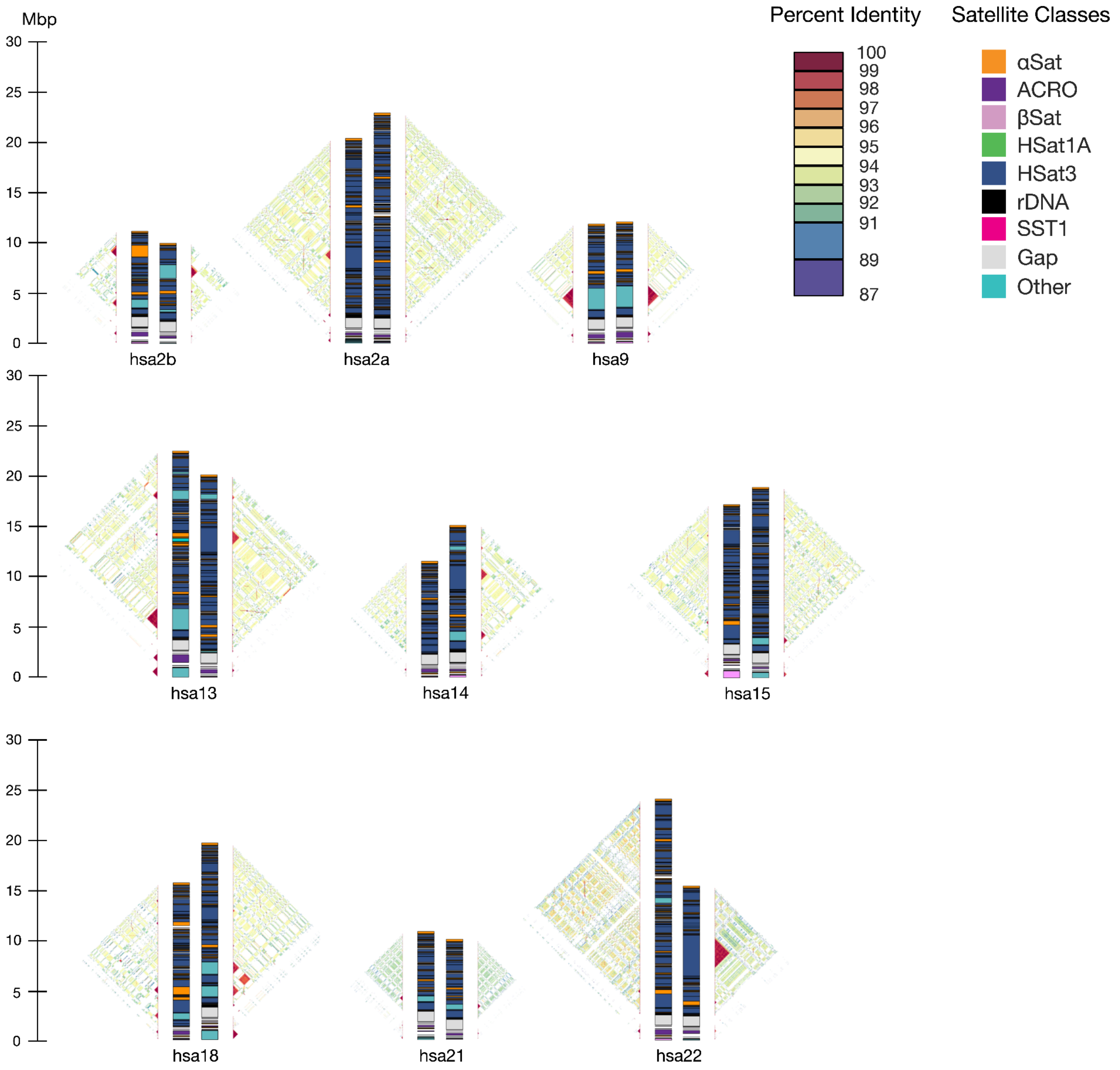


**Figure S2. Self-alignment heatmaps for the short arms of the B. orangutan acrocentrics.** ModDotPlot figures illustrate the differences in size and content between primary (left) and alternate (right) haplotypes. Corresponding satellite annotation tracks are provided in between each plot.

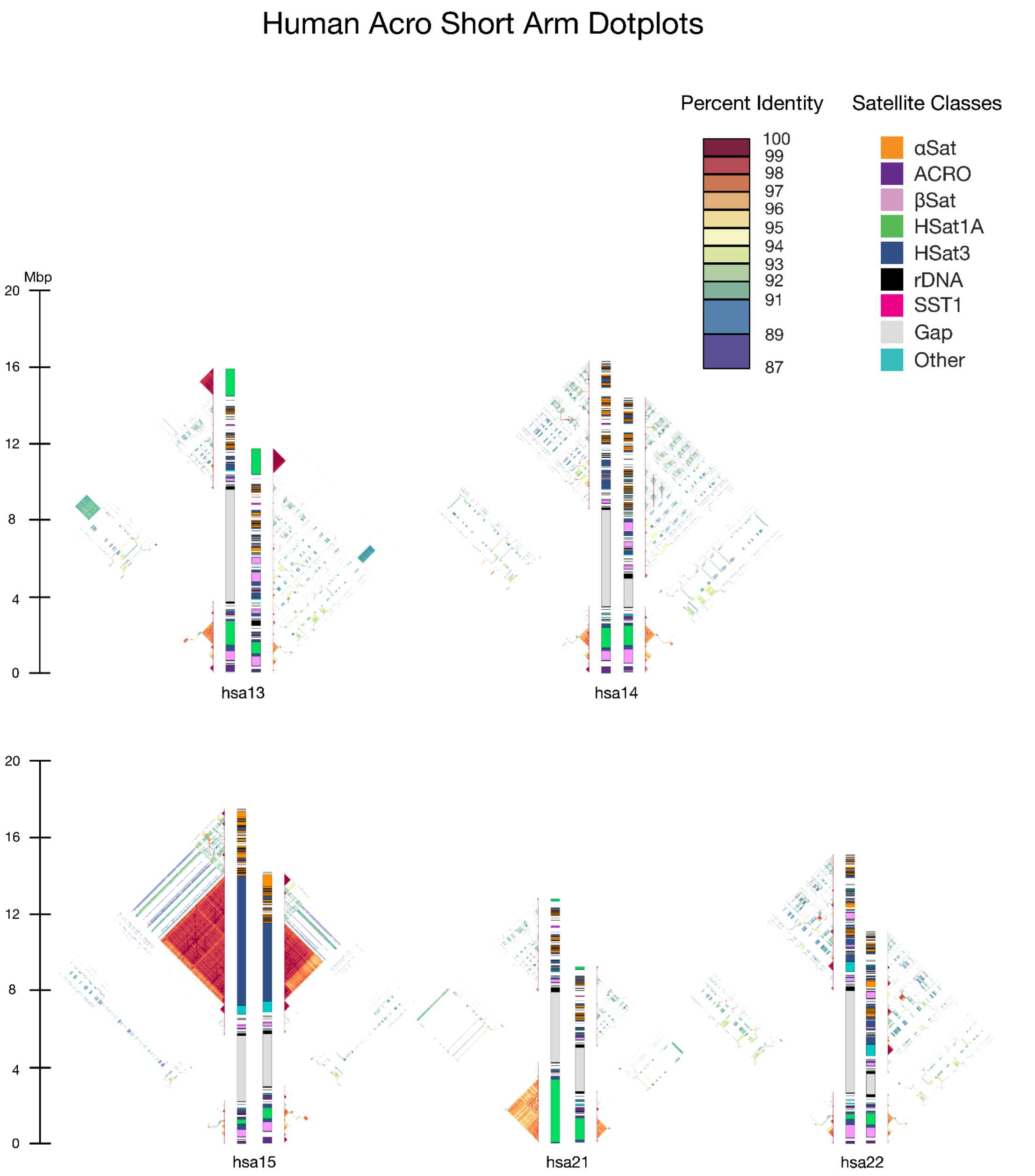


**Figure S3. Self-alignment heatmaps for the short arms of the human (HG002) acrocentrics.** ModDotPlot figures illustrate the differences in size and content between maternal (left) and paternal (right) haplotypes. Corresponding satellite annotation tracks are provided in between each plot.

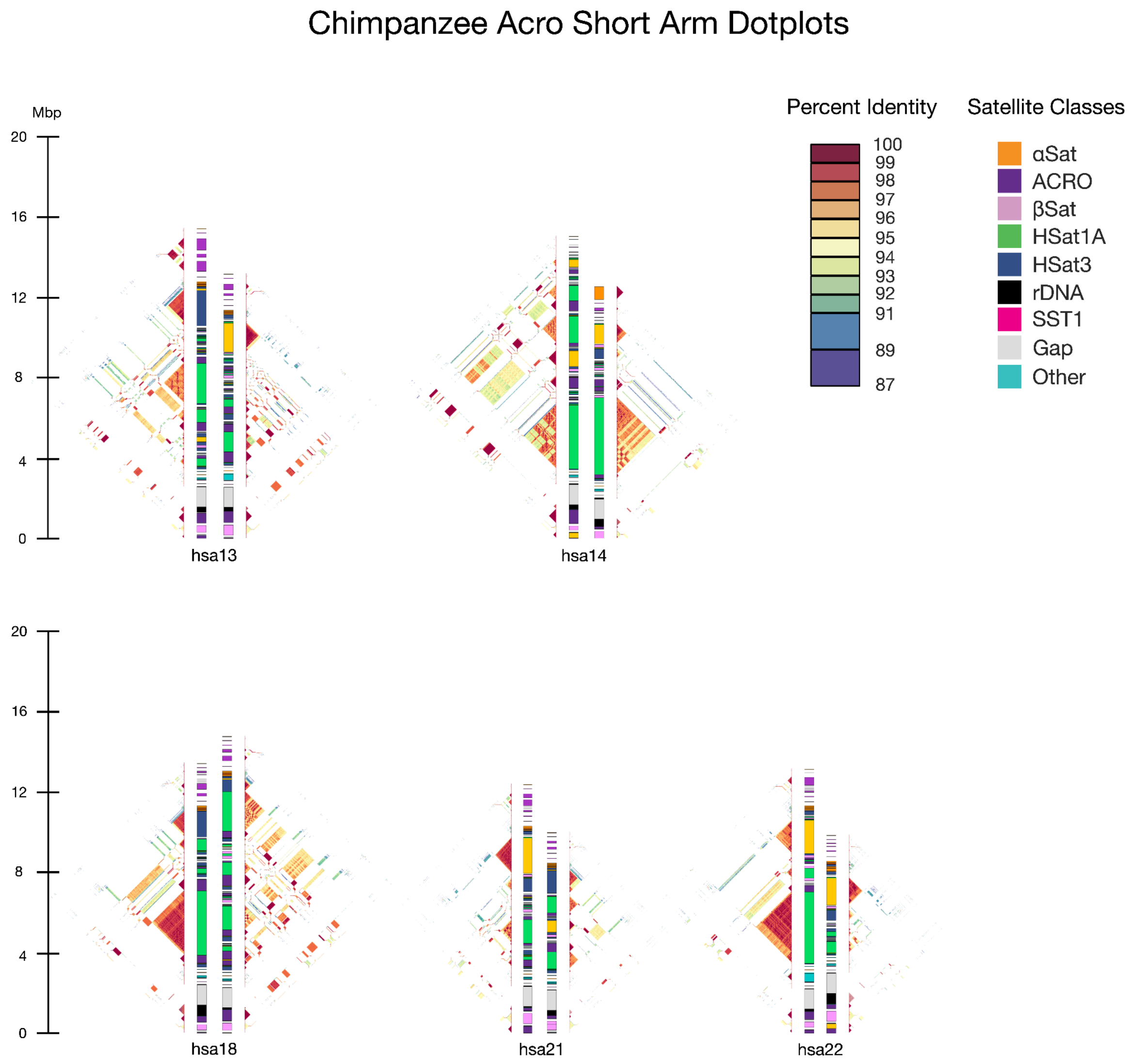


**Figure S4. Self-alignment heatmaps for the short arms of the chimp acrocentrics.** ModDotPlot figures illustrate the differences in size and content between primary (left) and alternate (right) haplotypes. Corresponding satellite annotation tracks are provided in between each plot.

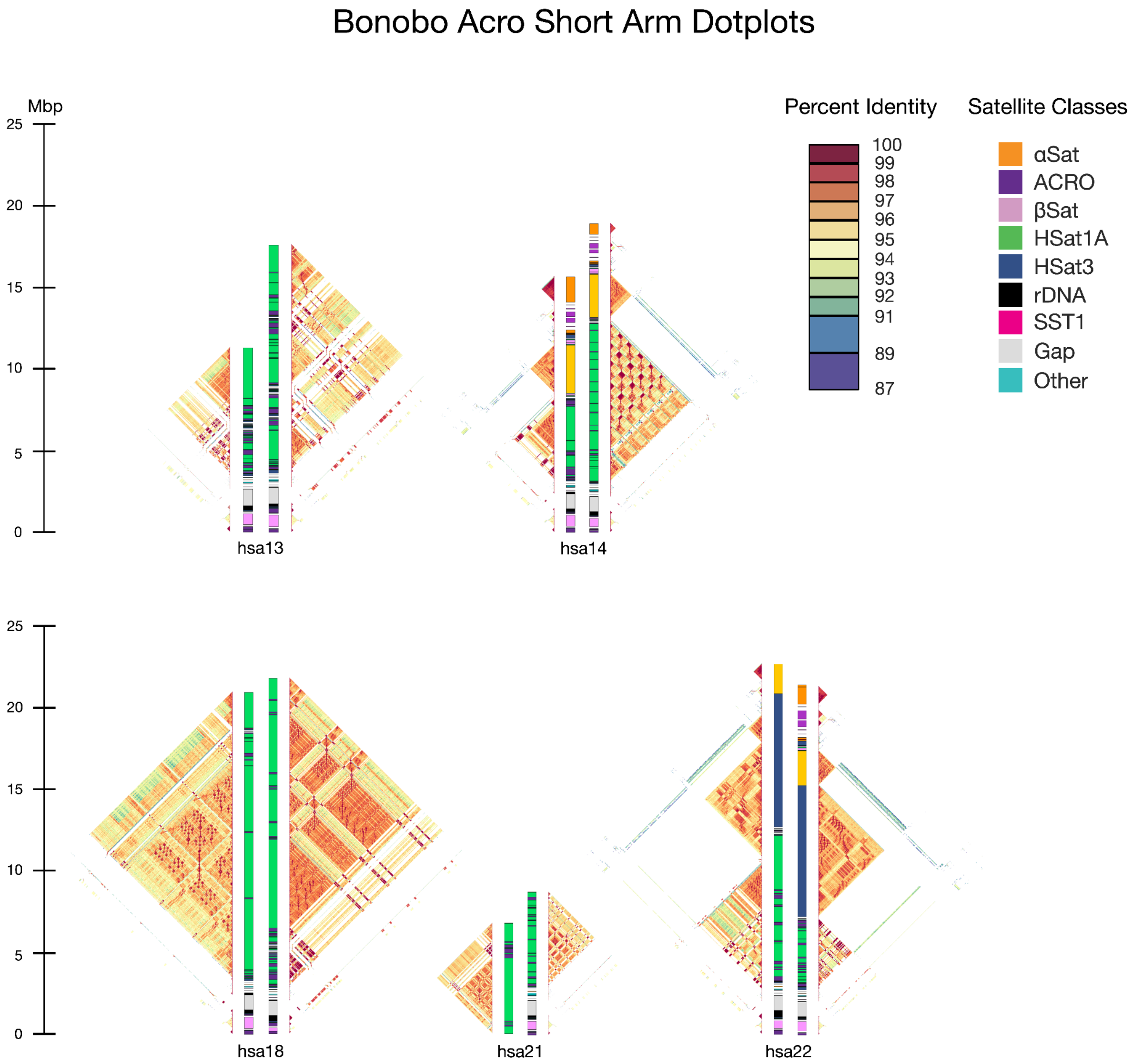


**Figure S5. Self-alignment heatmaps for the short arms of the bonobo acrocentrics.** ModDotPlot figures illustrate the differences in size and content between maternal (left) and paternal (right) haplotypes. Corresponding satellite annotation tracks are provided in between each plot.

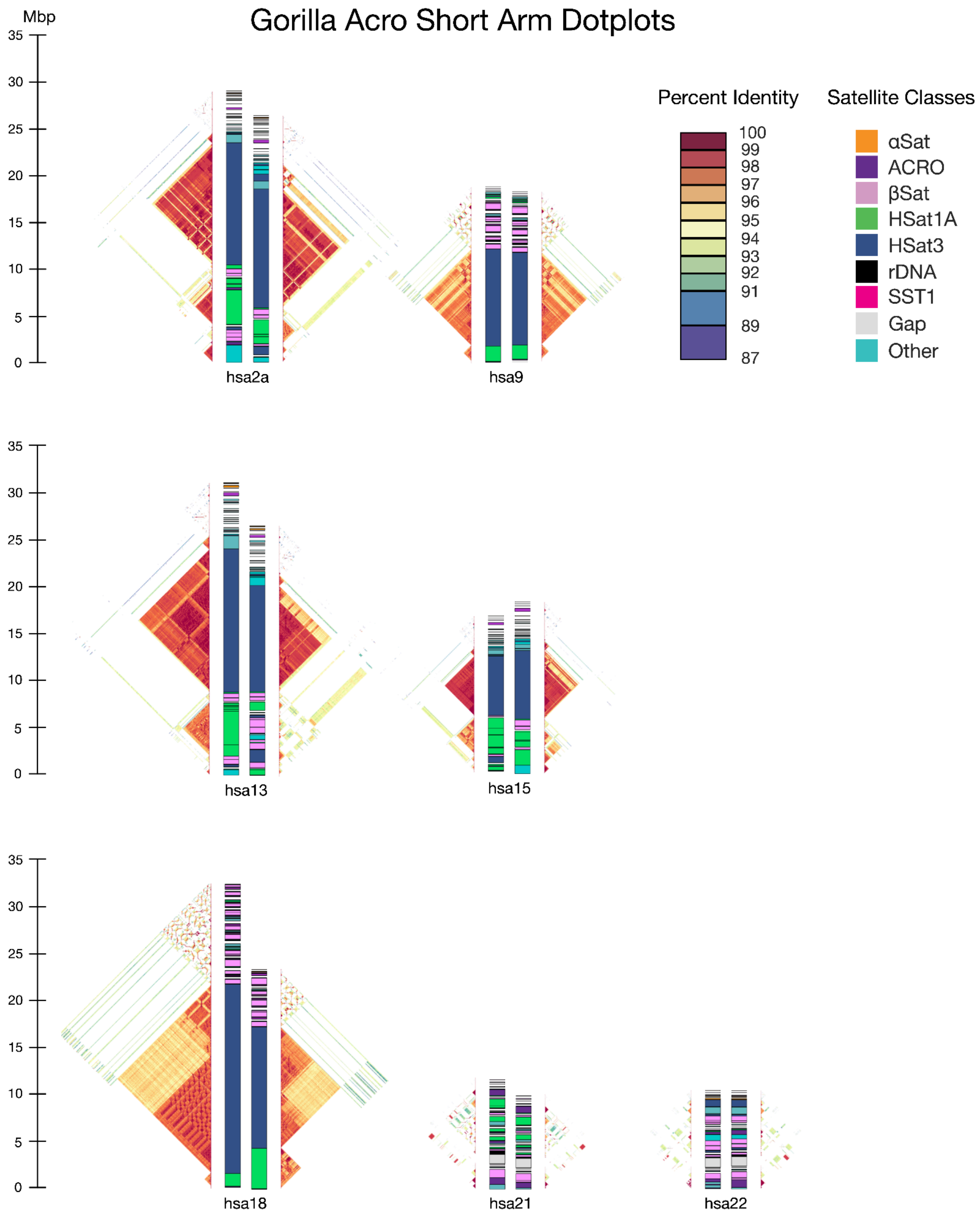


**Figure S6. Self-alignment heatmaps for the short arms of the gorilla acrocentrics.** ModDotPlot figures illustrate the differences in size and content between maternal (left) and paternal (right) haplotypes. Corresponding satellite annotation tracks are provided in between each plot.

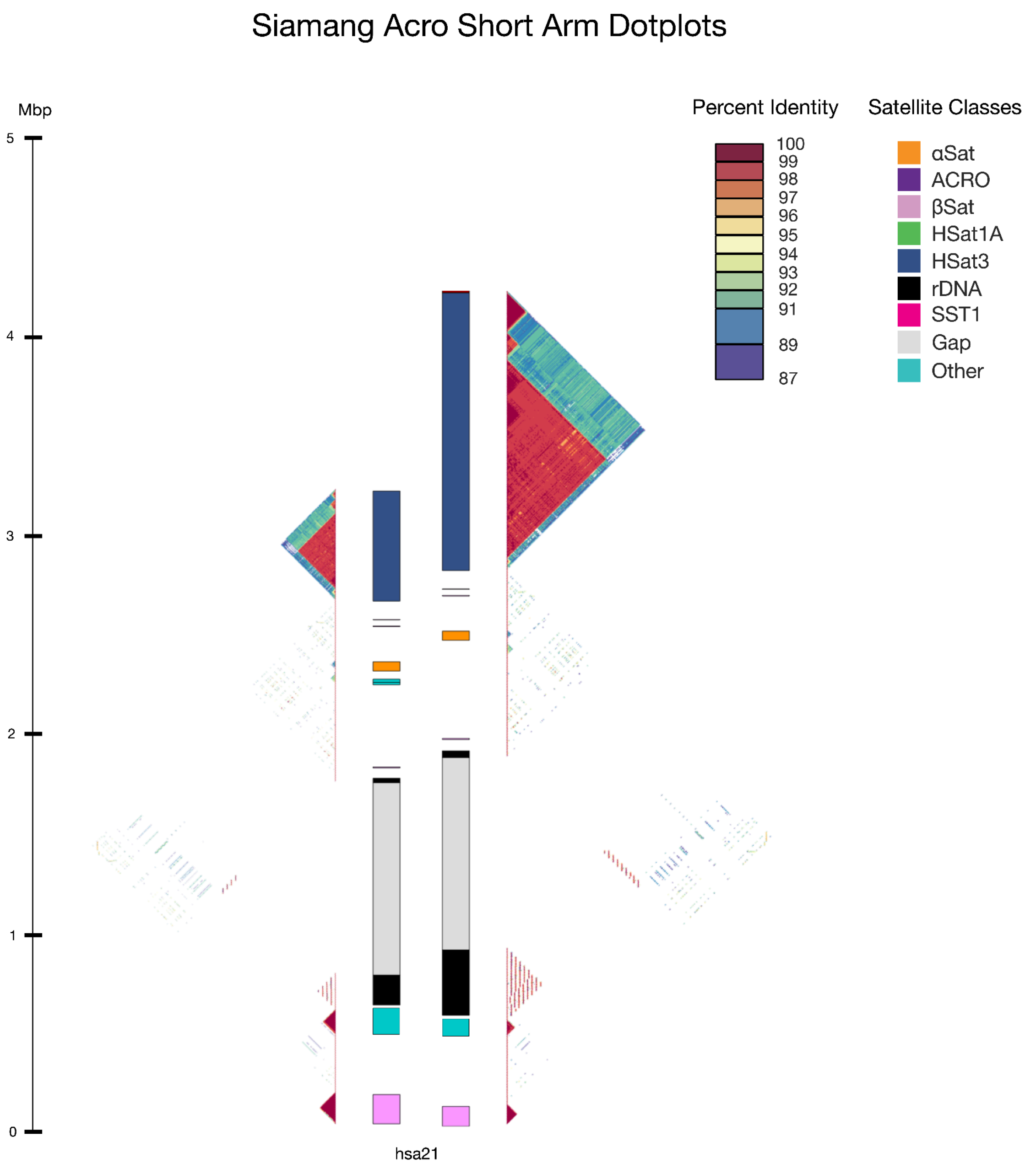


**Figure S7. Self-alignment heatmaps for the short arms of the siamang acrocentrics.** ModDotPlot figures illustrate the differences in size and content between primary (left) and alternate (right) haplotypes. Corresponding satellite annotation tracks are provided in between each plot.

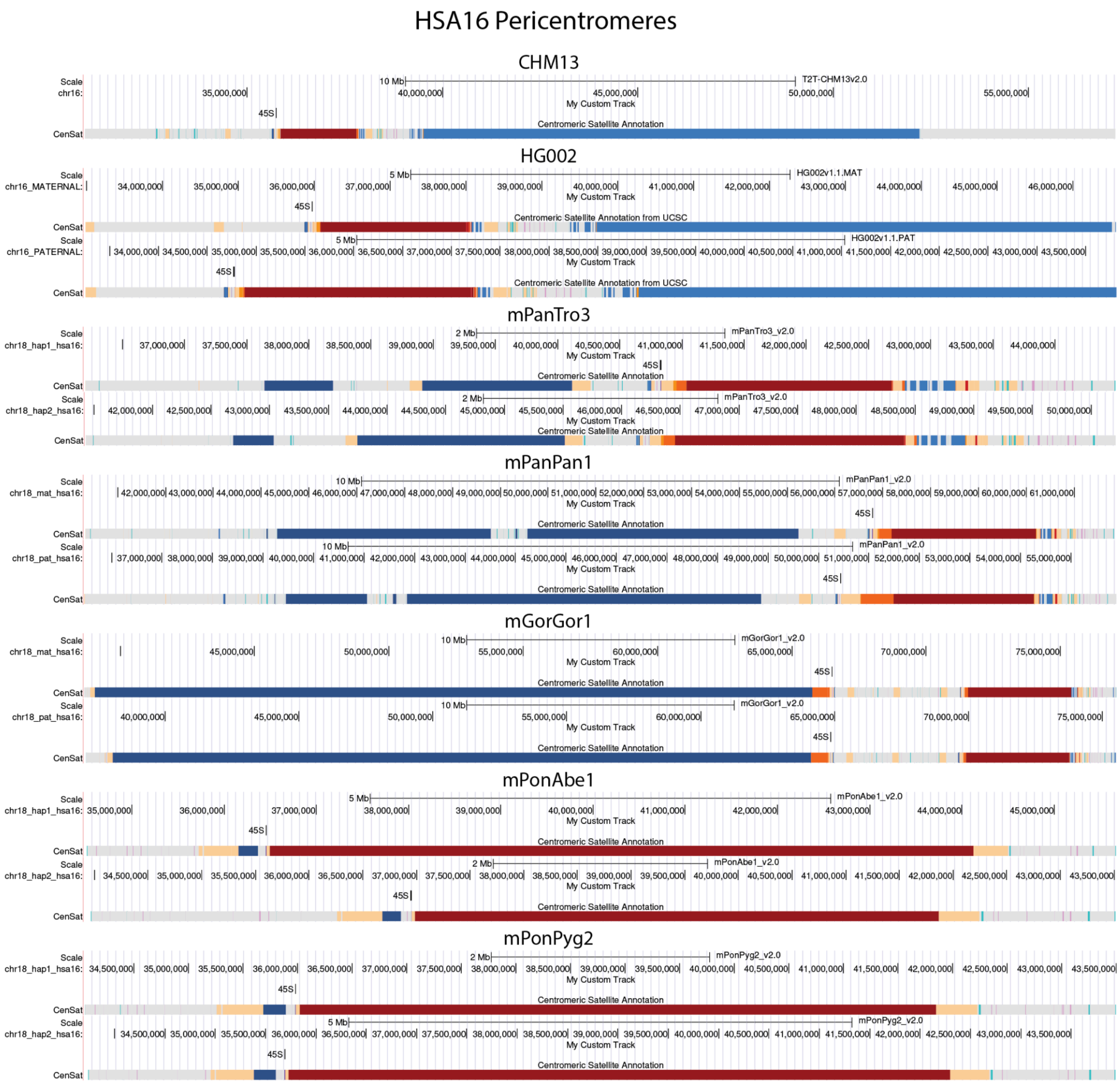


**Figure S8. Location of 45S rRNA pseudogenes in the pericentromeres of HSA16.** Both haplotypes of all great ape assemblies are shown. Satellite annotation tracks use the color code from the UCSC Genome Browser, with HSat2,3 as blue and the active, centromeric αSat as red. The 45S rRNA pseudogenes are labeled “45S” and marked above each satellite annotation. They are directly adjacent to the active αSat array in all species, except gorilla where they are slightly more distal and adjacent to a large HSat3 array.

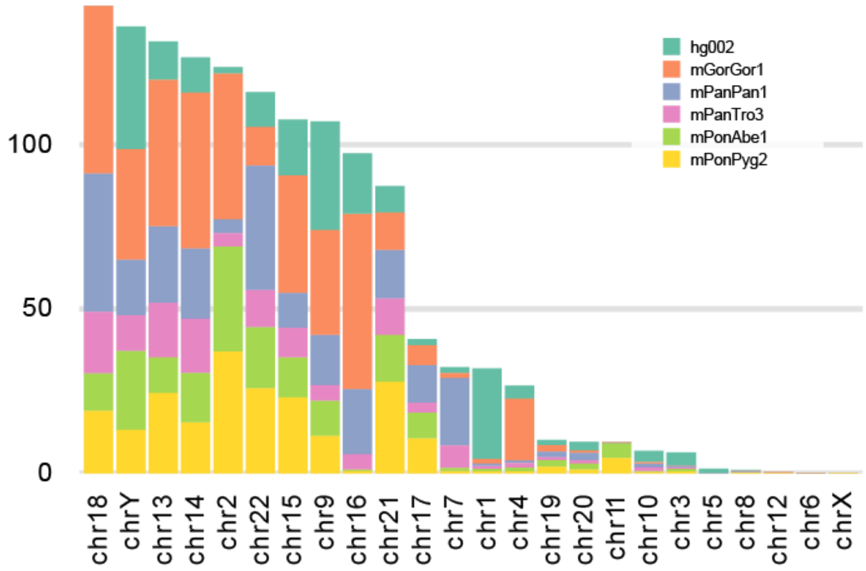


**Figure S9. Acrocentric-like satellites per chromosome for each of the great apes.** Total bases of acrocentric-associated satellites HSat2,3, HSat1A, βSat, ACRO, SST1, and CER, summed over each homologous human chromosome in the great apes and stacked by species. Most species have a similar amount of satellite DNA per chromosome, with notable exceptions of HSA1, HSA4, and HSA7, for which the sums are dominated by species-specific satellite expansions in human, gorilla, and chimp/bonobo, respectively.

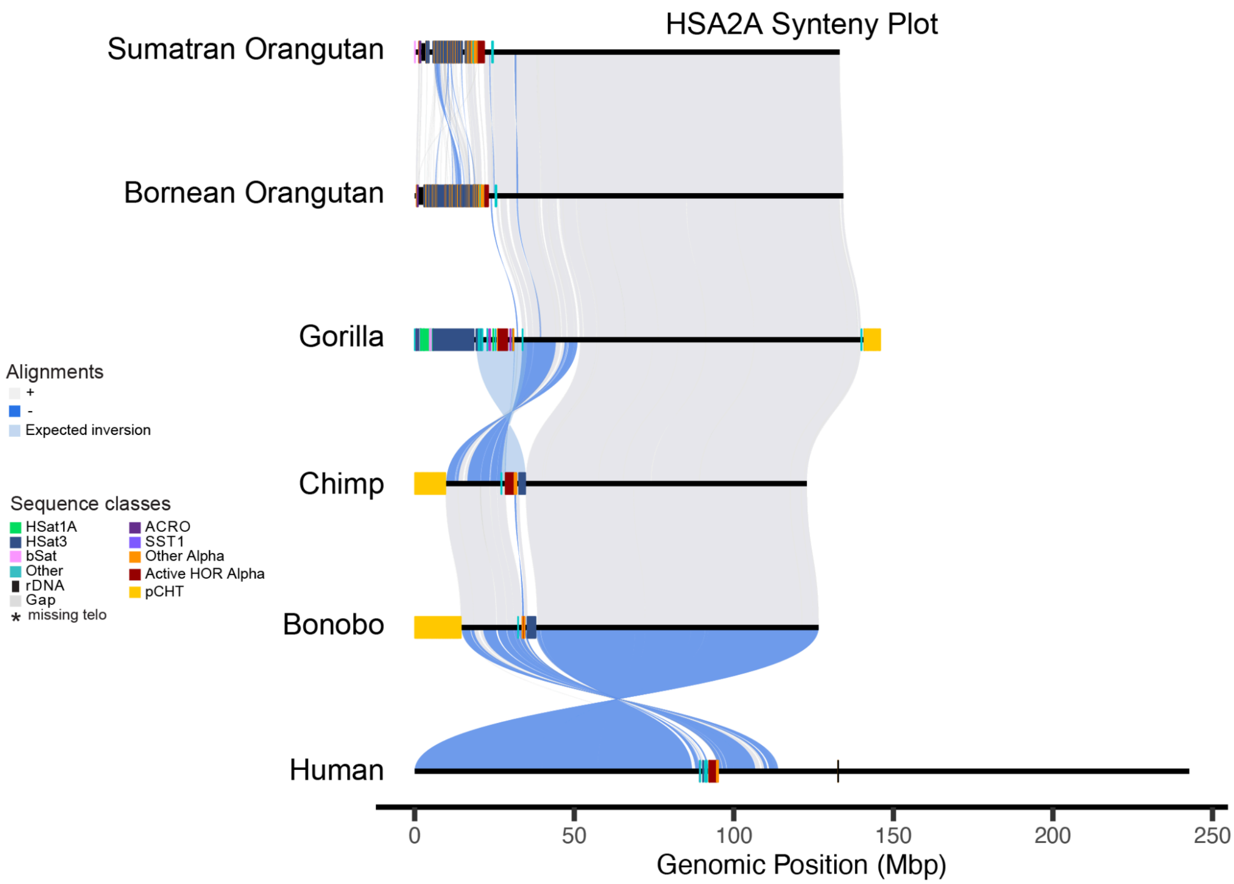


**Figure S10. Whole-chromosome pairwise alignments for HSA2A.** SVbyEye visualization of wfmash alignments between pairs of great ape chromosomes. Forward alignments are in grey and reverse complement in blue. Satellite annotations are given along each genome's axis with the centromeres marked in red. The whole p-arm inversion between gorilla and chimp is shown. The inversion between bonobo and human is artifactual and simply reflects the opposite orientation of the larger human fusion chromosome Chr2.

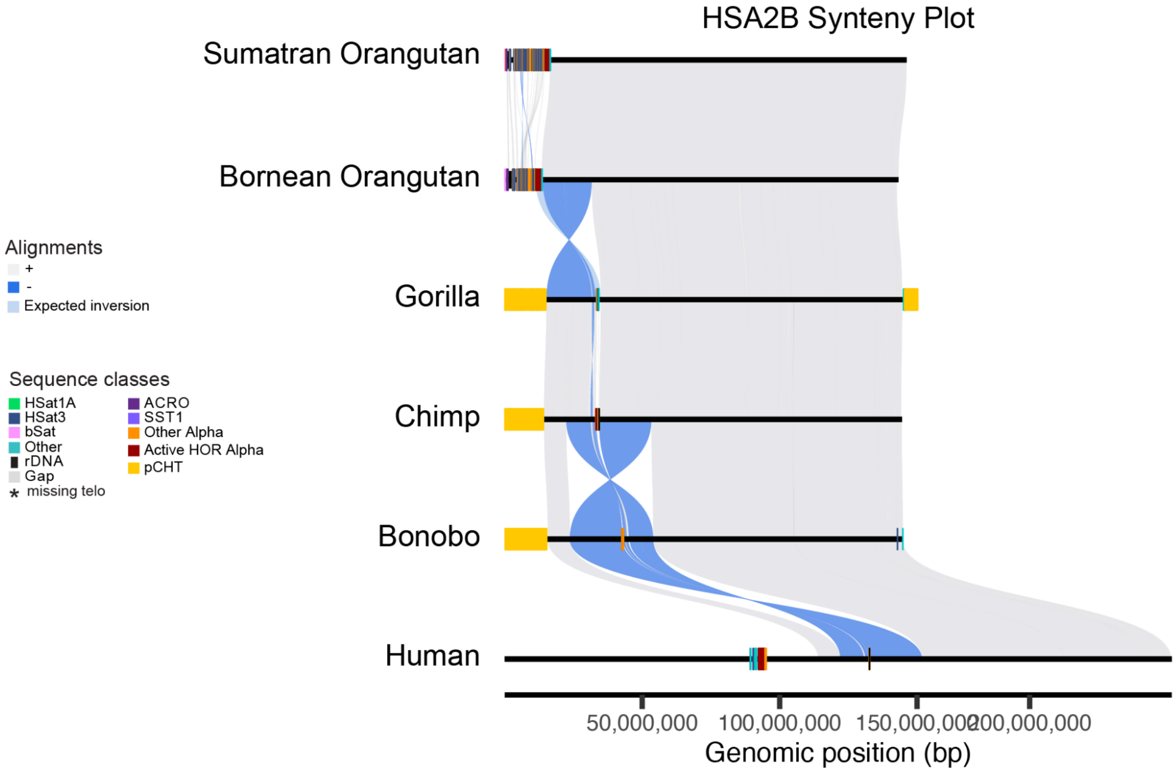


**Figure S11. Whole-chromosome pairwise alignments for HSA2B.** SVbyEye visualization of wfmash alignments between pairs of great ape chromosomes. Forward alignments are in grey and reverse complement in blue. Satellite annotations are given along each genome's axis with the centromeres marked in red. The whole p-arm inversion between orangutan and gorilla is shown, followed by a further paracentric inversion in humans, likely preceding the 2A-2B fusion.

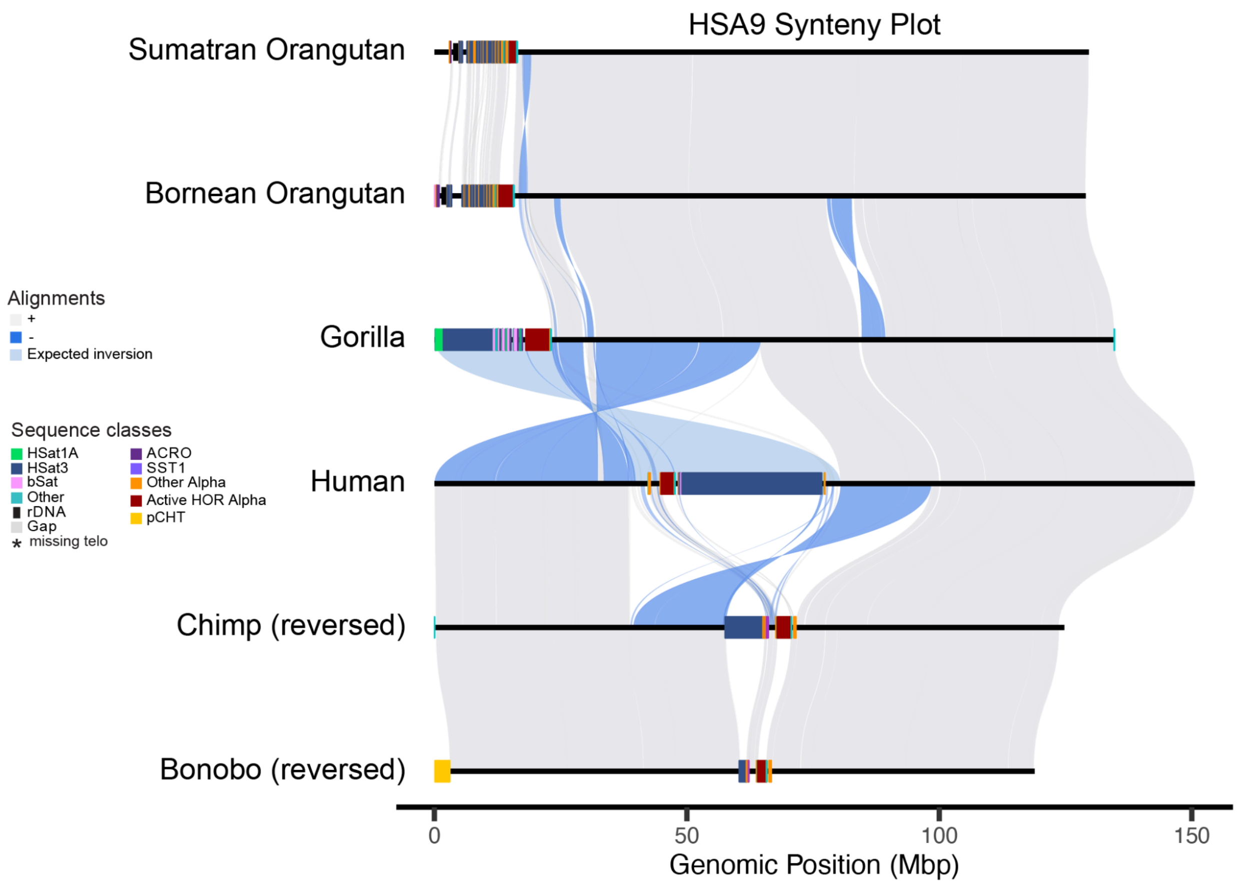


**Figure S12. Whole-chromosome pairwise alignments for HSA9.** SVbyEye visualization of wfmash alignments between pairs of great ape chromosomes. Forward alignments are in grey and reverse complement in blue. Satellite annotations are given along each genome's axis with the centromeres marked in red. The whole p-arm inversion between gorilla and human is shown.

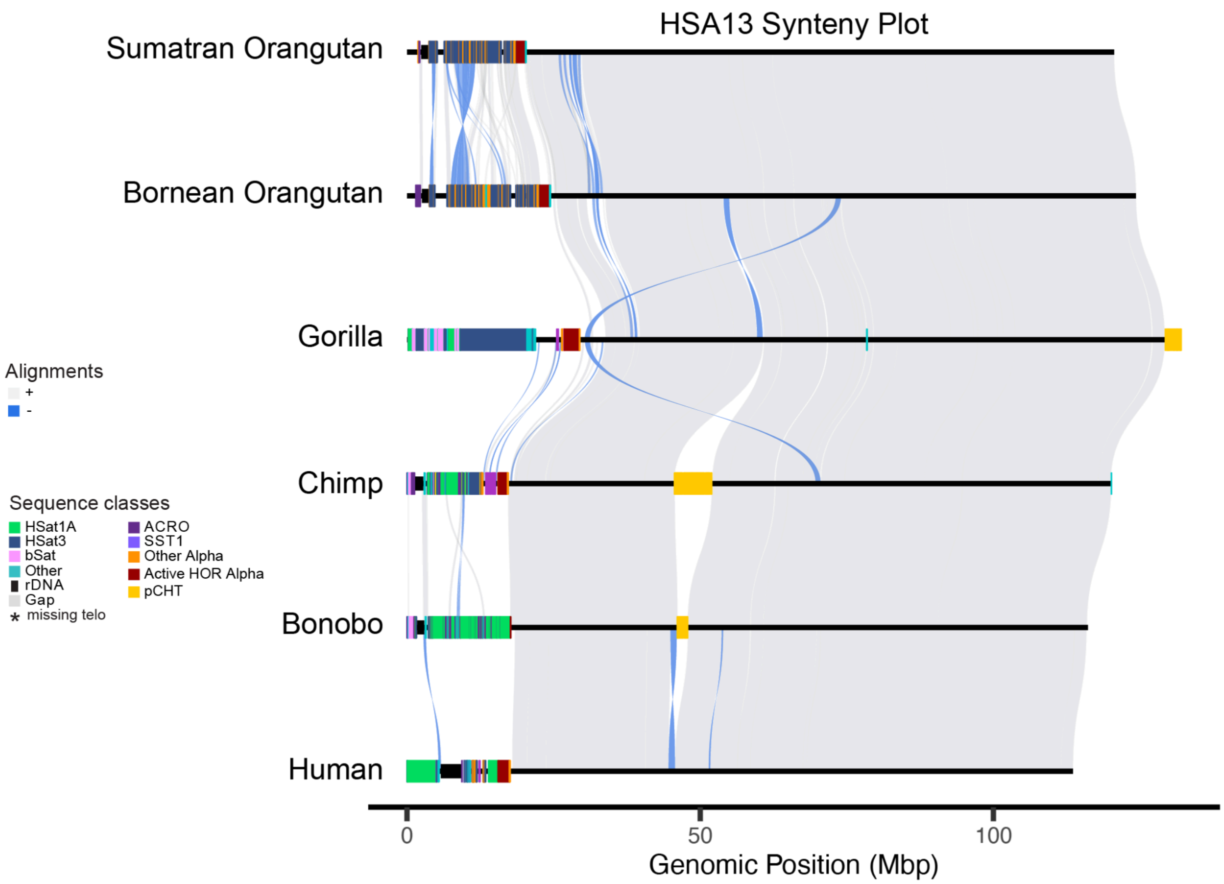


**Figure S13. Whole-chromosome pairwise alignments for HSA13.** SVbyEye visualization of wfmash alignments between pairs of great ape chromosomes. Forward alignments are in grey and reverse complement in blue. Satellite annotations are given along each genome's axis with the centromeres marked in red. There are no apparent whole p-arm inversions.

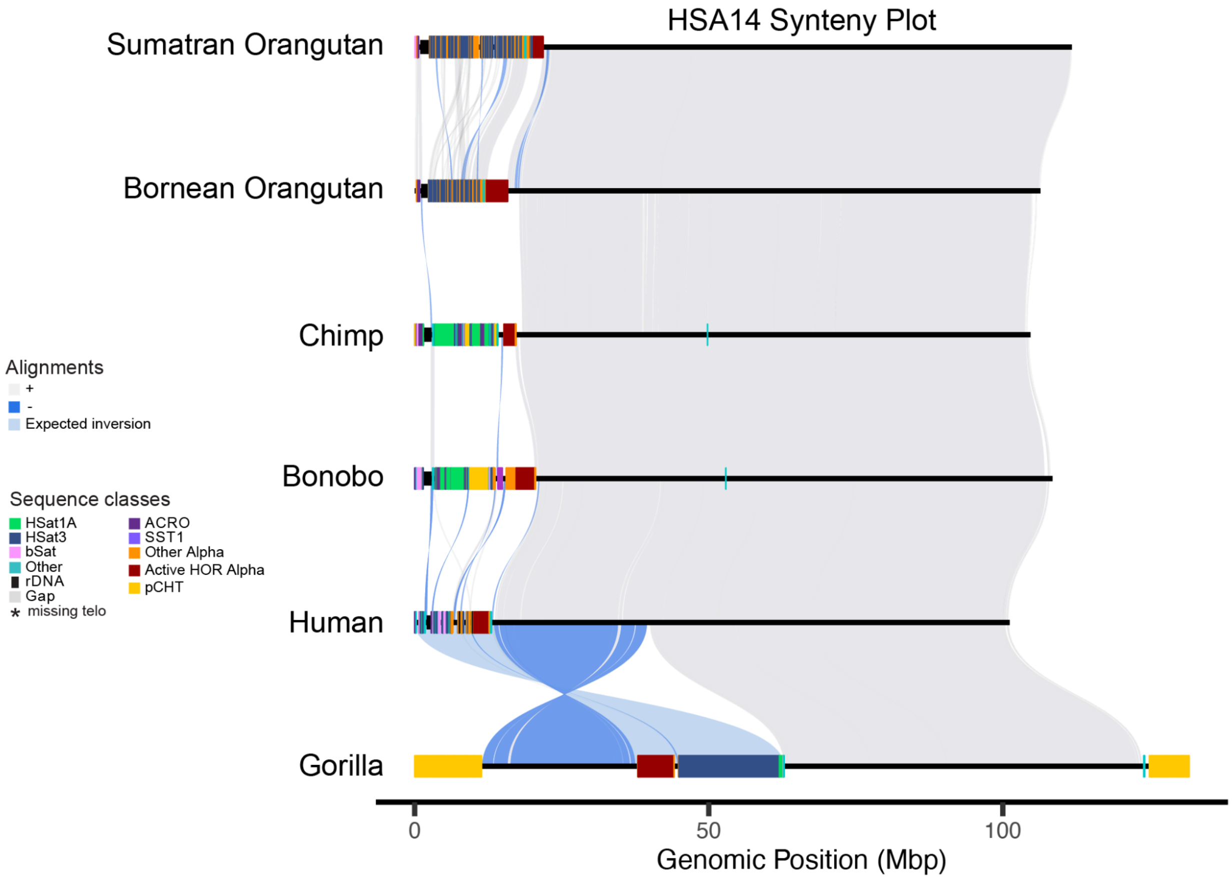


**Figure S14. Whole-chromosome pairwise alignments for HSA14.** SVbyEye visualization of wfmash alignments between pairs of great ape chromosomes. Forward alignments are in grey and reverse complement in blue. Satellite annotations are given along each genome's axis with the centromeres marked in red. The whole p-arm inversion between human and gorilla is shown.

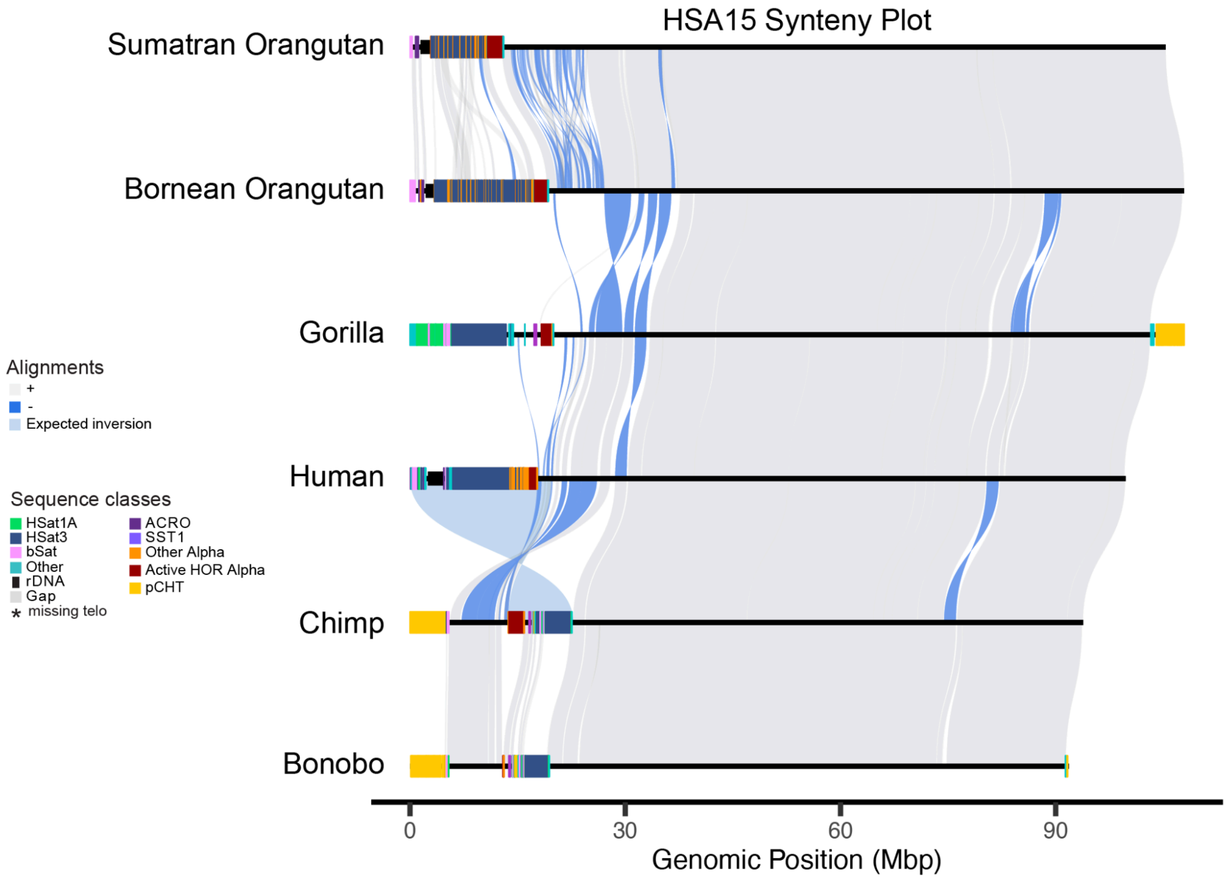


**Figure S15. Whole-chromosome pairwise alignments for HSA15.** SVbyEye visualization of wfmash alignments between pairs of great ape chromosomes. Forward alignments are in grey and reverse complement in blue. Satellite annotations are given along each genome's axis with the centromeres marked in red. The whole p-arm inversion between human and chimp is shown.

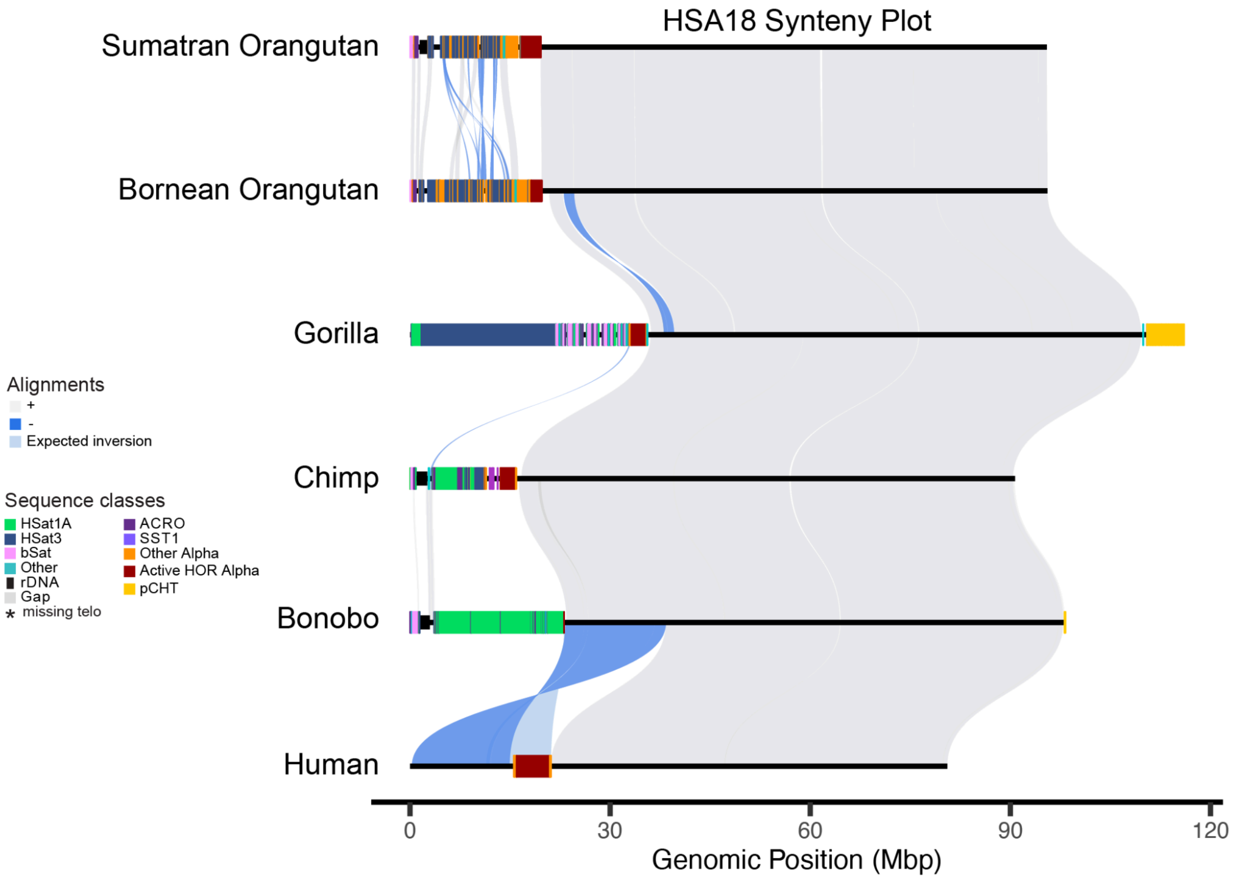


**Figure S16. Whole-chromosome pairwise alignments for HSA18.** SVbyEye visualization of wfmash alignments between pairs of great ape chromosomes. Forward alignments are in grey and reverse complement in blue. Satellite annotations are given along each genome's axis with the centromeres marked in red. The whole p-arm inversion between bonobo and human is shown.

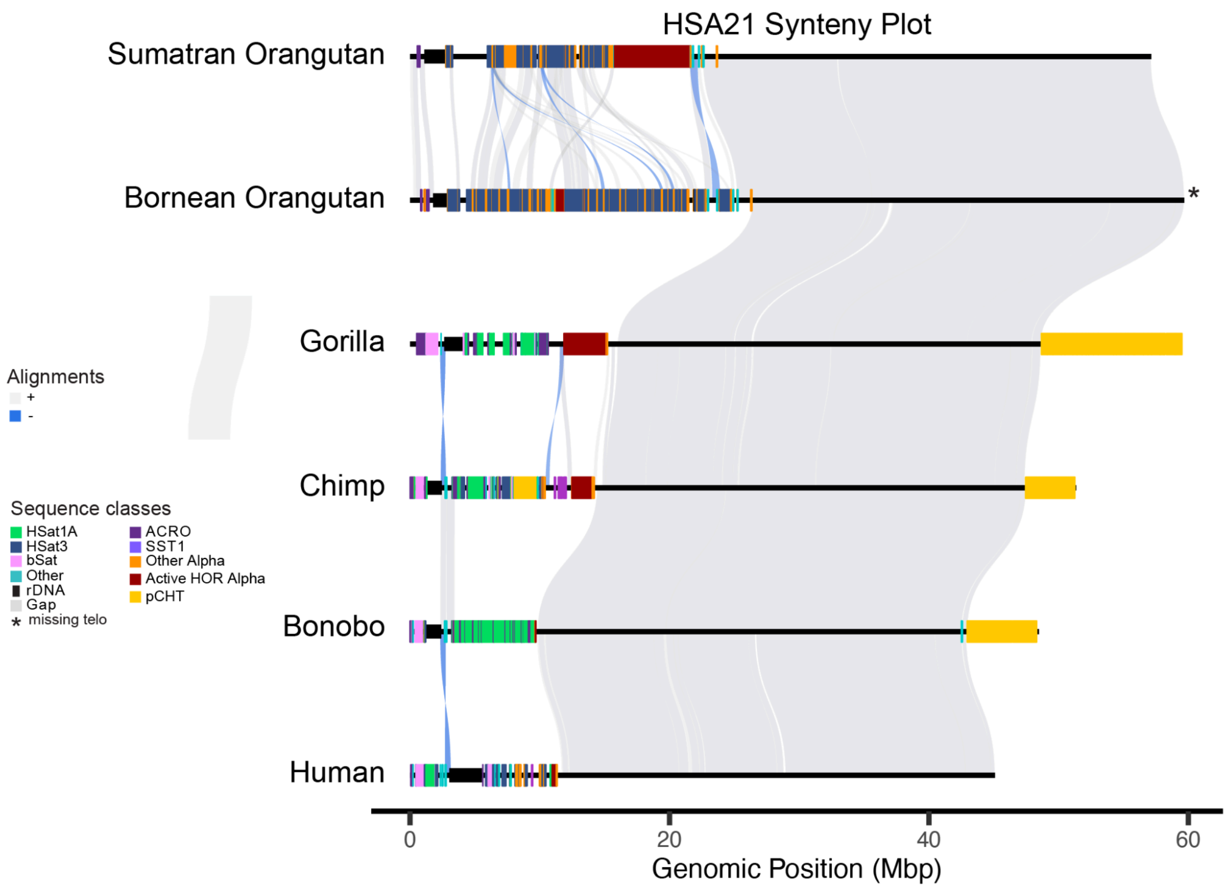


**Figure S17. Whole-chromosome pairwise alignments for HSA21.** SVbyEye visualization of wfmash alignments between pairs of great ape chromosomes. Forward alignments are in grey and reverse complement in blue. Satellite annotations are given along each genome's axis with the centromeres marked in red. There are no apparent whole p-arm inversions.

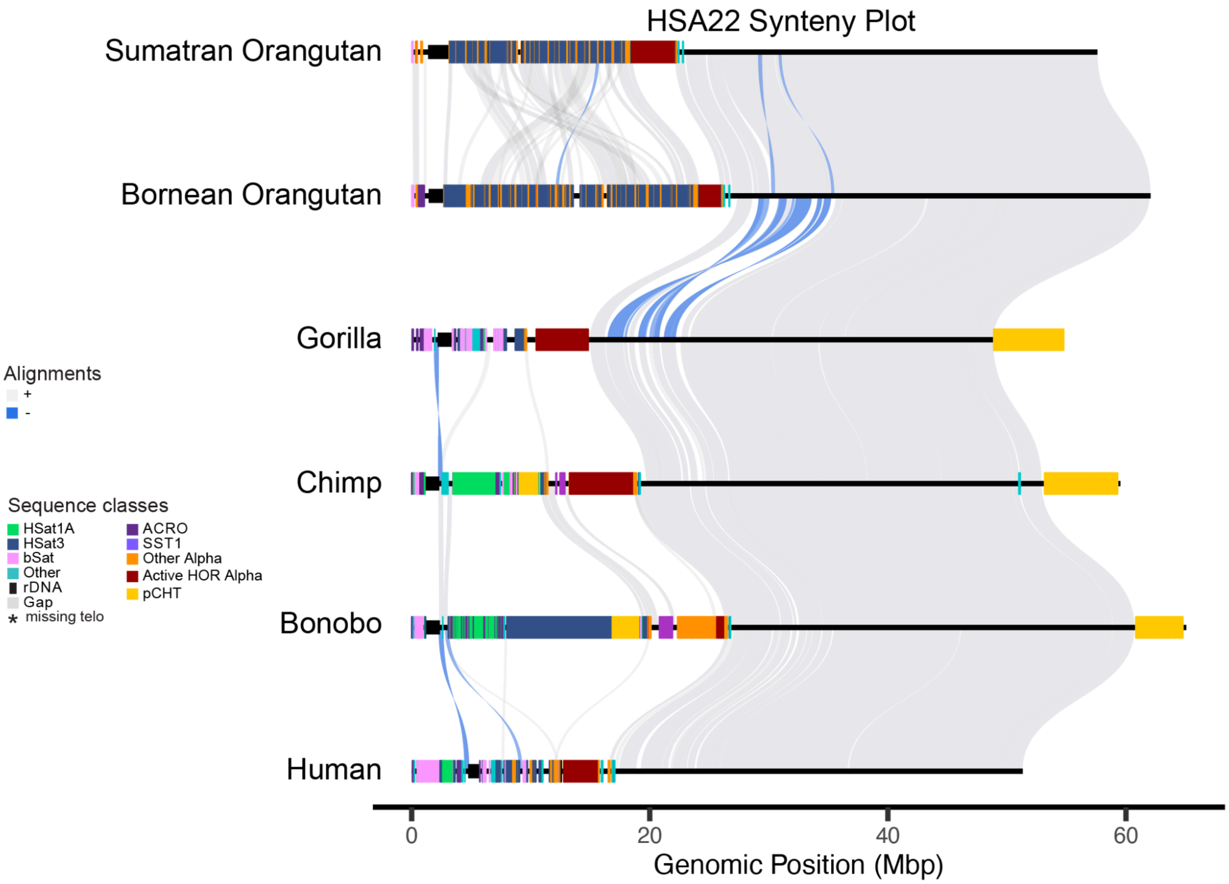


**Figure S18. Whole-chromosome pairwise alignments for HSA22.** SVbyEye visualization of wfmash alignments between pairs of great ape chromosomes. Forward alignments are in grey and reverse complement in blue. Satellite annotations are given along each genome's axis with the centromeres marked in red. There are no apparent whole p-arm inversions.

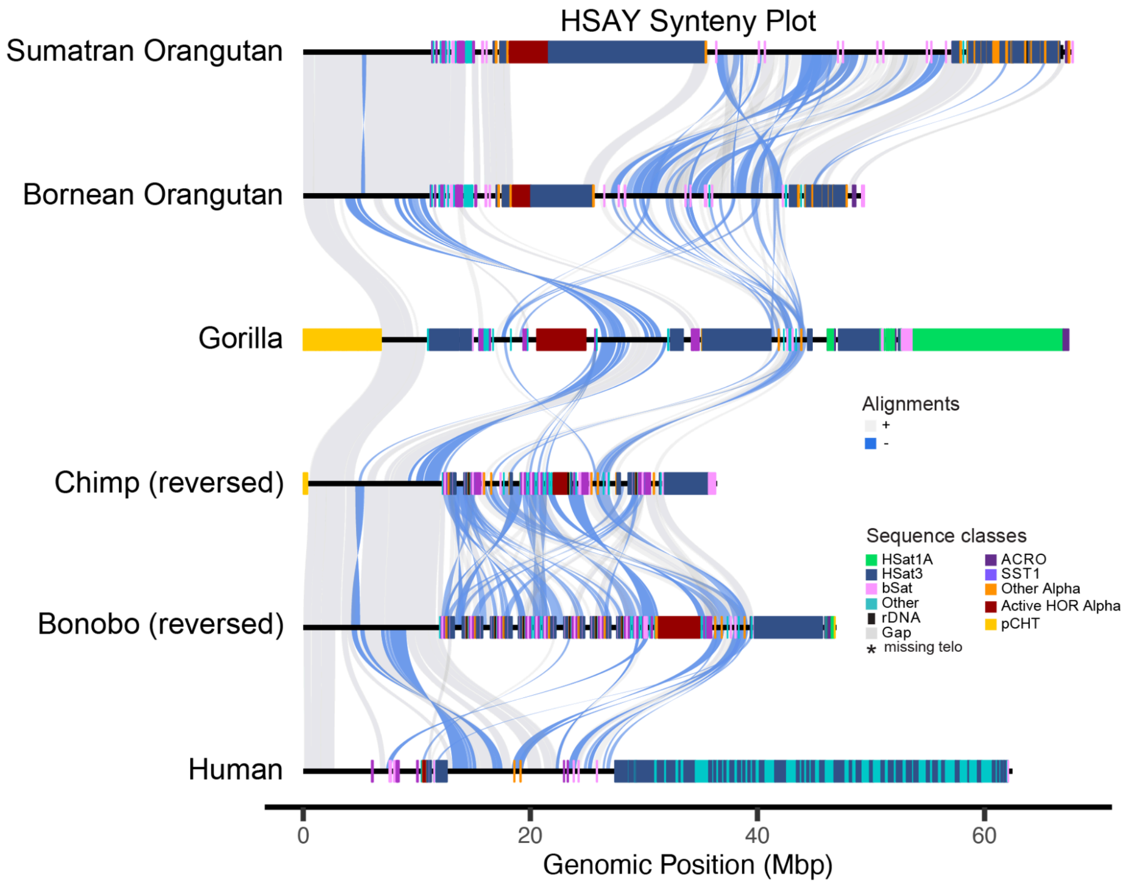


**Figure S19. Whole-chromosome pairwise alignments for HSAY.** SVbyEye visualization of wfmash alignments between pairs of great ape chromosomes. Forward alignments are in grey and reverse complement in blue. Satellite annotations are given along each genome's axis with the centromeres marked in red. All chromosomes are oriented with their pseudoautosomal region (PAR1) on the left, regardless of centromere position.

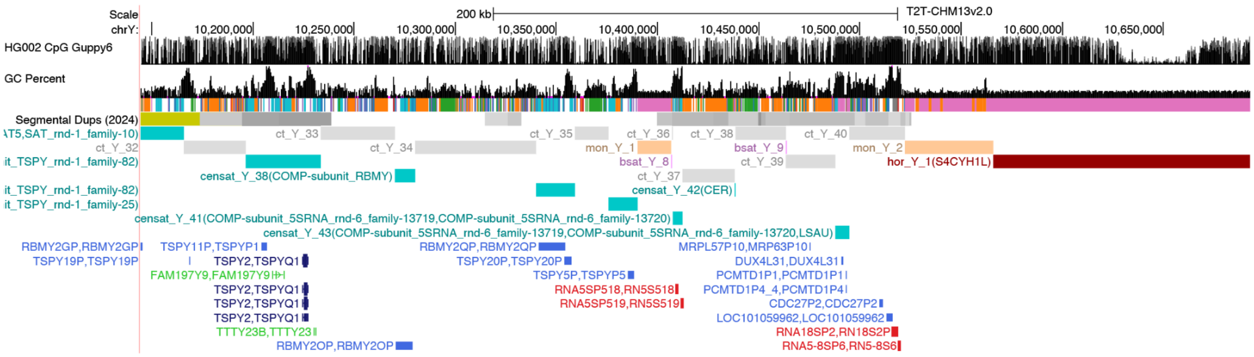


**Figure S20. Location of 45S rDNA fragments in the pericentromere of human HG002 ChrY.** UCSC Genome Browser tracks from top to bottom show: %methylation, %GC, RepeatMasker annotations, segmental duplications, satellite annotations, and gene annotations. The centromeric, active αSat array is marked in dark red (hor_Y_1). Annotated rDNAs are below in red, including both 5S (left) and 45S/18S/5.8S (right).

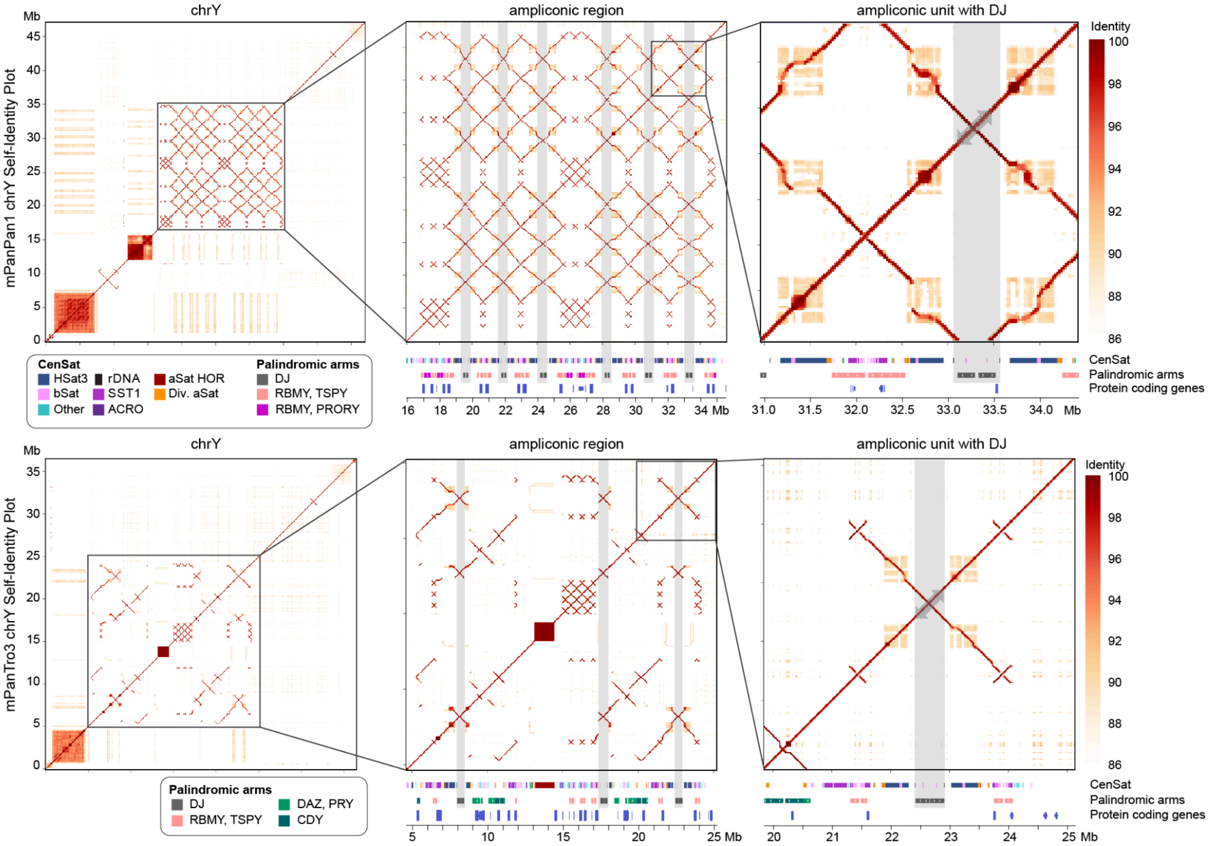


**Figure S21. Amplification of a 45S rDNA DJ-like sequence alongside *TSPY* and *RBMY* on ChrY of chimp and bonobo.** Chimp and bonobo Y chromosomes contain amplified, palindromic arrays of *TSPY* and *RBMY* genes. Spacing between these amplified gene copies is the palindromic repeat unit characteristic of the 45S rDNA distal junction sequence (highlighted in gray). These amplified regions are progressively zoomed from left to right to show the fine-grained structure.

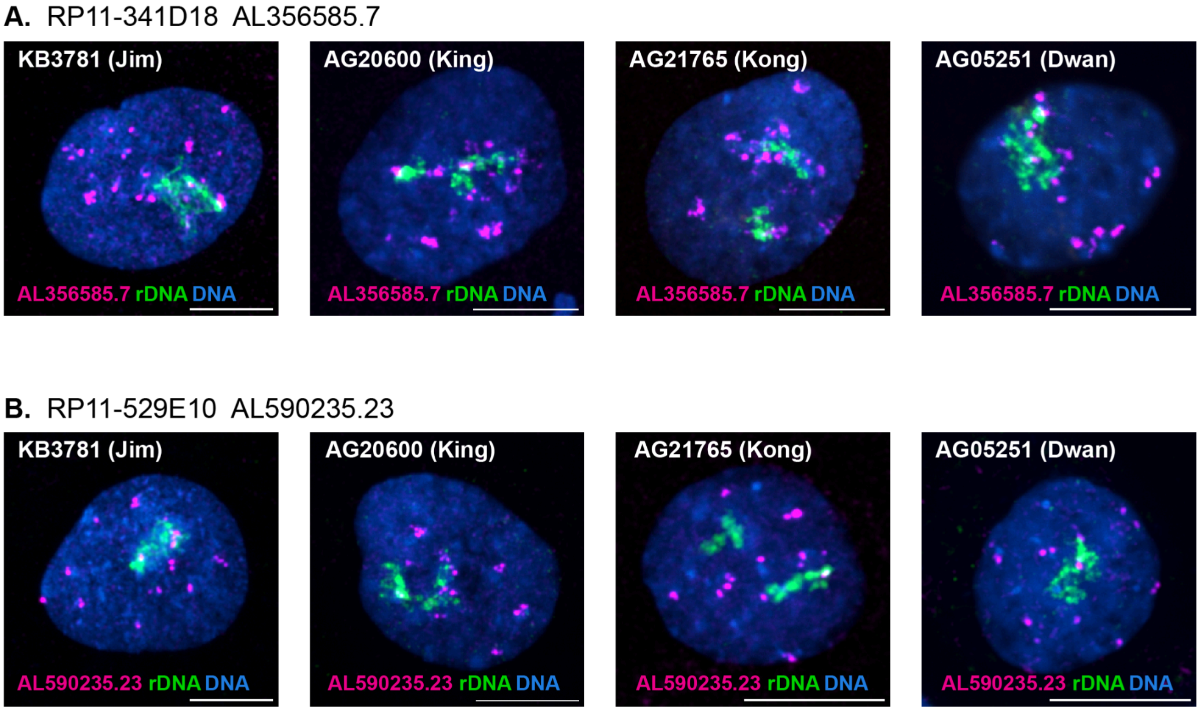


**Figure S22. Localization of amplified gene regions and rDNA in representative interphase fibroblast nuclei of four different gorillas.** (A) Most *GGT*-associated signals from BAC RP11-341D18 (AL356585.7, magenta) were not associated with rDNA (BAC RP11-450E20, green). (B) Nor were the *LRPAP1*-associated signals from BAC RP11-529E10 (AL590235.23). This suggests the gorilla rDNA– chromosomes do not organize with the rDNA+ chromosomes and may not participate in the formation of nucleoli. Nuclei were counter-stained with DAPI. Bar, 10μm.

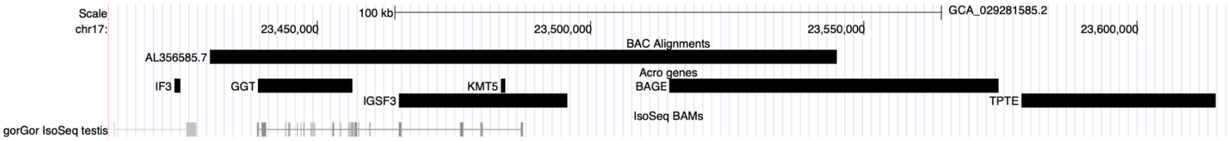


**Figure S23. Alignment of BAC RP11-341D18 to the gorilla genome.** Human BAC RP11-341D18 (AL356585.7) is shown aligned to GGO-HSA-18 (gorilla chr17) overlapping the amplified gorilla genes: *IGSF3-GGT* (fusion) and *BAGE2*. Overlaid transcript evidence from gorilla testes is shown at bottom.

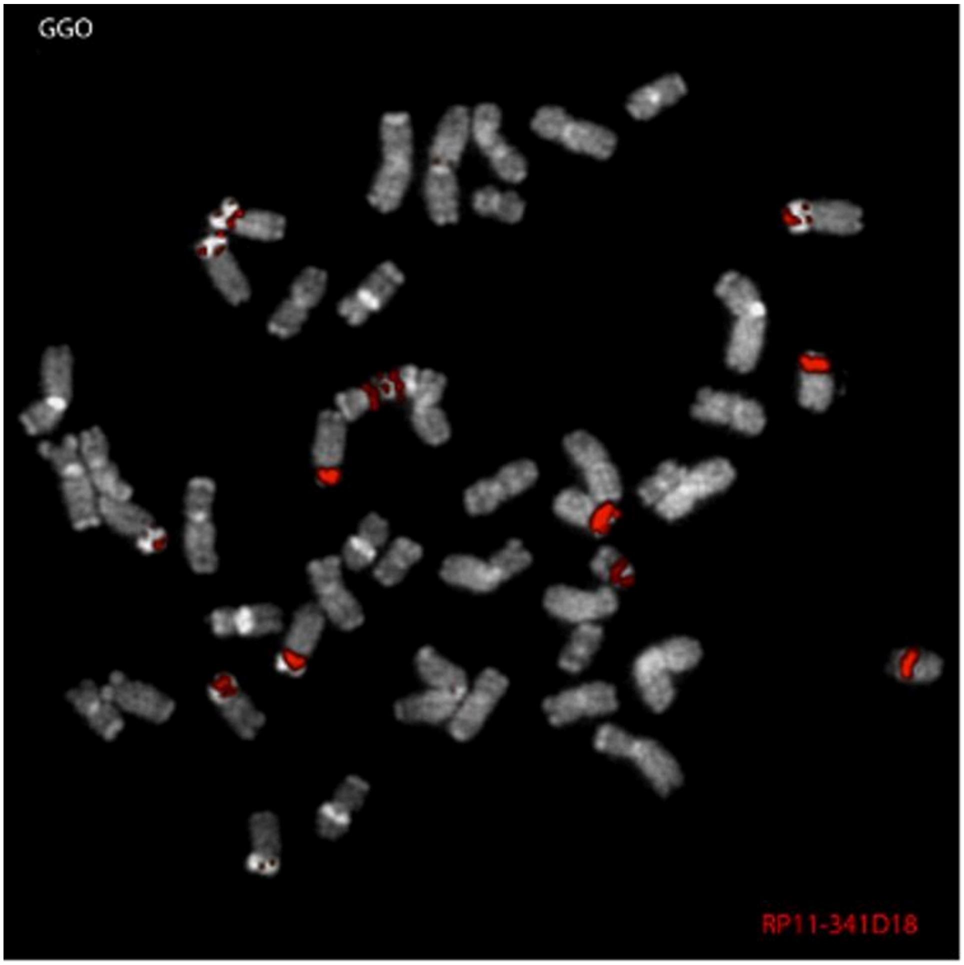


**Figure S24. Metaphase FISH of the *GGT* amplification in gorilla.** FISH analysis using BAC probe RP11-341D18 (AL356585.7) mapping to the amplified *IGSF3-GGT* locus on metaphase chromosomes from *Gorilla gorilla gorilla* (individual “GGO”), confirming amplification across the rDNA– acrocentrics. Consistent hybridization patterns were observed across six different gorilla individuals as summarized in Supplementary Table S3.

RP11-341D18 AL356585.7

**KB3781 (Jim)**

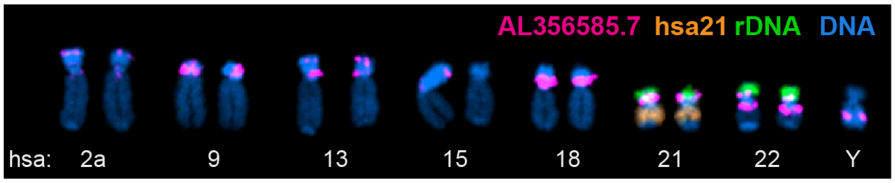


**AG20600 (King)**

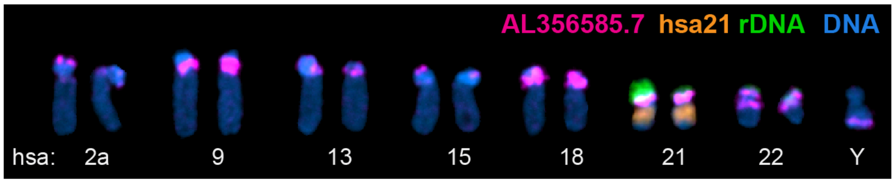


**AG21765 (Kong)**

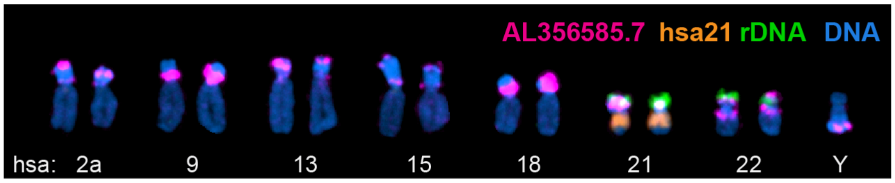


**AG05251 (Dwan)**

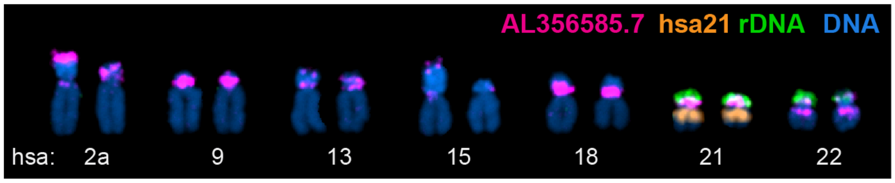


**Figure S25. Chromosome-labeled FISH of the *GGT* amplification in gorilla.** Representative acrocentric karyograms from primary fibroblast cell lines of four different gorilla individuals (3 male, 1 female) labeled by FISH using BAC probe RP11-341D18 (AL356585.7, magenta). rDNA was labeled with BAC RP11-450E20 (green) and HSA21 was labeled with human Chromosome 21 paint (orange). Chromosomes were counterstained with DAPI. The *IGSF3-GGT* locus is consistently amplified across HSA9/18/21, with scattered distal copies on the other short arms, consistent with the T2T genome annotation. The ancestral location of *GGT1/2* on the long arm of HSA22 is also shown. Note that “Jim” is the same animal used for the T2T reference genome assembly.

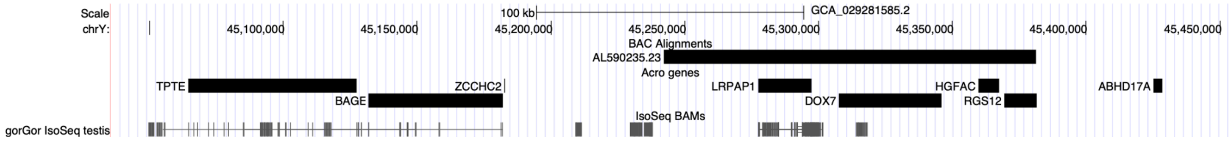


**Figure S26. Alignment of BAC RP11-529E10 to the gorilla genome.** Human BAC RP11-529E10 (AL590235.23) is shown aligned to gorilla ChrY overlapping the amplified gorilla gene *LRPAP1*. Overlaid transcript evidence from gorilla testes is shown at bottom.

RP11-529E10  AL590235.23

**KB3781 (Jim)**

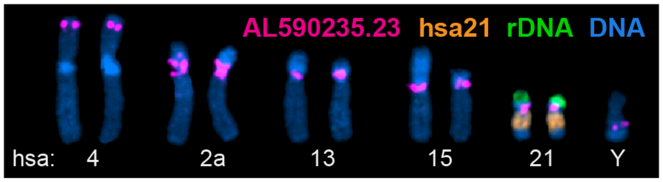


**AG20600 (King)**

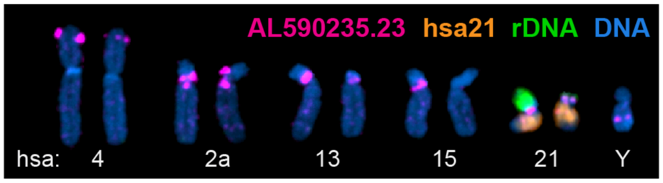


**AG21765 (Kong)**

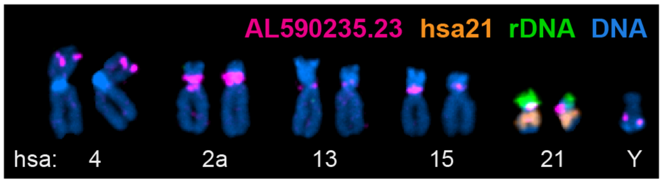


**AG05251 (Dwan)**

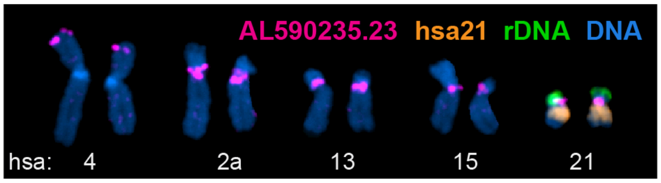


**Figure S27. Chromosome-labeled FISH of the *LRPAP1* (*FRG1*) amplification in gorilla.** Representative acrocentric karyograms from primary fibroblast cell lines of four different gorilla individuals (3 male, 1 female) labeled by FISH using BAC probe RP11-529E10 (AL590235.23, magenta) mapping to the *LRPAP1* locus. rDNA was labeled with BAC RP11-450E20 (green), HSA21 was labeled with human Chromosome 21 paint (orange). Chromosomes were counterstained with DAPI. Based on the gorilla genome annotation, *LRPAP1* genes are typically found in the same amplified cluster as *FRG1*. Amplification of this locus across the pericentromeres of HSA2A/13/15 and HSA21 is visible, consistent with the T2T genome annotation. The ancestral location of *LRPAP1* on the short arm of HSA4 is also shown. Note that “Jim” is the same animal used for the T2T reference genome assembly.

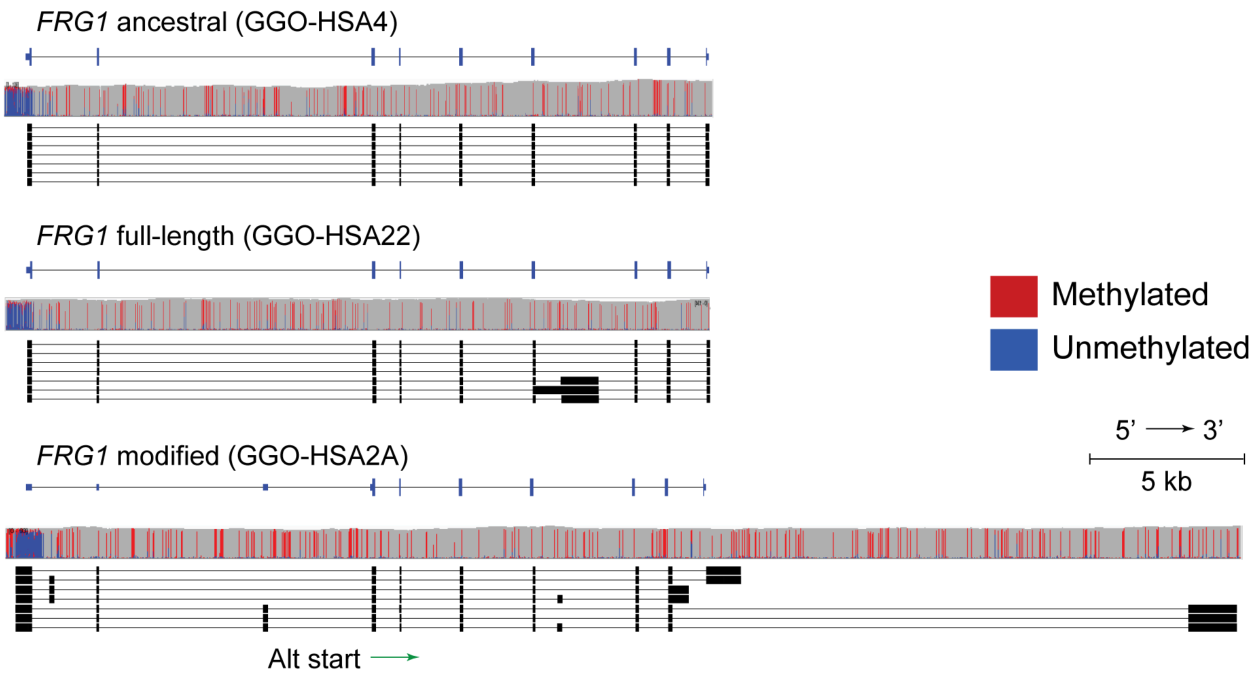


**S28. Methylation pattern and transcript structure across the ancestral, duplicated full copies, and modified FRG1 copies.** ANC, the ancestral HSA4 copy present on GGO-HSA4; DUP, an example duplicated copy from GGO-HSA22; MOD, an example modified copy from GGO-HSA2A. The canonical gorilla FRG1 transcript is shown at top. For each copy group, the upper track shows per-read methylation (red, methylated; blue, unmethylated) and the lower tracks show long-read transcript isoforms from the fibroblast Iso-Seq data; the green arrow marks the alternative start codon in exon 3 used by the modified copies.

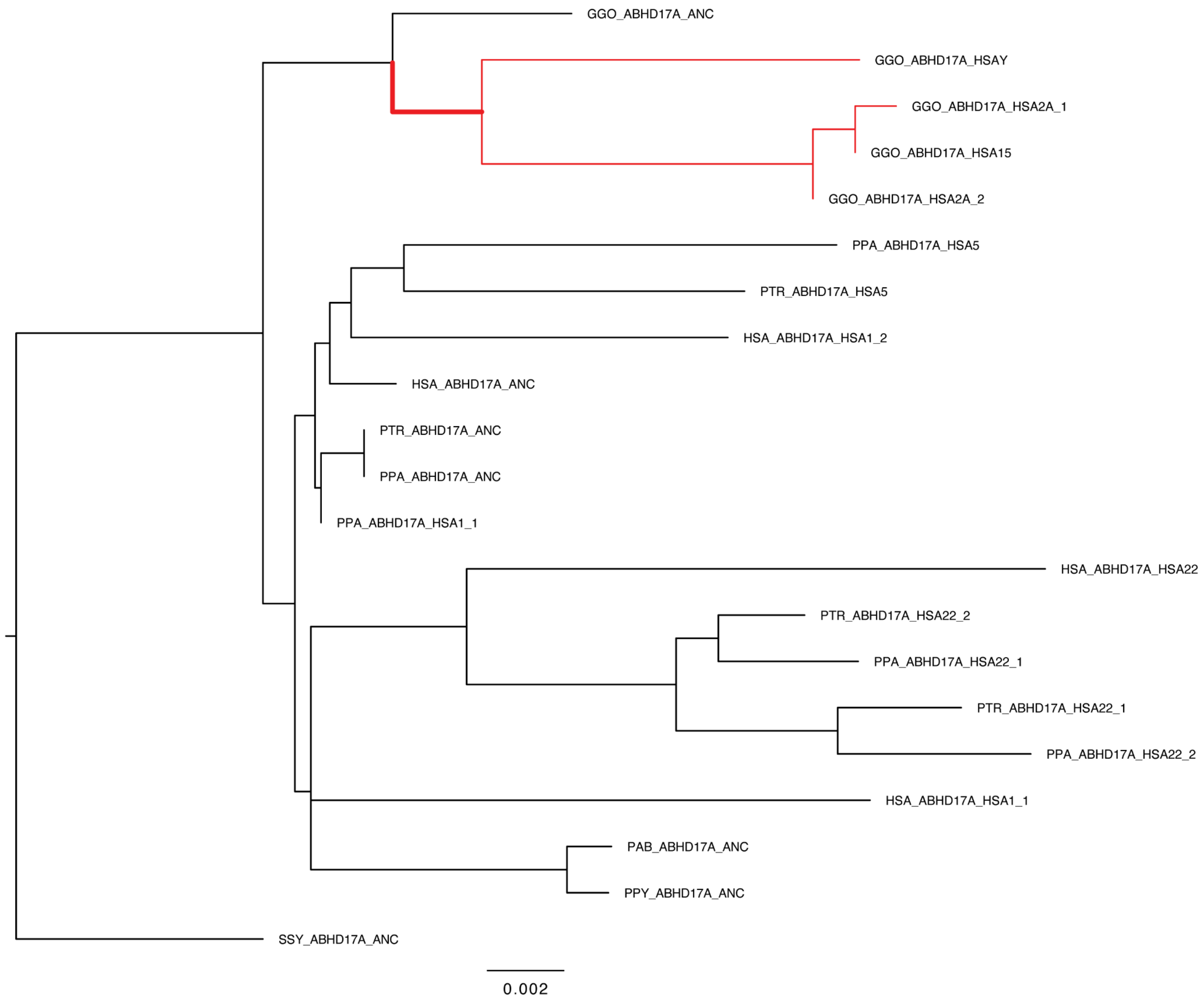


**Figure S29. CDS tree for *ABHD17A*.** ML tree built from the CDS alignment. Studied expansions are indicated with the red-highlighted branches with all selection tests run by using these entire sets or subsets as foreground.

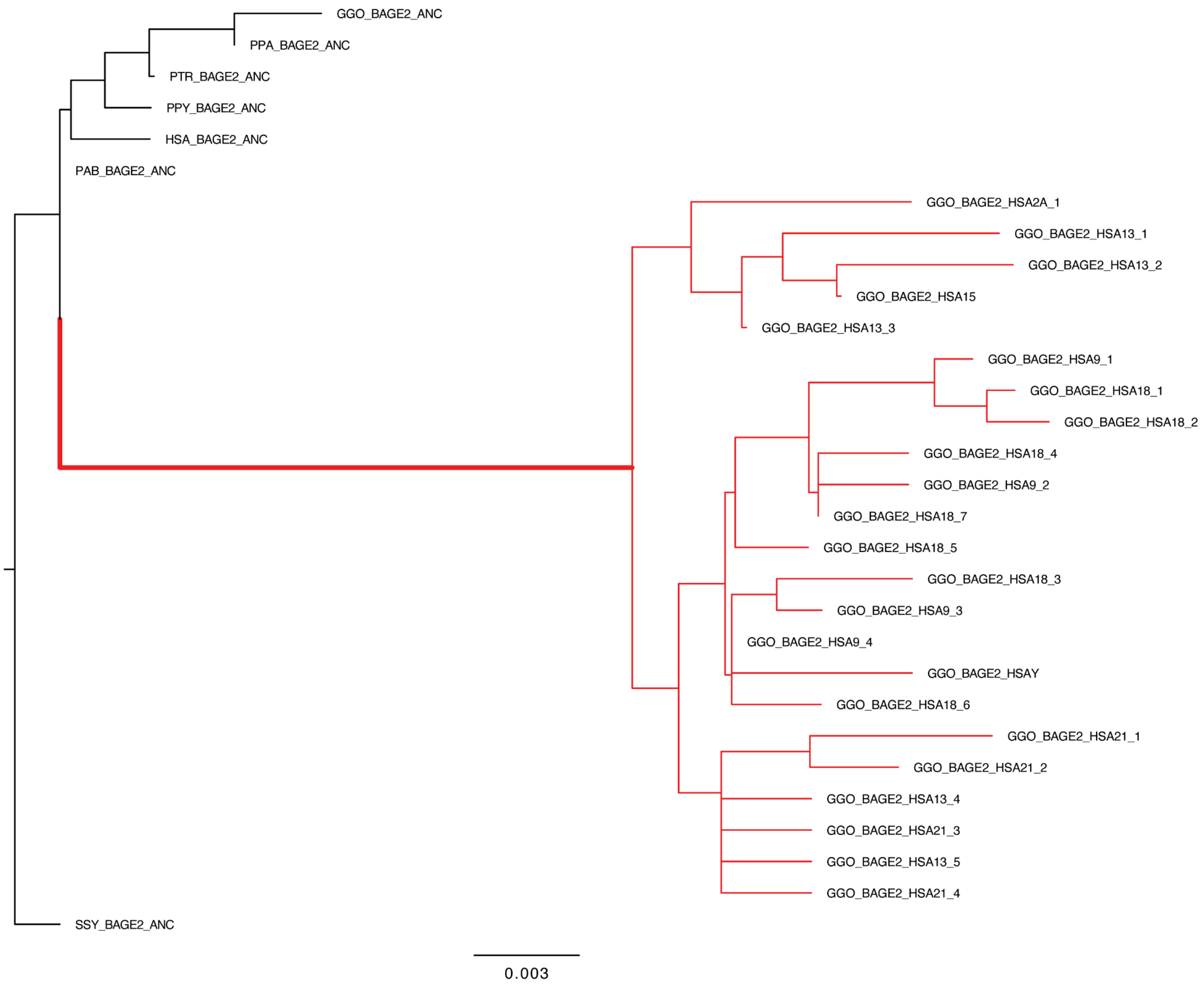


**Figure S30. CDS tree for *BAGE2*.** ML tree built from the CDS alignment. Studied expansions are indicated with the red-highlighted branches with all selection tests run by using these entire sets or subsets as foreground.

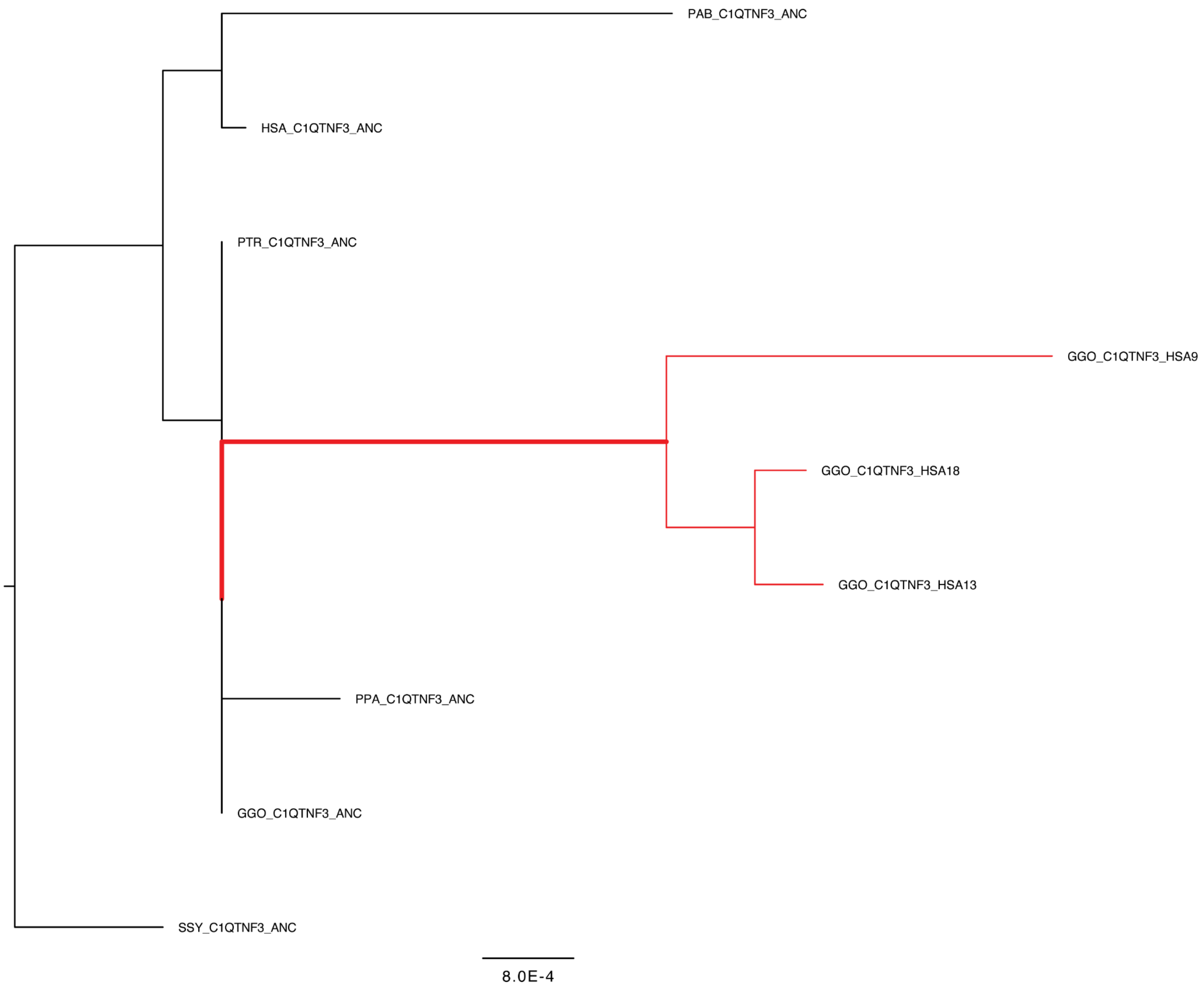


**Figure S31. CDS tree for *C1QTNF3*.** ML tree built from the CDS alignment. Studied expansions are indicated with the red-highlighted branches with all selection tests run by using these entire sets or subsets as foreground.

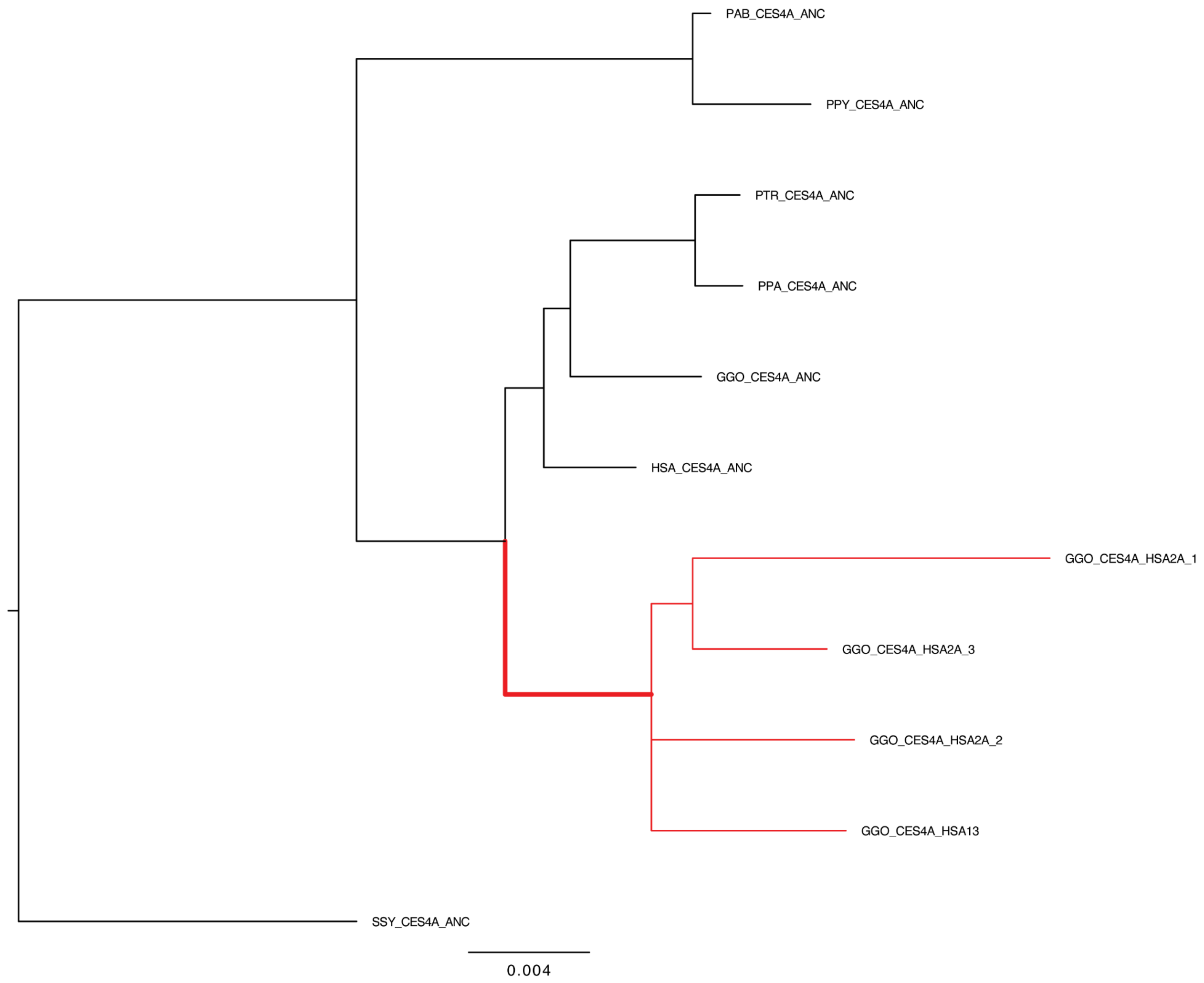


**Figure S32. CDS tree for *CES4A*.** ML tree built from the CDS alignment. Studied expansions are indicated with the red-highlighted branches with all selection tests run by using these entire sets or subsets as foreground.

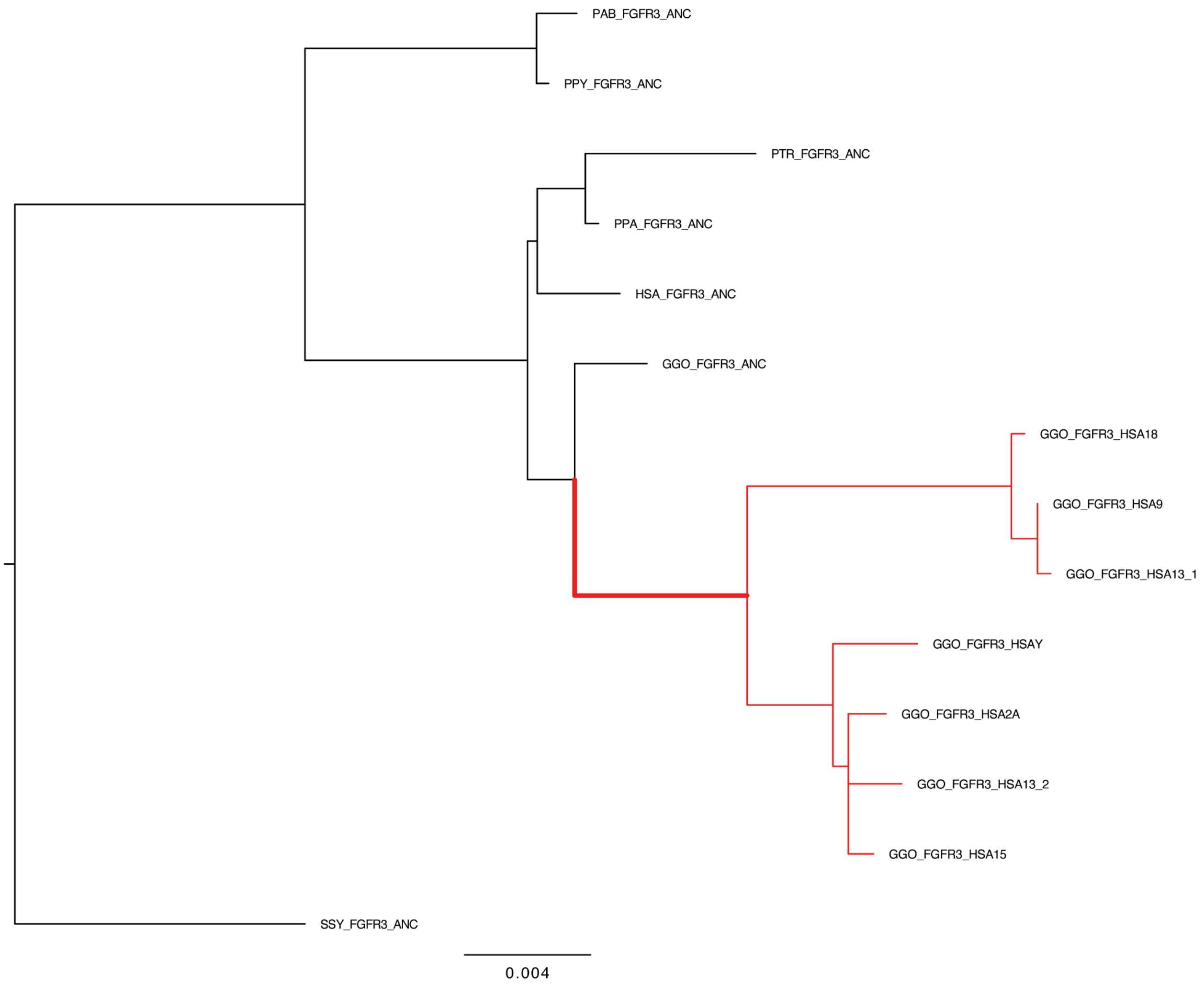


**Figure S33. CDS tree for *FGFR3*.** ML tree built from the CDS alignment. Studied expansions are indicated with the red-highlighted branches with all selection tests run by using these entire sets or subsets as foreground.

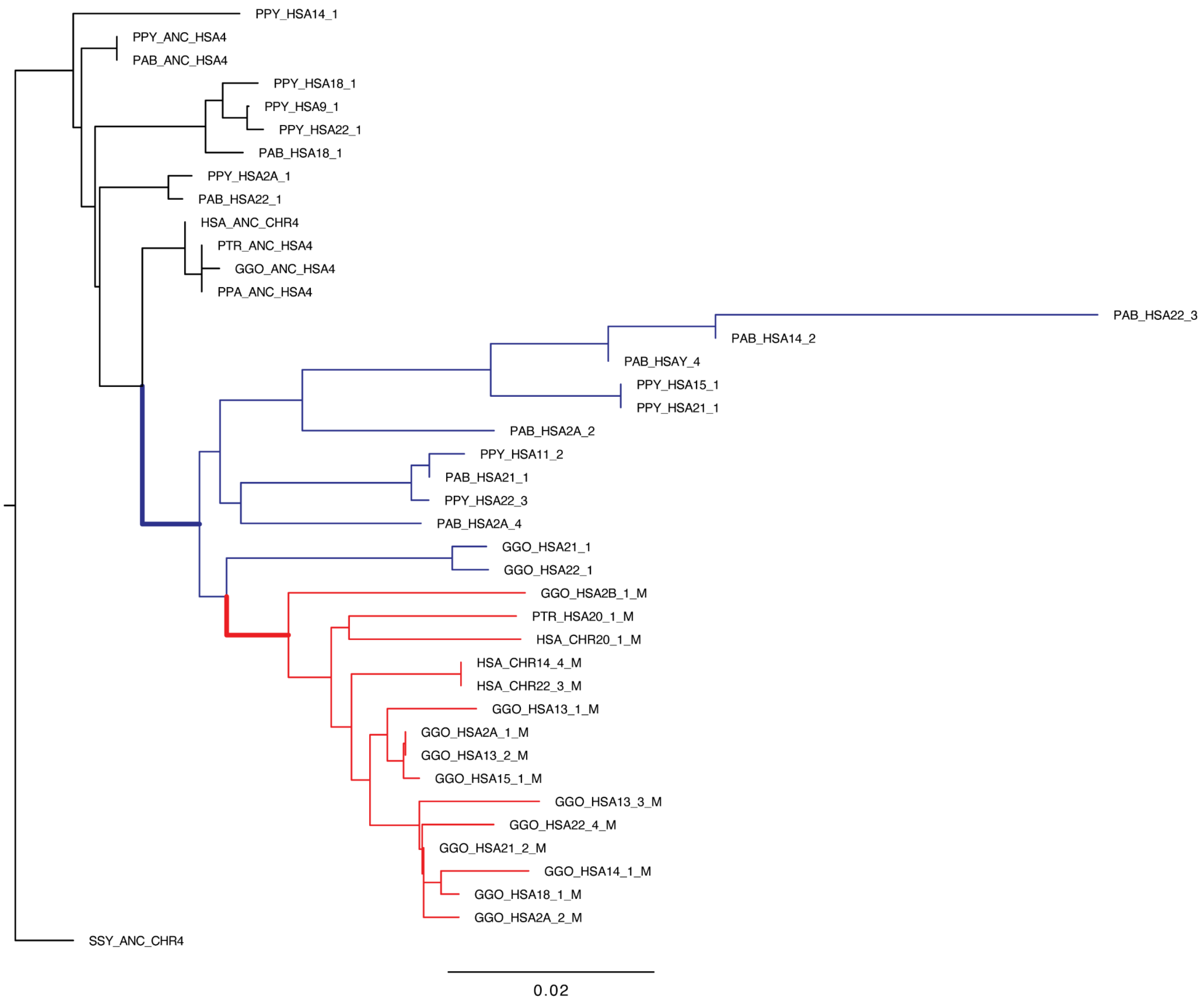


**Figure S34. CDS tree for *FRG1*.** ML tree built from the CDS alignment. BUSTED was run separately with two foreground sets: the great ape expansion (in blue & red), and the modified copies of gorilla, human, and chimp (in red).

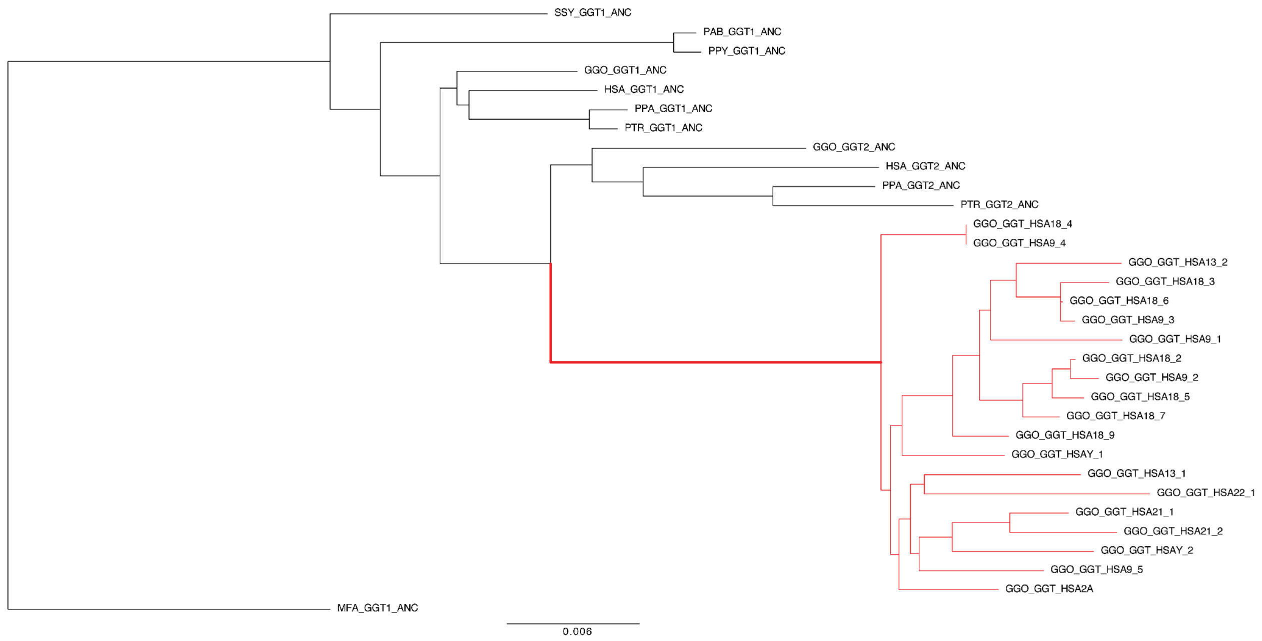


**Figure S35. CDS tree for *GGT*.** ML tree built from the CDS alignment. Studied expansions are indicated with the red-highlighted branches with all selection tests run by using these entire sets or subsets as foreground.

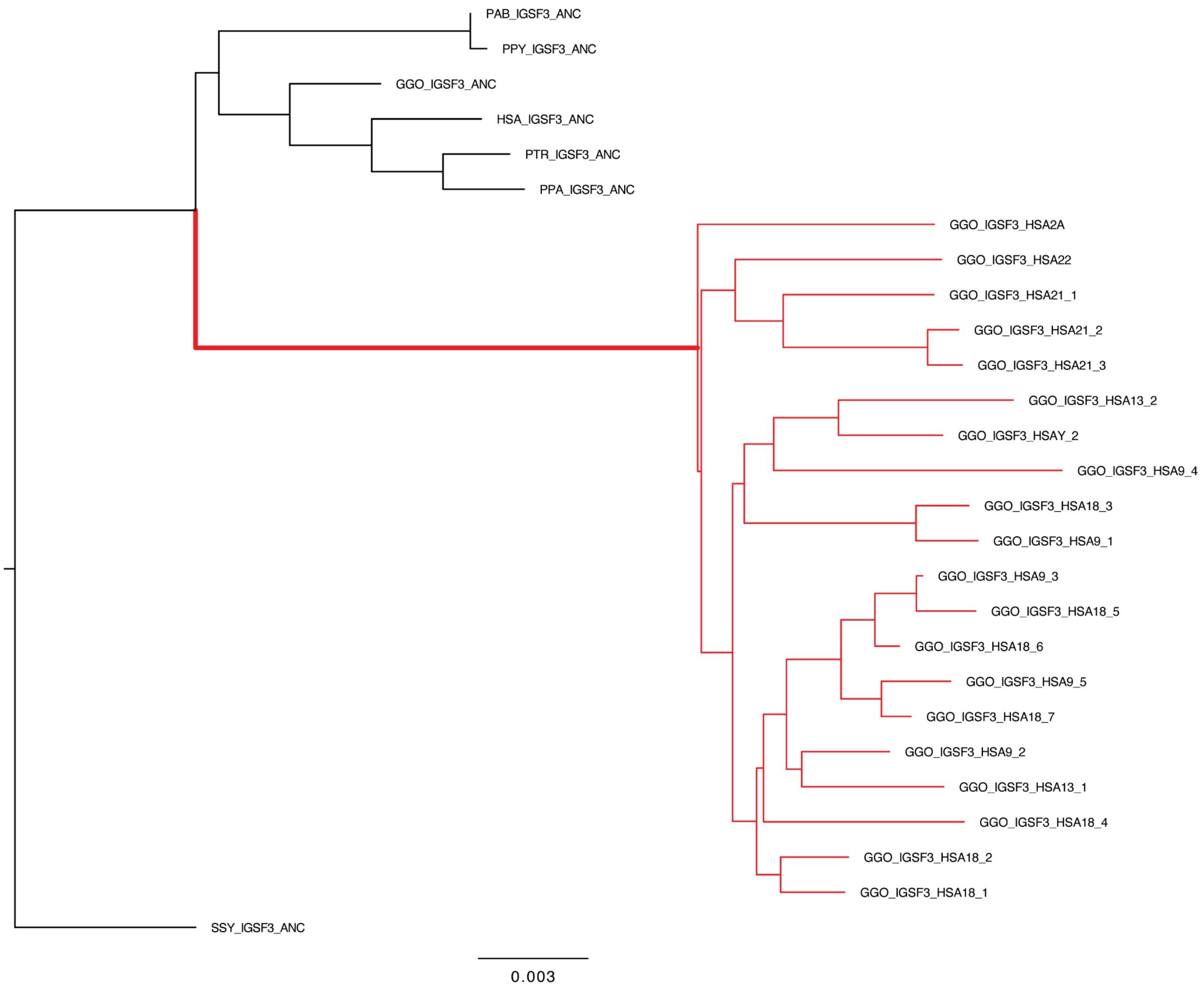


**Figure S36. CDS tree for *IGSF3*.** ML tree built from the CDS alignment. Studied expansions are indicated with the red-highlighted branches with all selection tests run by using these entire sets or subsets as foreground.

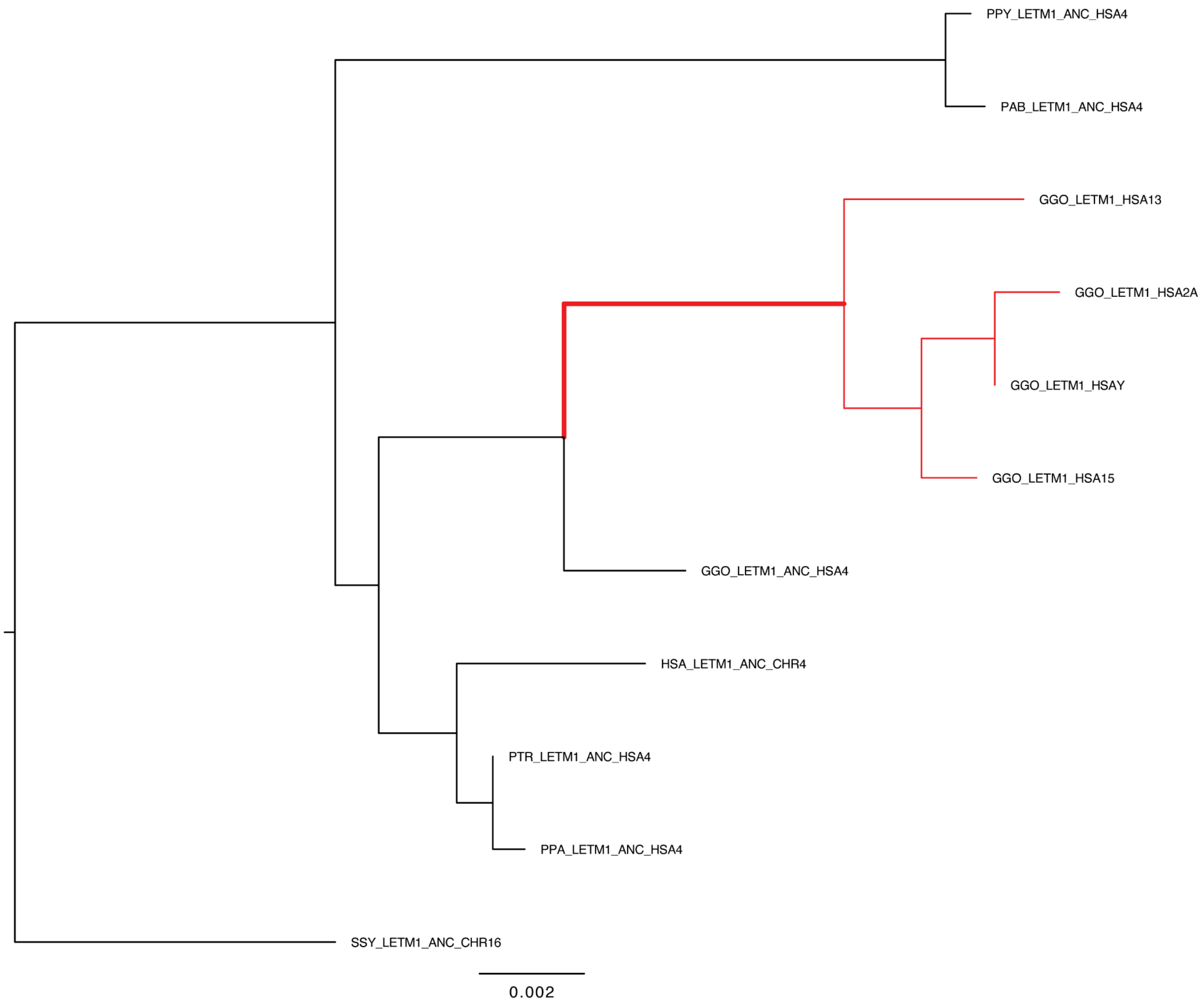


**Figure S37. CDS tree for *LETM1*.** ML tree built from the CDS alignment. Studied expansions are indicated with the red-highlighted branches with all selection tests run by using these entire sets or subsets as foreground.

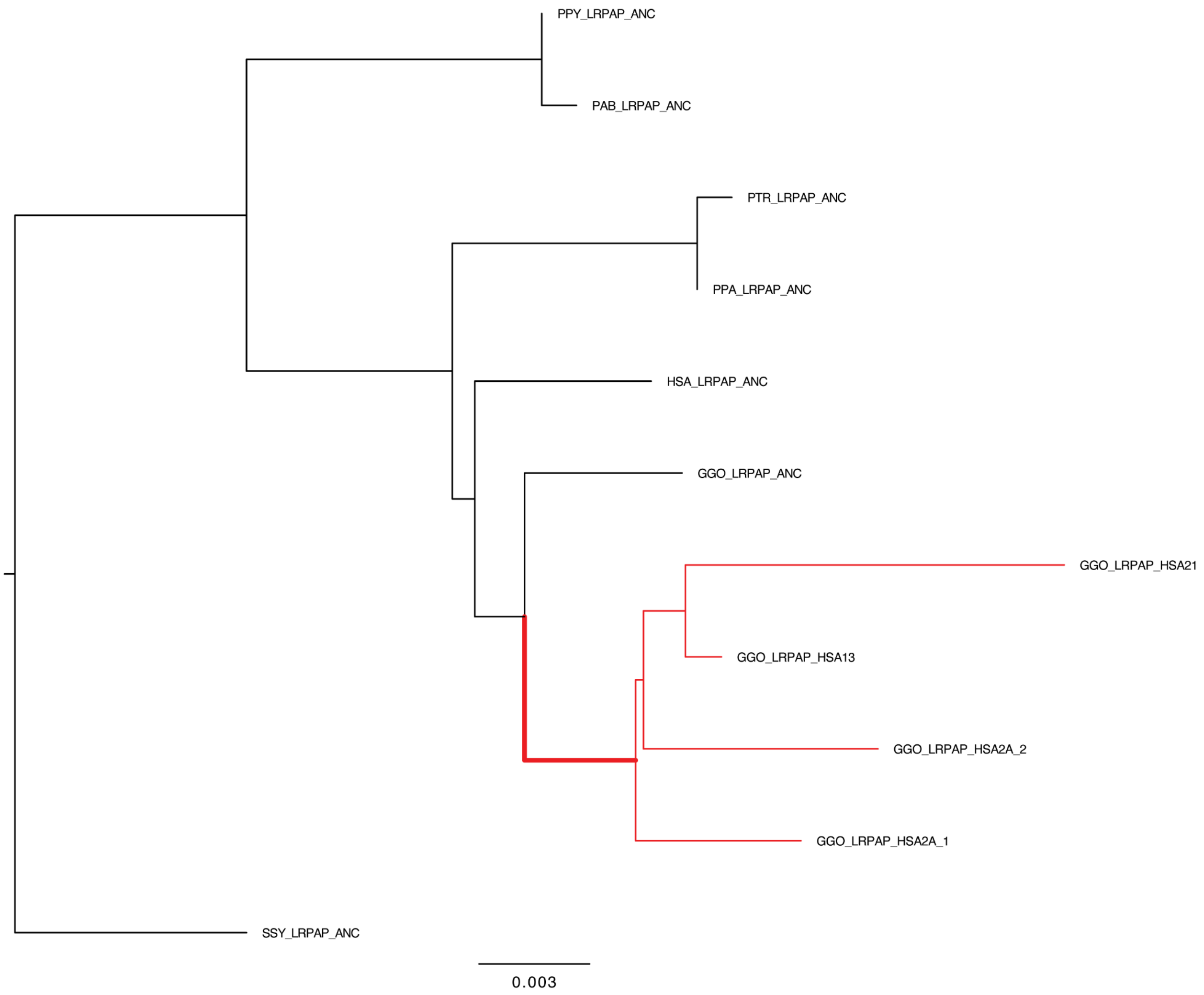


**Figure S38. CDS tree for *LRPAP1*.** ML tree built from the CDS alignment. Studied expansions are indicated with the red-highlighted branches with all selection tests run by using these entire sets or subsets as foreground.

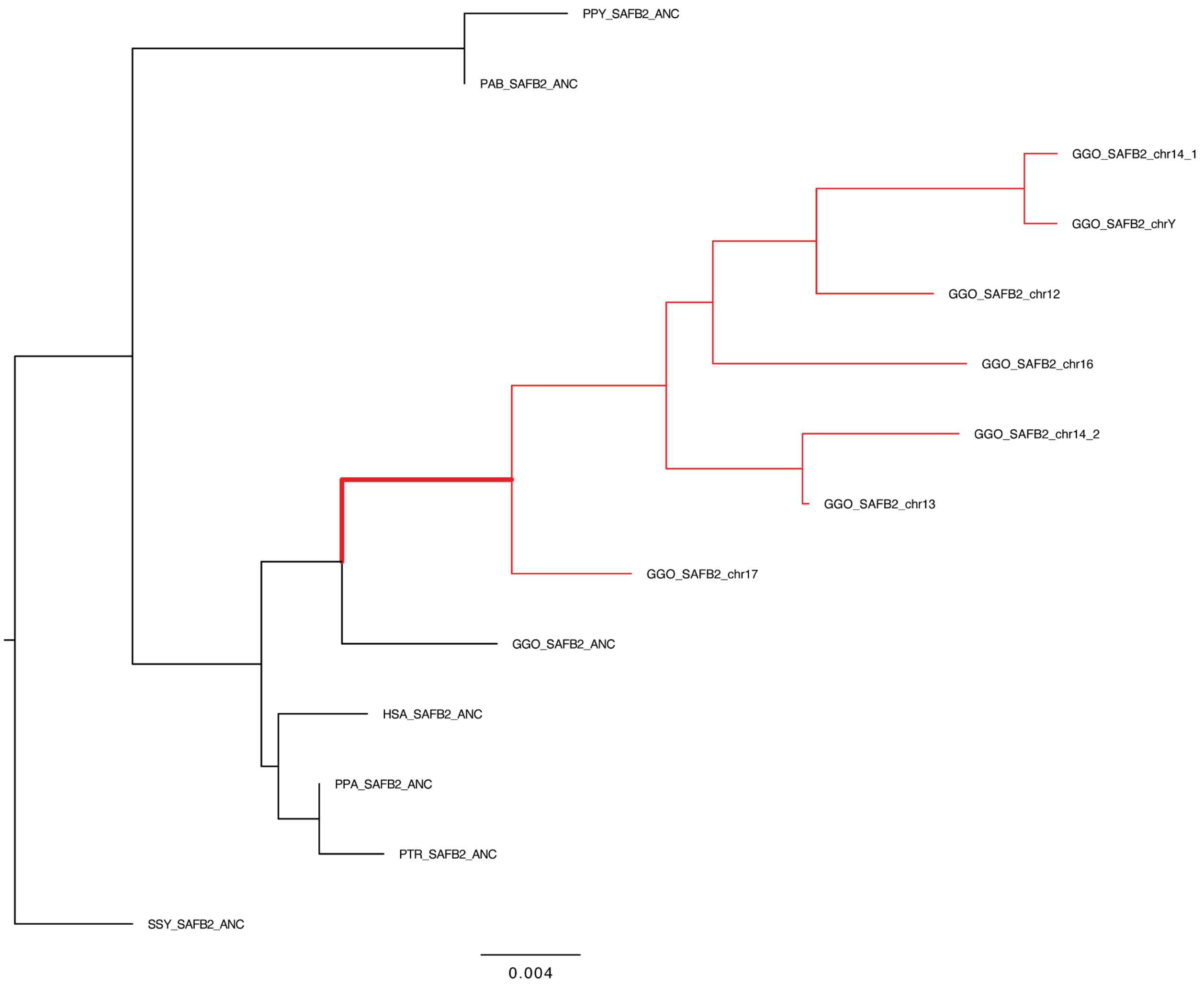


**Figure S39. CDS tree for *SAFB2*.** ML tree built from the CDS alignment. Studied expansions are indicated with the red-highlighted branches with all selection tests run by using these entire sets or subsets as foreground.

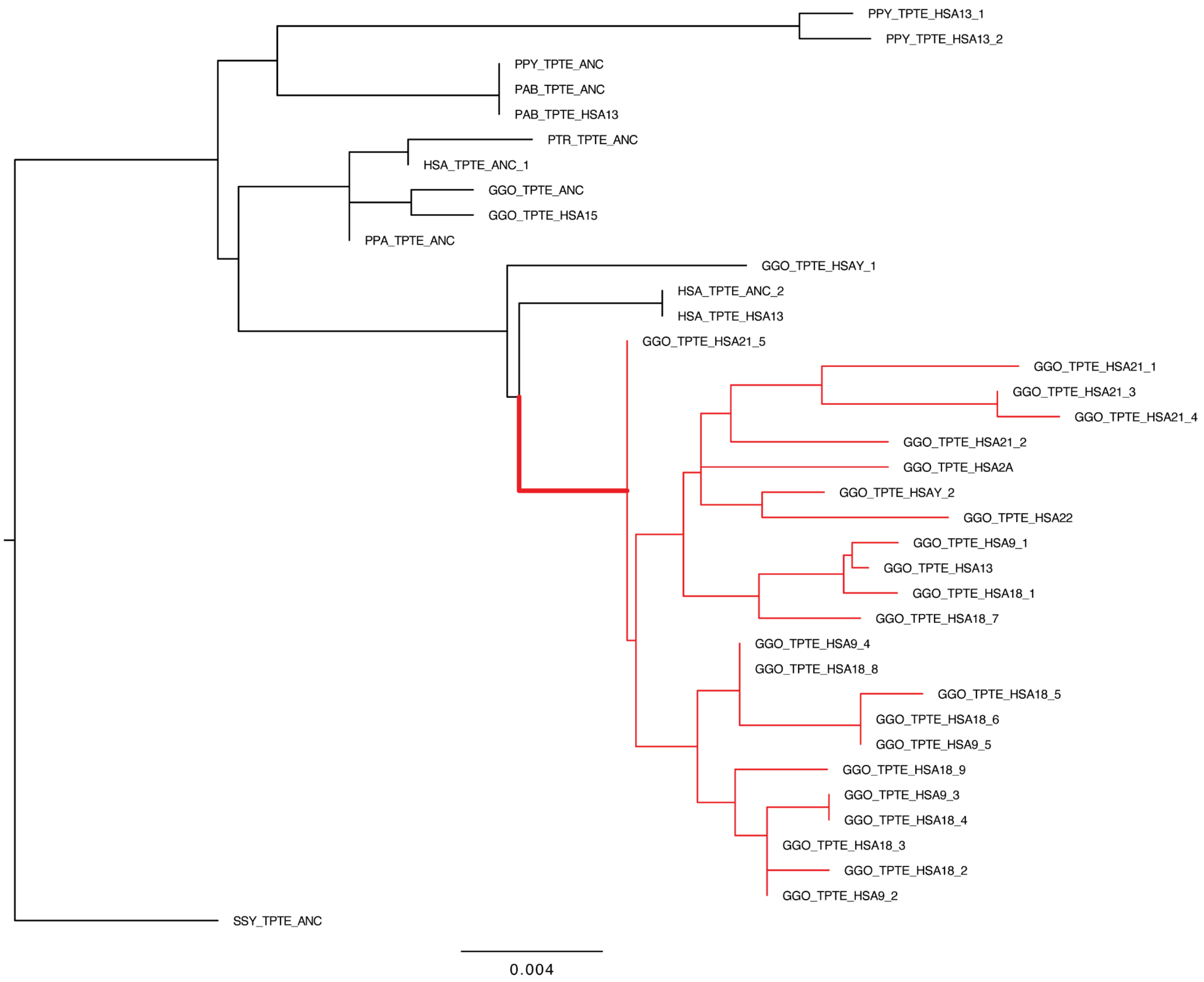


**Figure S40. CDS tree for *TPTE*.** ML tree built from the CDS alignment. Studied expansions are indicated with the red-highlighted branches with all selection tests run by using these entire sets or subsets as foreground.

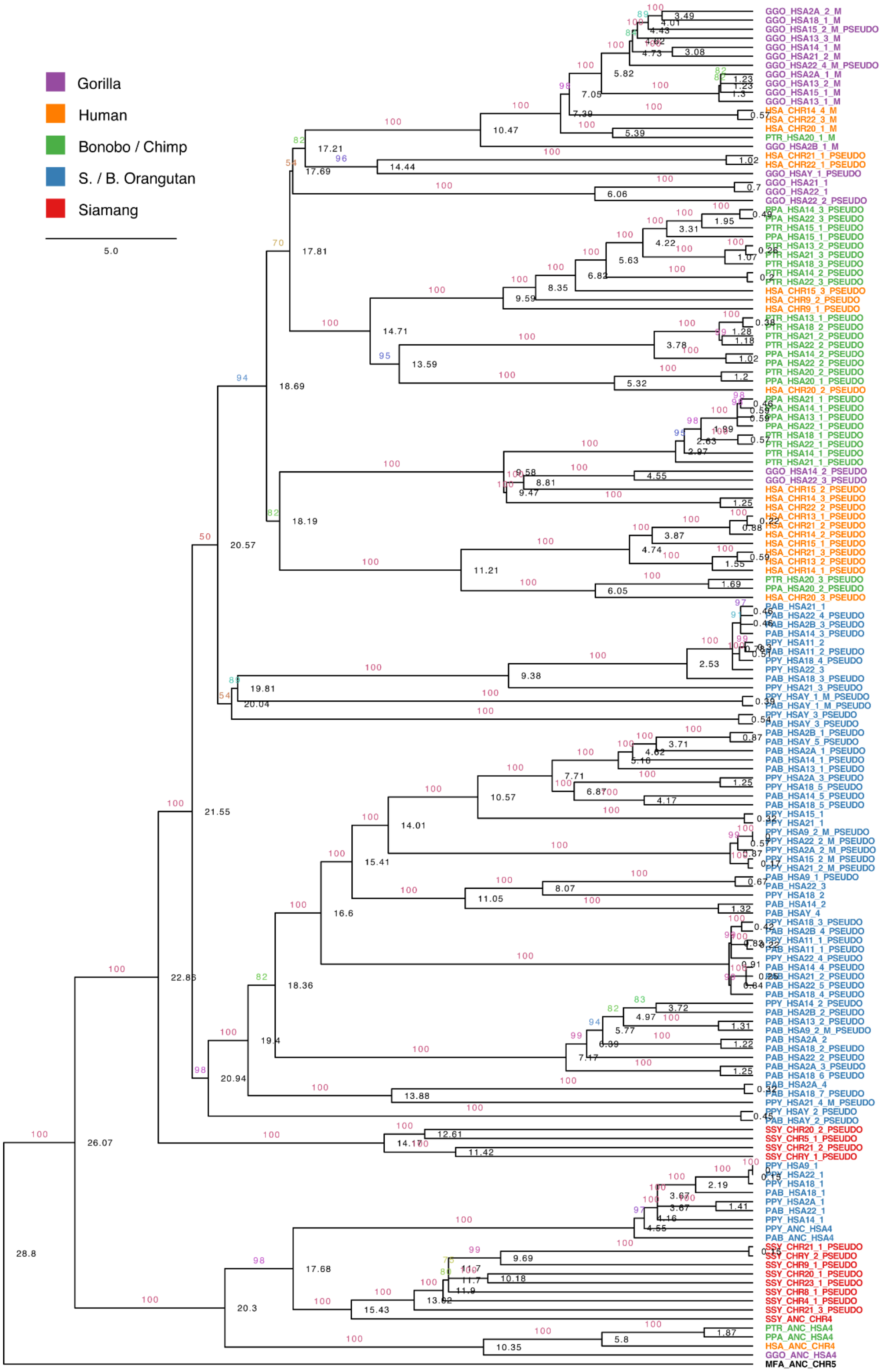

**Figure S41. Time-measured intron phylogeny for *FRG1*.** Tree was rooted with marmoset *FRG1* (branch not shown) and estimated divergence times of 28.8 Ma and 43 Ma were used for the human/macaque and human/marmoset splits, respectively. Leaves are colored by species, branches are labeled by bootstrap, and nodes are labeled by estimated time in Ma.

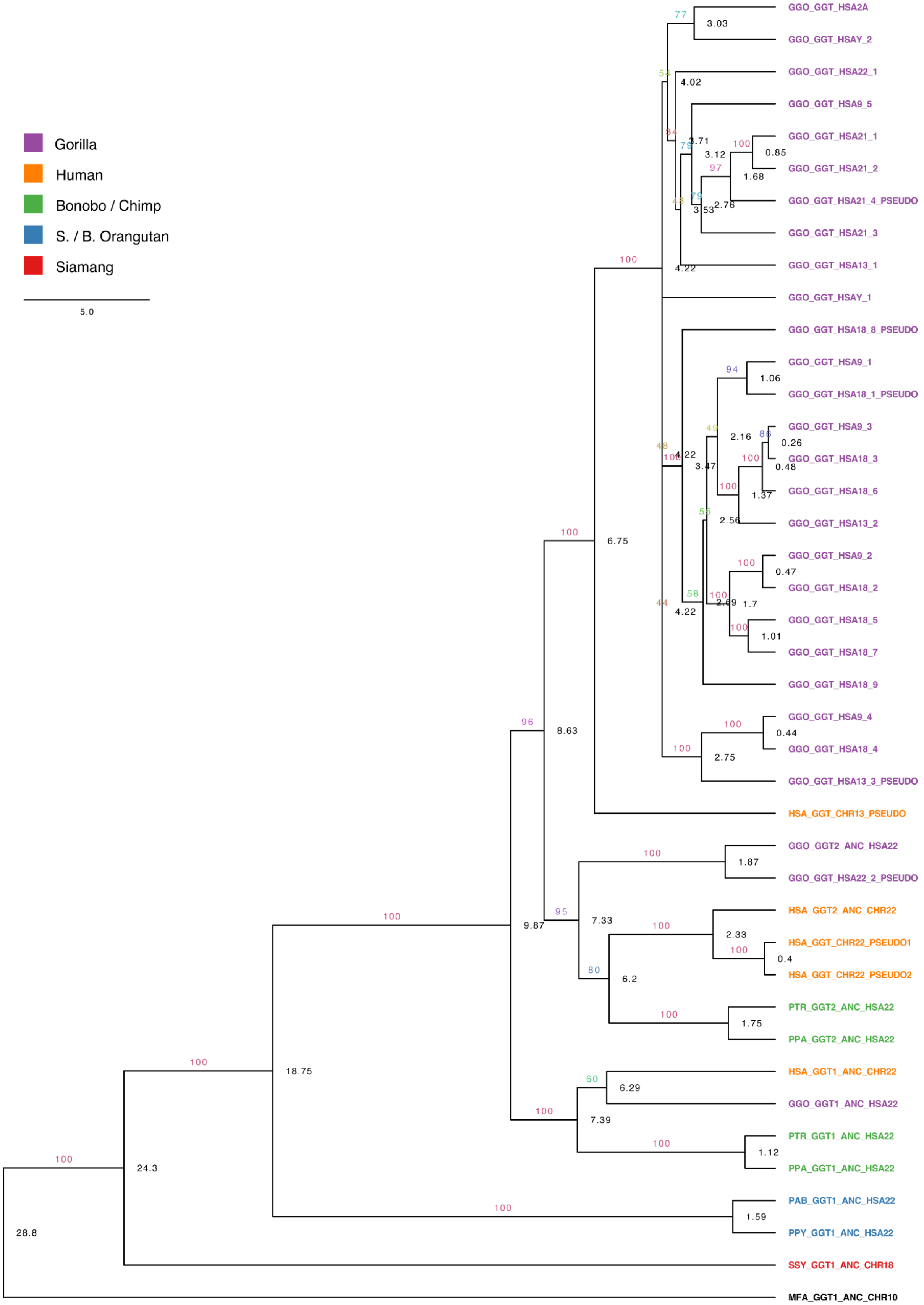
Gorilla
Human
Bonobo / Chimp
S. / B. Orangutan
Siamang
5.0
GGO_GGT_HSA2A
GGO_GGT_HSAY_2
GGO_GGT_HSA22_1
GGO_GGT_HSA9_5
GGO_GGT_HSA21_1
GGO_GGT_HSA21_2
GGO_GGT_HSA21_4_PSEUDO
GGO_GGT_HSA21_3
GGO_GGT_HSA13_1
GGO_GGT_HSAY_1
GGO_GGT_HSA18_8_PSEUDO
GGO_GGT_HSA9_1
GGO_GGT_HSA18_1_PSEUDO
GGO_GGT_HSA9_3
GGO_GGT_HSA18_3
GGO_GGT_HSA18_6
GGO_GGT_HSA13_2
GGO_GGT_HSA9_2
GGO_GGT_HSA18_2
GGO_GGT_HSA18_5
GGO_GGT_HSA18_7
GGO_GGT_HSA18_9
GGO_GGT_HSA9_4
GGO_GGT_HSA18_4
GGO_GGT_HSA13_3_PSEUDO
HSA_GGT_CHR13_PSEUDO
GGO_GGT2_ANC_HSA22
GGO_GGT_HSA22_2_PSEUDO
HSA_GGT2_ANC_CHR22
HSA_GGT_CHR22_PSEUDO1
HSA_GGT_CHR22_PSEUDO2
PTR_GGT2_ANC_HSA22
PPA_GGT2_ANC_HSA22
HSA_GGT1_ANC_CHR22
GGO_GGT1_ANC_HSA22
PTR_GGT1_ANC_HSA22
PPA_GGT1_ANC_HSA22
PAB_GGT1_ANC_HSA22
PPY_GGT1_ANC_HSA22
SSY_GGT1_ANC_CHR18
MFA_GGT1_ANC_CHR10

**Figure S42. Time-measured intron phylogeny for *GGT*.** Tree was rooted with marmoset *GGT1* (branch not shown) and estimated divergence times of 28.8 Ma and 43 Ma were used for the human/macaque and human/marmoset splits, respectively. Leaves are colored by species, branches are labeled by bootstrap, and nodes are labeled by estimated time in Ma. All copies on the long-arm of the HSA22 orthologs apart from the *GGT1* orthologs have a fragment of *IGSF3* upstream. This includes all African great ape *GGT2* copies and the pseudogenized copies of *GGT*-like genes on the long arms of HSA22 found in human and gorilla. This fragment is mostly intronic, containing around half of *IGSF3*'s 50 kb long intron 2 and *IGSF3* exon 3. The copies on the acrocentric short arms of human & gorilla contain intact *IGSF3-GGT* fusions with several *IGSF3* exons maintaining an ORF as described in Fig 6C. However, the copy on human chr13 has premature stop codons and hence is pseudogenized.

**Table S2. Gorilla amplified gene CPMs**
Counts per million (CPM) in gorilla for the acrocentric-amplified gene families. Ancestral (ANC) and duplicated copies are separated where they can be confidently distinguished.
For *LRPAP1*, duplicated transcript structure is unmodified, so may not be accurately distinguished.

| Gene | CPM | CPM in Fibroblasts | CPM in Testes |
|---|---|---|---|
| ABHD17A | 4.464685 | 2.88624 | 16.934763 |
| ABHD17A_ANC | 1.465001 | 1.597862 | 0.413516 |
| BAGE2 | 34.941313 | 0 | 149.612684 |
| C1QTNF3 | 0.290028 | 0.3192 | 0 |
| C1QTNF3_ANC | 51.748689 | 65.835951 | 1.871144 |
| CES4A_ANC | 0.411012 | 0.424353 | 0 |
| FGFR3 | 1.757402 | 0 | 7.552179 |
| FGFR3_ANC | 0.442204 | 0 | 1.900311 |
| FRG1 | 52.86666 | 47.322086 | 74.048347 |
| FRG1_ANC | 16.670041 | 6.159575 | 15.088376 |
| IGSF3-GGT | 6.4645 | 0 | 29.351571 |
| GGT_ANC | 10.570456 | 0 | 45.424989 |
| IGSF3_ANC | 26.713233 | 24.63103 | 50.186948 |
| LETM1_ANC | 28.751217 | 14.845539 | 66.630843 |
| LRPAP1 | 211.055956 | 84.46084 | 589.320736 |
| LRPAP1_ANC | 198.397018 | 67.589349 | 507.22353 |
| SAFB2 | 5.983104 | 0 | 25.716897 |
| SAFB2_ANC | 43.578798 | 27.785882 | 104.79806 |
| TPTE | 4.222014 | 2.556549 | 11.403389 |
| TPTE_ANC | 1.334252 | 0 | 3.987568 |

**Table S3. Gorilla GGT FISH summary**
Summary of FISH results using BAC probe RP11-341D18 (AL356585.7) mapping to the IGSF3–GGT, BAGE locus on metaphase chromosomes from multiple Gorilla gorilla individuals. Signal intensity is reported semi-quantitatively (+ to ++++), and localization is specified when outside the pericentromeric region. Very weak signals are indicated as "weak," differences between haplotypes are shown with "/," and ";" indicates a double signal on the same haplotype. Signals on chromosomes 12, 16, 22, and 23 were consistently detected across all individuals, whereas chromosomes 14 and 16 displayed lower or variable intensity, suggesting potential copy-number polymorphisms.

| | **GGO** | **GGO1** | **GGO2** | **GGO5** | **GGO9** | **GGO14** |
|---|---|---|---|---|---|---|
| **Sex** | female | female | male | female | male | male |
| **chr1 (GGO-HSA1)** | weak p cen | weak p cen | weak p cen | weak p cen | weak p cen | weak +/- |
| **chr12 (GGO-HSA2A)** | ++ p ter; p cen | ++ p ter; p cen | ++ p ter; p cen | ++ p ter; p cen | ++ p ter; p cen | ++ p ter; p cen |
| **chr13 (GGO-HSA9)** | ++++ | ++++ | ++++ | ++++ | ++++ | ++++ |
| **chr14 (GGO-HSA13)** | ++ | + | + | + | + | + |
| **chr16 (GGO-HSA15)** | +/+++ | +/+++ | +/+++ | +/+++ | +/+++ | +/+++ |
| **chr17 (GGO-HSA18)** | ++++ | ++++ | ++++ | ++++ | ++++ | ++++ |
| **chr22 (GGO-HSA21)** | ++++ | +++ | +++ | +++ | +++ | +++ |
| **chr23 (GGO-HSA22)** | ++ q cen; p cen | ++ q cen; p cen | ++ q cen; p cen | ++ q cen; p cen | ++ q cen; p cen | ++ q cen; p cen |
| **chrY** | | | ++ | | + | ++ |

**Table S5. Selection testing**

For each gene, selection was tested on the branch(es) associated with the expansion. For *FRG1*, both the great ape expansion and the "modified" copy expansion were tested. Results are given for two foreground sets: the single leading branch to the expansion, and all branches of the expanded clade ("full exp."). Mean dN/dS with 95% CI calculated with FitMG94, branch length, nonsynonymous (nN) and synonymous (nS) substitution counts estimated with SLAC, number of branches, the p-values for dN/dS ≠ 1, BUSTED (test for episodic diversifying selection), and RELAX (intensified selection pressure), and RELAX K (K < 1 relaxed, K > 1 intensified) are reported. The GARD row gives inferred recombination breakpoints. q-values were calculated using the Benjamini-Hochberg correction.

| | | Gene | ABHD17A | BAGE2 | C1QTNF3 | CES4A |
|---|---|---|---|---|---|---|
| | | **GARD Recombination Breakpoints** | 1 found at position 775 | No evidence | No evidence | 4 found at positions 425, 455, 1358, 1459 |
| | | **foreground** | Gorilla | Gorilla | Gorilla | Gorilla |
| **Leading Branch to the expansion** | | **Mean dN/dS (CI)** | 0.369 (0.113-0.856) | 0.728 (0.225-1.697) | 0.440 (0.073-1.356) | 0.377 (0.101-0.910) |
| | | **Branch Length** | 0.007 | 0.016 | 0.004 | 0.005 |
| | | **nN** | 4 | 4 | 2 | 4 |
| | | **nS** | 2 | 2 | 2 | 4 |
| | **p-val** | **Mean dN/dS ≠ 1** | 0.386 | 0.722 | 0.419 | 0.2 |
| | | **BUSTED** | 0.5 | 0.193 | 0.5 | 0.5 |
| | | **RELAX** | **0.0001** | 0.762 | 0.454 | 0.926 |
| | **q-val (BH)** | **Mean dN/dS ≠ 1** | 0.681 | 0.853 | 0.681 | 0.681 |
| | | **BUSTED** | 0.5 | 0.5 | 0.5 | 0.5 |
| | | **RELAX** | **0.00065** | 0.901 | 0.656 | 0.926 |
| | | **RELAX K** | 6.78 | 0.51 | 1.77 | 0.8 |
| **Full expansion** | | **Mean dN/dS (CI)** | 0.166 (0.098-0.372) | 0.821 (0.550-1.173) | 0.261 (0.065-0.678) | 0.825 (0.600-1.094) |
| | | **Number of branches** | 5 | 42 | 4 | 7 |
| | **p-val** | **BUSTED** | 0.5 | 0.5 | 0.5 | 0.5 |
| | | **RELAX** | 0.673 | **0.0045** | 0.478 | 0.36 |
| | **q-val (BH)** | **BUSTED** | 0.5 | 0.5 | 0.5 | 0.5 |
| | | **RELAX** | 0.673 | **0.0073** | 0.518 | 0.425 |
| | | **RELAX K** | 1.24 | 10.29 | 2.55 | 1 |

| | | **Gene** | **FGFR3** | **FRG1** | | **GGT** |
|---|---|---|---|---|---|---|
| | | **GARD Recombination Breakpoints** | 3 found at positions 37, 380, 1697 | No evidence | | 2 found at positions 312, 778 |
| | | **foreground** | Gorilla | Modified copies | All great apes | Gorilla |
| **Leading Branch to the expansion** | | **Mean dN/dS (CI)** | 0.980 (0.507-1.678) | 1.148 (0.297-2.840) | 0.348 (0.087–0.879) | 0.294 (0.156-0.494) |
| | | **Branch Length** | 0.006 | 0.003 | 0.006 | 0.016 |
| | | **nN** | 11 | 3 | 4 | 14 |
| | | **nS** | 3 | 1 | 3 | 9 |
| | **p-val** | **Mean dN/dS ≠ 1** | 0.949 | 0.5 | 0.206 | **0.0064** |
| | | **BUSTED** | 0.5 | 0.4921 | 0.5 | 0.5 |
| | | **RELAX** | **0.0001** | 0.876 | 0.315 | **0.019** |
| | **q-val (BH)** | **Mean dN/dS ≠ 1** | 0.994 | 0.722 | 0.681 | 0.0832 |
| | | **BUSTED** | 0.5 | 0.5 | 0.5 | 0.5 |
| | | **RELAX** | **0.00065** | 0.926 | 0.585 | 0.0617 |
| | | **RELAX K** | 0.03 | 1.22 | 0.48 | 0.5 |
| **Full expansion** | | **Mean dN/dS (CI)** | 0.786 (0.672-1.381) | 1.053 (0.810-1.345) | 0.740 (0.632–0.858) | 0.987 (0.789-1.237) |
| | | **Number of branches** | 11 | 29 | 47 | 34 |
| | **p-val** | **BUSTED** | 0.5 | 0.4574 | **0.0012** | **0.0072** |
| | | **RELAX** | **0.0001** | 0.133 | **0.0001** | **0.0001** |
| | **q-val (BH)** | **BUSTED** | 0.5 | 0.5 | **0.0156** | **0.0468** |
| | | **RELAX** | **0.00022** | 0.192 | **0.00022** | **0.00022** |
| | | **RELAX K** | 0.02 | 1.26 | 2.67 | 0.17 |

| | | Gene | IGSF3 | LETM1 | LRPAP1 | SAFB2 | TPTE |
|---|---|---|---|---|---|---|---|
| | | **GARD Recombination Breakpoints** | 3 found at positions 711, 984, 1880 | 1 found at position 702 | 3 found at positions 188, 551, 570 | 3 found at positions 245, 299, 527 | 1 found at position 428 |
| | | **foreground** | Gorilla | Gorilla | Gorilla | Gorilla | Gorilla |
| **Leading Branch to the expansion** | | **Mean dN/dS (CI)** | 0.962 (0.616-1.412) | 0.544 (0.211-1.105) | 16.66 (4.17-43.7) | 2.262 (0.898-4.619) | 0.448 (0.026–1.979) |
| | | **Branch Length** | 0.0146 | 0.0063 | 0.003 | 0.009 | 0.036 |
| | | **nN** | 23 | 6 | 3 | 6 | 2 |
| | | **nS** | 9 | 4 | 1 | 1 | 1 |
| | **p-val** | **Mean dN/dS ≠ 1** | 0.994 | 0.379 | 0.24 | 0.396 | 0.578 |
| | | **BUSTED** | 0.5 | 0.5 | 0.475 | 0.364 | 0.5 |
| | | **RELAX** | **0.044** | **0.028** | 0.402 | **0.017** | 0.696 |
| | **q-val (BH)** | **Mean dN/dS ≠ 1** | 0.994 | 0.681 | 0.681 | 0.681 | 0.751 |
| | | **BUSTED** | 0.5 | 0.5 | 0.5 | 0.5 | 0.5 |
| | | **RELAX** | 0.0953 | 0.0728 | 0.653 | 0.0617 | 0.901 |
| | | **RELAX K** | 0.06 | 0.26 | 2.33 | 0 | 3.4 |
| **Full expansion** | | **Mean dN/dS (CI)** | 0.839 (0.714-0.975) | 0.517 (0.291-0.838) | 0.762 (0.466-1.155) | 0.659 (0.424-0.966) | 0.676 (0.440–0.983) |
| | | **Number of branches** | 35 | 7 | 5 | 10 | 35 |
| | **p-val** | **BUSTED** | 0.5 | 0.5 | **0.012** | 0.5 | 0.5 |
| | | **RELAX** | **0.0001** | **0.0009** | **0.0001** | **0.0001** | 0.289 |
| | **q-val (BH)** | **BUSTED** | 0.5 | 0.5 | 0.052 | 0.5 | 0.5 |
| | | **RELAX** | **0.00022** | **0.0017** | **0.00022** | **0.00022** | 0.376 |
| | | **RELAX K** | 6.61 | 0.28 | 3.81 | 0.11 | 6.97 |
